# A cost-free CURE: Using bioinformatics to identify DNA-binding factors at a specific genomic locus

**DOI:** 10.1101/2022.10.21.513244

**Authors:** Casey A. Schmidt, Lauren J. Hodkinson, H. Skye Comstra, Leila E. Rieder

## Abstract

Research experiences provide diverse benefits for undergraduates. Many academic institutions have adopted course-based undergraduate research experiences (CUREs) to improve student access to research opportunities. However, potential instructors of a CURE might still face financial or practical hurdles that prevent implementation. Bioinformatics research offers an alternative that is free, safe, compatible with remote learning, and may be more accessible for students with disabilities. Here, we describe a bioinformatics CURE that leverages publicly available datasets to discover novel proteins that target an instructor-determined genomic locus of interest. We use the free, user-friendly bioinformatics platform Galaxy to map ChIP-seq datasets to a genome, which removes the computing burden from students. Both faculty and students directly benefit from this CURE, as faculty can perform candidate screens and publish CURE results. Students gain not only basic bioinformatics knowledge, but also transferable skills, including scientific communication, database navigation, and primary literature experience. The CURE is flexible and can be expanded to analyze different types of high-throughput data or to investigate different genomic loci in any species.

## INTRODUCTION

Undergraduate research experiences are invaluable to students. Documented benefits include retention in STEM (1), increased confidence in research abilities (2), and inclusion of underrepresented populations (3). Yet many students struggle to find a space in laboratories already at capacity. Course-based undergraduate research experiences, or CUREs, can remedy this problem, as they offer students authentic research experiences within the context of a classroom (4). Not only do CUREs involve many more undergraduates in research than the traditional “apprentice” model, but they also allow faculty (especially those with high teaching loads) to make research progress. For example, the instructor of a CURE course can perform a screen (5, 6), follow up on an interesting result from their lab (7), or increase the rigor and reproducibility of a research project through replication by different lab groups or sections.

Despite these clear benefits, there are often limitations to running bench-based CUREs. For example, large schools with high enrollment might face space and time constraints. In addition, the materials required to perform wet-lab experiments may be expensive and time-consuming to prepare for large classes. Overall, these and other limitations can be prohibitive to implementing CUREs (8).

Bioinformatics CUREs can skirt these hurdles. Because laboratory space is not necessary, the class can be held in a computer lab, a classroom (if the students have access to personal laptops), or completely virtually. There are no costly reagents to purchase or biohazard concerns. Bioinformatics research can offer students with disabilities a less physically demanding alternative to bench-based experiments. It is also compatible with remote or asynchronous teaching, which became necessary during the early COVID-19 pandemic (9, 10).

Although bioinformatics research is typically performed on expensive computing clusters, we instead use Galaxy (11), which is a free, user-friendly platform that integrates many widely-used bioinformatics tools. All memory-intensive computing is performed on Galaxy’s servers, allowing students to simply set up commands, execute, and log off; no sophisticated programming knowledge or computing power is needed. Therefore, bioinformatics research is easily integrated into students’ busy schedules, and each activity can typically be completed in less time than a traditional 3-hour wet laboratory. Students participating in bioinformatics CUREs report high sense of achievement and high levels of satisfaction with their projects (12). Furthermore, students can publish their discoveries, which fosters a sense of belonging to the scientific community (13).

Here, we document a successful CURE that applies bioinformatics tools to discover candidate DNA-binding factors that interact with a genomic locus. Specifically, we investigated the *Drosophila melanogaster* histone gene array, which encodes the replication-dependent histones. Because histones undergo non-canonical mRNA processing and exhibit cell cycle-dependent expression, they require a unique suite of transcription and processing factors (14). Although many of these factors are known, the complete inventory of histone gene expression regulators remains incomplete.

In this CURE, students utilize a hypothesis-based candidate approach to identify existing high-throughput datasets (specifically, ChIP-seq or similar techniques). By mapping the reads from a ChIP-seq experiment to the *Drosophila* histone gene array, students determine if a transcription factor targets the locus, suggesting that it may contribute to histone biogenesis. Our approach functions as a primary screen to identify candidate regulatory proteins and provides opportunities for wet-lab follow-up undergraduate research projects (for example, co-immunostaining for the candidate and a known histone gene regulator to validate bioinformatics findings) (15).

We piloted our CURE remotely with students who were confined at home during the early COVID-19 pandemic. We then transitioned to an in-person experience during a 50-minute weekly “discussion” period attached to a sophomore-level genetics course. Over the course of a semester, each student chose at least one protein to investigate, identified appropriate datasets, aligned datasets to the *Drosophila* histone gene array using Galaxy, and produced alignment figures. The semester culminated in an open poster session, during which the students presented their findings to members of the Biology department (i.e., faculty, staff, and students).

The CURE presented here is beneficial to all parties involved: not only did the students obtain valuable research experience and transferable skills, but they also identified new candidate factors to further investigate in our wet laboratory (L. Hodkinson, personal communication) (15). There are thousands of ChIP-seq datasets across multiple repositories that are available for analysis. Future students could examine other types of high-throughput datasets, such as ATAC-seq, FAIRE-seq, and RNA-seq, to further probe the landscape of the histone gene locus. The bioinformatics analysis presented here can be extended to any annotated locus of interest in any organism. These seemingly endless possibilities support the sustainable implementation and adaptation of this CURE.

### Intended audience

We implemented this CURE in a 200-level genetics course that contained 25 sophomores, juniors, and seniors, most of whom were biology majors. Previously, we piloted the CURE virtually with smaller groups of college students of similar demographics. We also sponsored a remote high school student, indicating that students with a wide range of experience levels can perform the research with appropriate training. In addition, we guided students through the CURE both in-person and virtually, further extending the flexibility of this CURE.

### Learning time

The course had two 75-minute lecture periods and one 50-minute “discussion” period per week over a 14-week semester. Traditionally, the discussion period for this course was used for worksheets, activities, and/or literature discussions. Instead, we ran the CURE during this time. Students completed the majority of the activities in class.

### Prerequisite student knowledge

We covered all of the background information on conceptual topics, such as transcription factors and ChIP-seq, in the lecture portion of the class. Therefore, the only prerequisites for the CURE were the course prerequisites (freshmen-level introductory courses for biology majors). In addition, students did not need prior bioinformatics or computer science experience; all required skills are taught in the training modules.

### Learning objectives

Our overall goal was to provide students with an authentic bioinformatics research experience. Upon completion of this CURE, students will be able to:

1. Search peer-reviewed literature to identify candidate proteins that target the locus of interest.
2. Hypothesize outcomes based on background literature.
3. Identify appropriate datasets.
4. Map datasets to locus of interest.
5. Visualize data by producing alignment figures.
6. Synthesize data and form a conclusion.
7. Propose future directions.
8. Present findings to a wider audience (i.e., peers and department).

## PROCEDURE

### Materials

The following materials are required for this CURE:

- Computer and Internet access
- Galaxy account (free web-based platform, www.usegalaxy.org)
- Integrative Genomics Viewer software (free downloadable software, https://software.broadinstitute.org/software/igv/)
- Learning management software such as Canvas, or cloud storage program such as Google Drive or OneDrive to house files and course materials
- Customizable form software, such as Google forms, to assess weekly student progress. Alternatively, students could use a software such as Benchling, OneNote, or Google Docs as a lab notebook, and allow instructors access to monitor progress
- Poster making software, such as PowerPoint, Google Slides, or Biorender
- Poster printing facility or online poster platform, such as SpatialChat
- Optional: video production software such as Zoom, if the instructor is generating tutorials

### Student instructions

Our class met twice weekly for the lecture portion (75-minute periods, led by course instructor C.S.) and once weekly for the bioinformatics CURE portion (two sections of a 50-minute period, led by graduate teaching assistant L.H.) over 14 weeks (see Appendix 1 for the schedule). During lectures, we followed a “molecules first” rather than “Mendel first” approach (16) to introduce CURE-relevant concepts earlier. For example, concepts covered in the first weeks included transcription, transcriptional regulation, and epigenetics. Lecture topics also paid special attention to high-throughput procedures, such as ChIP-seq and RNA-seq. Students learned how scientists generate sequencing data, how to identify appropriate experimental controls, and the types of research questions that these techniques address. We gauged understanding through formative assessments. For example, one graded assignment was to draw the steps of ChIP-seq (Figure 1). This approach syncronized the lecture and discussion sessions and provided students with the required background knowledge for the CURE.

**Figure 1:**
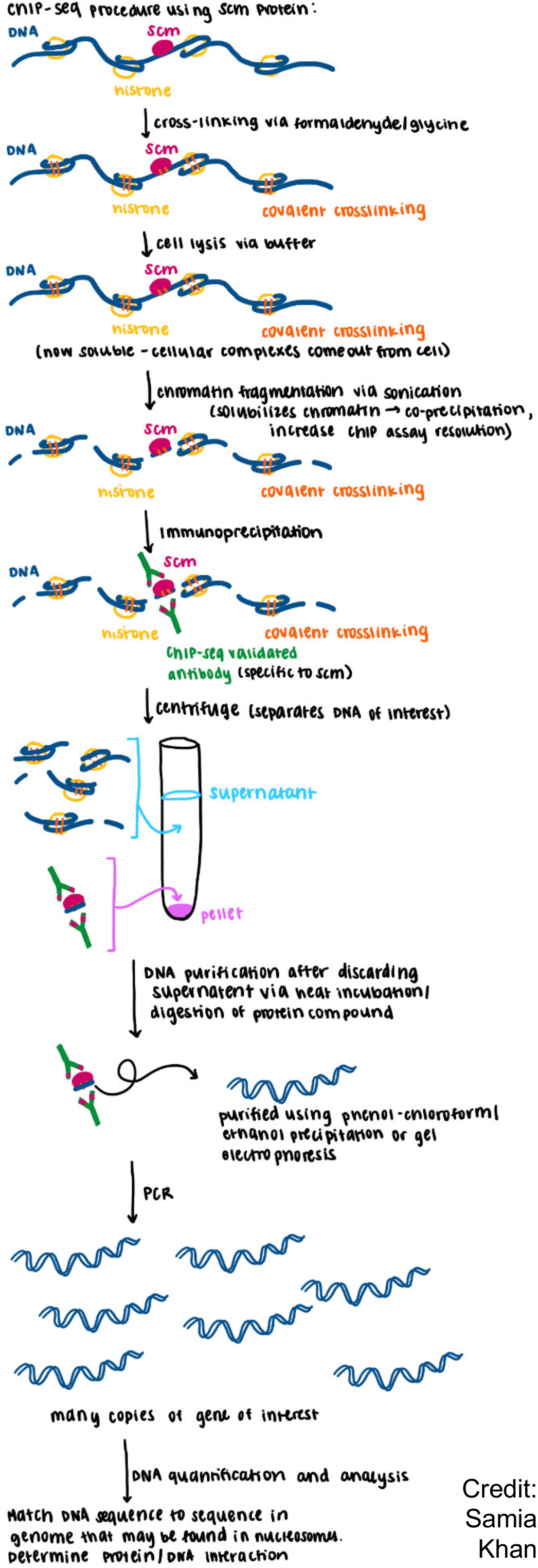
A formative assessment wherein students drew the process of ChIP-seq, using their selected candidate protein. We instructed students to create their own graphics (i.e. no screenshots from the book or other figures). This student chose to investigate Sex comb on midleg (Scm), because it interacts with a factor known to influence histone gene expression (Multi sex combs; Mxc). This assessment linked the CURE and lecture sessions, evaluated student understanding of the ChIP-seq process, and allowed the instructors to address any student misconceptions.

During the discussion period, we spent the first four weeks introducing students to *Drosophila* histone gene expression through literature discussions. Students read and discussed both a review article (14) and a research article that used a bioinformatics approach similar to that introduced in the CURE (17). For each paper, we assigned small groups a figure to annotate and present to the class (see appendix 2).

Students began the process of selecting their candidate protein(s) in the fourth week, with several guiding criteria, such as: (A) proteins that interact with known histone regulators, using protein interaction databases such as STRING (https://string-db.org/) (18); (B) transcription factors that act in the early *Drosophila* embryo, which requires rapid histone biosynthesis (19); (C) DNA-binding factors implicated in cell cycle progression, as histone expression is linked to S-phase (14); and (D) dosage compensation factors, because a prominent histone gene regulator is also involved in dosage compensation (17). Students also used PubMed and FlyBase (20) to gather information. Importantly, the candidate selection was student-driven with input from the instructors.

We followed these background and brainstorming sessions with three weeks of bioinformatics tutorials, during which we led students through analysis and visualization of example data using Galaxy (11) and Integrative Genomics Viewer software (IGV; (21)). We used ChIP-seq data from the background primary research article (17) to ensure that their results matched the published figures. See appendix 3 for the ChIP-seq analysis workflow. Due to computing demands on the Galaxy servers, some tools can take several hours to complete. During any downtime, students continued their background research on candidate proteins. We consulted with each student individually to provide guidance.

After these tutorials, the next six discussion periods functioned as work sessions for students to carry out their bioinformatics analyses. Because the majority of NIH-funded high-throughput sequencing experiments are deposited into public databases such as the NCBI Gene Expression Omnibus (GEO; https://www.ncbi.nlm.nih.gov/geo/), many students identified ChIP-seq datasets for their candidate protein(s) by directly searching this database. Others located accession numbers within primary literature. Some students formed strong hypotheses but could not locate appropriate ChIP-seq datasets; we guided these students to additional databases such as modENCODE (22), which contains ChIP-seq datasets for many *Drosophila* transcription factors. Once each student located usable data, they aligned the ChIP-seq reads to the *Drosophila* genome using Bowtie2 (23) in Galaxy and subsequently generated alignment figures using IGV (appendix 3). We consulted with each student individually to discuss their conclusions and next steps. Several students had time to investigate another (often related) candidate based on the conclusions from their first experiment.

We dedicated two of the work sessions to poster design. We presented examples of well-crafted posters, provided a template, shared our assessment rubric (appendix 4), and provided feedback on drafts before posters were printed. On the last day of the discussion period, the students presented their posters to their peers and a wider audience (i.e., Biology department faculty, staff, and students). In addition to presenting the poster, each student filled out three peer review forms for other students (appendix 5).

### Faculty instructions

We used this CURE to investigate protein factors that target the *Drosophila* histone gene array. Below, we provide options for customization.

## Designing the research project

Instructors can adapt this CURE to study any genomic locus (for example, an enhancer region that might attract transcription or other regulatory factors). The workflow is particularly suitable for repetitive regions (such as the histone gene array) because they are often excluded from genome-wide analyses. Galaxy contains many built-in genomes, but instructors can also provide a custom genome. We used a custom genome that contains a single copy of the histone gene array (24) because the sequences of the ~100 array copies are nearly identical in the *Drosophila melanogaster* genome (25). This approach also amplifies ChIP-seq signal (17).

## Ensuring availability of appropriate data

Before beginning this CURE, the instructor should browse for ChIP-seq datasets to familiarize themselves with the databases. An excellent starting point is the NCBI GEO, an NIH-sponsored public repository that houses next-generation sequencing data. Other databases include the European Nucleotide Archive (https://www.ebi.ac.uk/ena/browser/home), the ENCODE project (26), as well as modENCODE (22) for model organism data. These projects contain many ChIP-seq datasets. Instructors may also wish to have a set of backup datasets for students who have difficulty locating appropriate data.

Students can also map data from ChIP-seq variation techniques, such as ChIP-nexus (27), CUT&RUN (28), and CUT&TAG (29) using the same bioinformatic analysis as ChIP-seq. However, ChIP-chip (chromatin immunoprecipitation followed by microarray) datasets cannot be used with the bioinformatics tools in this CURE, because microarrays utilize different analyses. Unfortunately, some datasets do not contain appropriate controls. For example, we routinely find ChIP-seq datasets that do not include an input or control immunoprecipitation (e.g. IgG) condition, which are important to normalize the experimental ChIP data. The lack of normalization can sometimes lead to misleading or false positive results, wherein small local peaks appear as positive signal (for example, see Figure 3B) (15). Although there is no way to rectify the lack of controls, it allows for important discussions with students on what conclusions one can draw from their datasets.

## Preparing background material

We recommend covering basic conceptual aspects of the project during lecture if students are not already familiar. We also suggest devoting a few early CURE sessions to reading primary and/or secondary literature. This approach introduces the genomic locus of interest, allows students to gain experience in navigating scientific literature, and reinforces the idea that there are still many “unknowns” in scientific research.

## Work days

Instructors should test the Galaxy tools a few weeks before teaching students to ensure that the workflow is functional and to gain familiarity with the user interface. The instructor may also wish to generate their own tutorials if they modify the analysis pipeline. We recommend recording video tutorials through Zoom, which has screen-sharing capability.

## Preparing data for publication

This CURE functions as a primary screen to identify candidate regulators of a genomic locus. If instructors wish to publish their results, we recommend verification of the students’ bioinformatics findings. We employed one of our CURE students for this task and compiled the validated results from the class into a manuscript, with all participating students as co-authors (L. Hodkinson, personal communication).

### Suggestions for determining student learning

Student posters were the primary mode of assessment for our CURE (worth 25% of the discussion grade, plus 15% for the poster peer review assignment). The remaining 60% of the discussion grade was based on participation in the research, assessed through student-reported activity logs. It is sometimes difficult to assess inquiry-based research, and the bioinformatics component added additional hurdles for some students. For example, there may not exist appropriate datasets for a student’s selected candidate, Galaxy may perform slowly, or a dataset from a large study may contain many variables (e.g., environmental conditions, mutant genotypes, treatments, tissue types) such that students struggle to determine which samples are relevant (see appendix 3). Therefore, we emphasized progress and effort over results. At the end of each discussion session, the students filled out a Google form describing the activities they performed that day. These forms included space to upload a screenshot of Galaxy or IGV (Figure 2). Through the Google forms, we assessed participation and monitored progress so that we could intervene if necessary.

**Figure 2:**
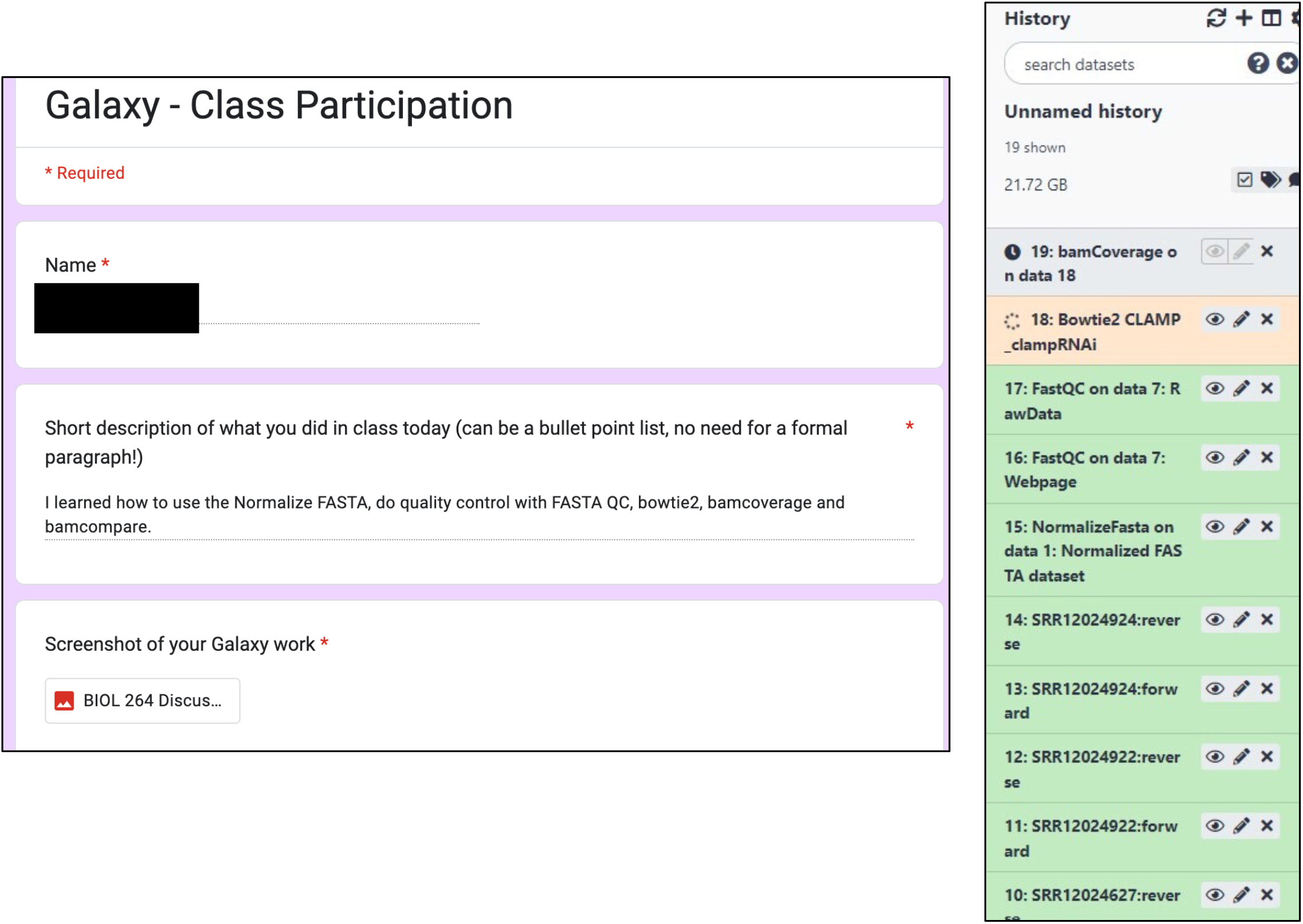
Example of a student’s Google form submission. This form was submitted during Week 6 (Appendix 1) when the students were completing Galaxy tutorials on practice data (Rieder 2017). The student’s screenshot (right) shows their Galaxy history, with several jobs complete (green), one in progress (orange), and one in the queue (gray).

An additional approach to determining student learning is to include formative assessments throughout the semester. For example, groups of students might complete a worksheet such as the Figure Facts template (30) that walks through a figure from a primary research article. Students could also gain presentation experience by sharing a research article that includes the dataset they plan to analyze. The instructor may choose to have students report their activities in graded lab notebooks. These assessments offer additional opportunities for instructor feedback but may be impractical in a larger class.

### Sample data

Students mapped all relevant conditions, including controls, ChIP, and normalization (if applicable), for their candidate(s), and visualized alignments using IGV software (21). From there, students formed conclusions about their candidate targeting the histone gene array. Below, we present “positive” (Figure 3A) and “negative” (Figure 3B) candidates, both from our remote summer high school student. We are also compiling the results from other iterations for publication (L. Hodkinson, personal communication) (15).

**Figure 3:**
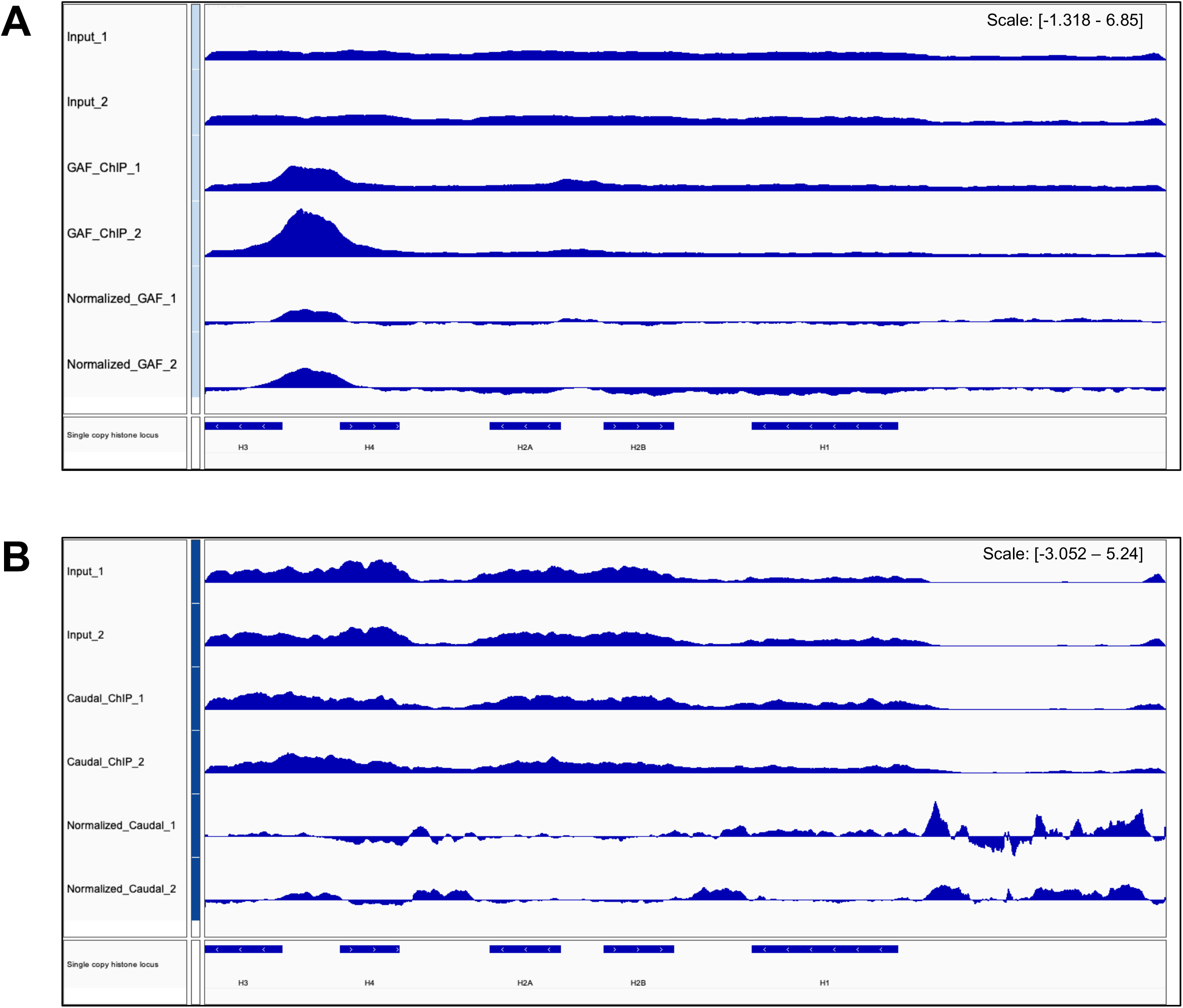
Sample data. (A) ChIP-seq alignment of GAGA Factor (GAF) in stage 3 *Drosophila* embryos. The figure shows two replicates from the same study. There is a clear peak between the H3 and H4 genes, suggesting that GAF localizes to this region. This finding was surprising, given that GAF does not target the histone gene array in cultured S2 cells or by immunofluorescence in early embryos (Rieder 2017). Data from (Gaskill eLife 2021). (B) ChIP-seq alignment of Caudal in 0-4hr embryos. The figure shows two replicates from the same study. Although there is a signal upstream of H1 in the normalized panels, the peaks are not reflected in the ChIP panels, suggesting that they are not true signal. Thus, there is no clear enrichment of Caudal at the histone gene array. Data from (modENCODE).

### Safety issues

Because this activity does not involve a traditional laboratory setup, we do not foresee any safety issues.

## DISCUSSION

### Field testing

We began this bioinformatics project as a strategy to engage our junior laboratory members in remote work during the early COVID-19 pandemic. During the fall of 2020, undergraduates at our institution were not permitted to work in research buildings. Instead, our undergraduate laboratory researchers collectively learned basic bioinformatics skills (led by H.S.C.). Four students each chose a protein to study, identified datasets, mapped data to the histone gene array, and presented their findings to the larger laboratory group. After this first pilot, we recruited nine naive undergraduates to remotely study the chromatin landscape of the *Drosophila* histone gene array in the spring of 2021. For this iteration, students chose a histone post-translational modification and mapped ChIP-seq data from the modENCODE project (22). The students presented their findings to a wider audience via a virtual poster session.

Our laboratory also sponsored a remote high school student that continued the bioinformatics project during the summer of 2021 (Figure 3). This student investigated several early *Drosophila* embryo patterning factors, providing our wet laboratory with candidates for follow-up studies. Most recently (spring 2022), we implemented the project as a CURE in a 200-level genetics course with 25 students, as described in the “procedure” section of this manuscript.

The class size will likely contribute to the effectiveness of this CURE. Our weekly discussion period was split into two 50-minute sections, with 14 students in one and 11 in the other. This small size allowed us the opportunity to grant individual attention to each student. Because several of our students ran into difficulties finding appropriate ChIP-seq datasets for their chosen candidate factor, we found that this 1:1 time was necessary to ensure the success of all students, and we recommend a ratio of one instructor to no more than 15 students. If individual meetings are not feasible, the instructor could employ additional experienced TAs to consult with the students, or students could operate in small groups.

### Evidence of student learning

We primarily evaluated the learning objectives through the student posters, which served as a summative assessment. Learning objectives 1-7 were reflected in the poster rubric (Figure 4). The posters were worth 50 points in total. We allowed students to receive instructor feedback on the poster before the final submission. Student grades for the poster ranged between 80-100%. The majority of deductions were related to data presentation, as we instructed students to change the default labels and font size in the IGV plots (Appendix 4).

**Figure 4:**
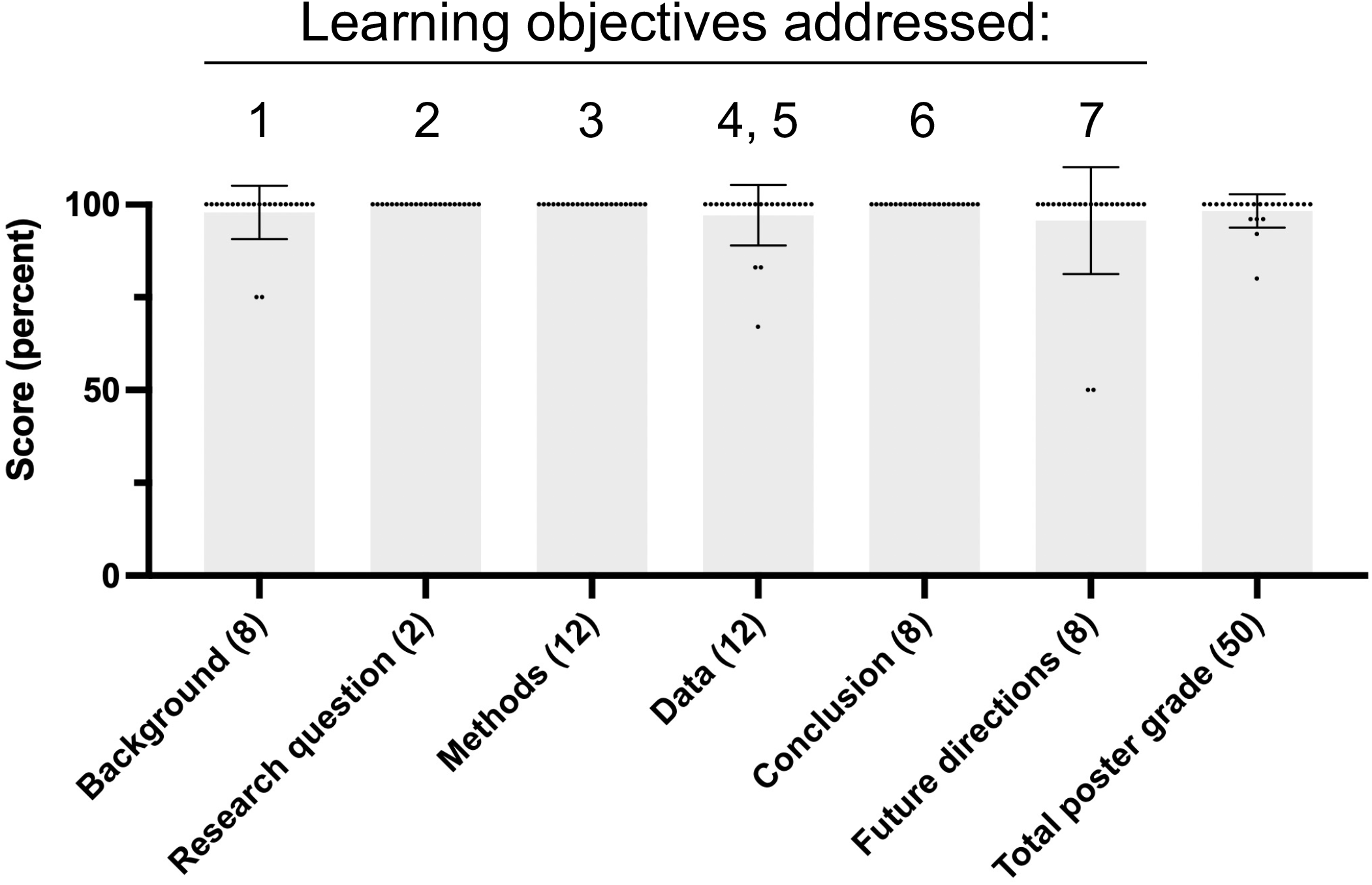
Student posters were the primary form of summative assessment for this CURE. Bar graph representing score (as a percent) for individual poster sections and the entire poster. Each dot represents an individual student. The point value of each poster section is listed in parentheses. Error bars denote SEM. Learning objectives addressed by each poster section are listed above the bars.

We also documented student learning in CURE-related exam questions, which were answered correctly by at least 80% of students (Figure 5). For example, we asked what experiment a student would perform to determine the genomic localization of a hypothetical new histone variant protein. This question, which we classify in the “Apply” level of Bloom’s taxonomy (31), required students to recall that histones are DNA binding proteins and to differentiate between types of high-throughput experiments (Figure 5A). In addition, we asked students to draw the results of a ChIP-seq experiment if the researcher forgot to add the antibody (Figure 5B). We classify this question in the “Analyze” level of Bloom’s taxonomy, because it addresses the role of different reagents in an experiment. Collectively, these results demonstrate that our students displayed higher-order reasoning on CURE-related topics in their exams.

**Figure 5:**
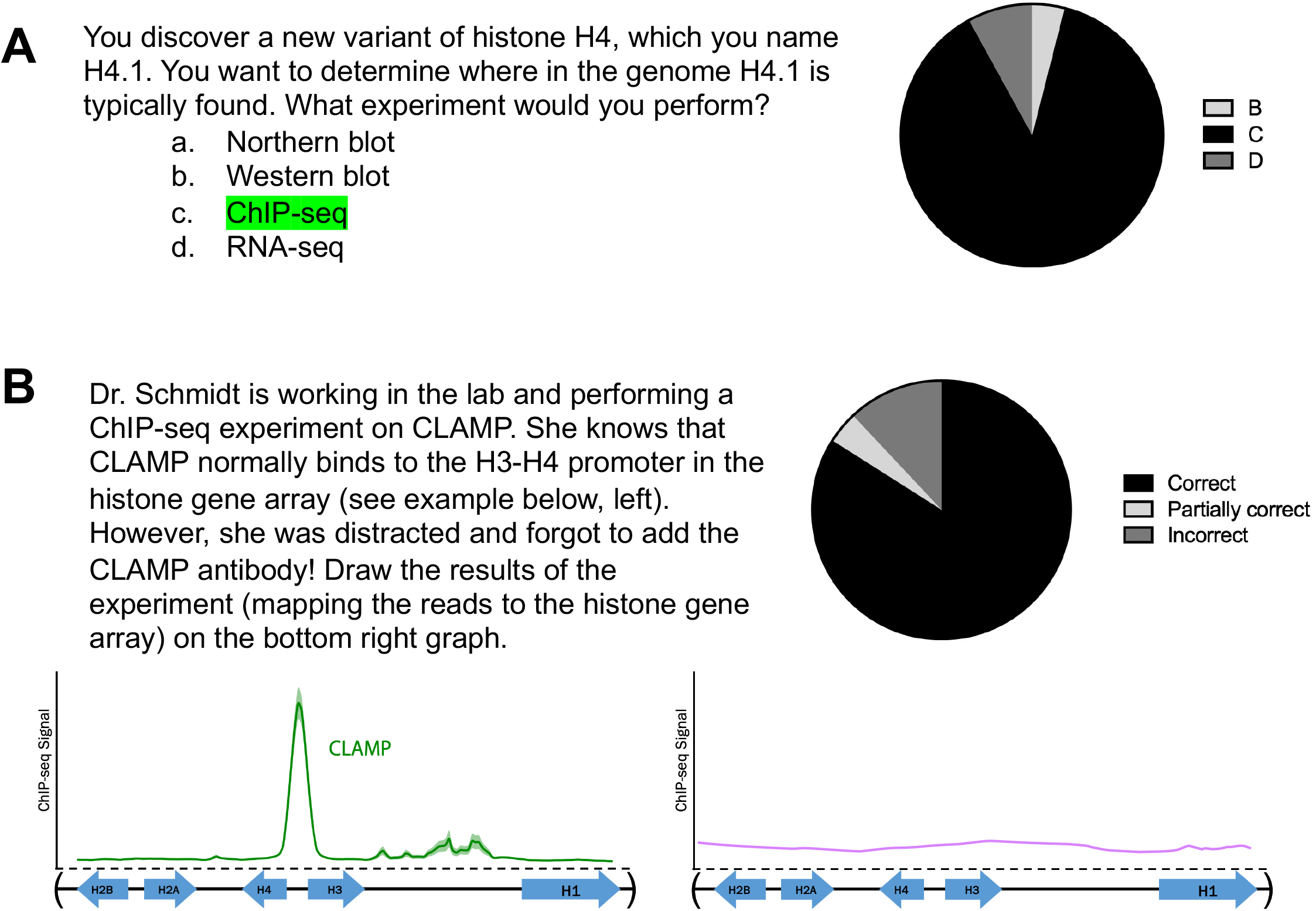
CURE-related exam questions. (A) Multiple choice question answered correctly by 88% of students; the correct answer is highlighted in green. (B) Open-ended question answered correctly by 84% of students; a correct answer is drawn on the right panel by the instructor in purple. CLAMP data from (Rieder 2017).

### Possible modifications

There are many ways instructors could modify this CURE. As presented in the “Faculty instructions” section of this manuscript, students can apply the bioinformatics analysis pipeline to any locus of interest in any species. For example, they could search for factors that target regions near a known gene; this approach might identify unannotated enhancers. Students could also examine chromatin landscape data, such as ATAC-seq or FAIRE-seq, and compare to histone modification ChIP-seq datasets that correlate with different chromatin states (32). In addition, students can map RNA-seq datasets if the locus of interest includes genes; however, the treatment of the data is different from ChIP-seq and will require additional tutorials (for example, https://qubeshub.org/publications/118/4).

An exciting follow-up to the bioinformatics CURE is to confirm positive candidates with wet lab experiments. *Drosophila melanogaster* is a particularly useful model organism for these follow-up studies due to the wealth of available mutant and RNAi lines in public stock centers, as well as established protocols for staining tissues such as polytene chromosomes (15), wing discs, and embryos. There are also numerous custom antibodies that researchers can request from individual laboratories or purchase from stock centers such as the Developmental Studies Hybridoma Bank (https://dshb.biology.uiowa.edu/). These wet-lab experiments can provide a platform for future studies: for example, testing histone gene expression in the absence of a validated protein that targets the histone gene locus (17).

The data generated from this CURE will ultimately add to the growing body of knowledge regarding transcription factor targeting of genomic loci. In addition, the CURE provides students with an authentic research experience, especially in situations where in-person wet laboratory research is not feasible. Students also gain transferable skills that are important for STEM education, including: (A) reading and interpreting primary literature; (B) forming hypotheses based on prior research; (C) navigating complex databases; (D) drawing conclusions from data; and (E) proposing future studies. Furthermore, students interested in continuing bioinformatics research will require less training because they have learned basic bioinformatics techniques. The skills gained during this CURE are crucial to both research science and critical thinking.

## Supporting information

Tutorial

## ACKNOWLEDGEMENTS

We are grateful to the Emory University students who participated in our first (Dabin Cho, Greg Kimmerer, Mary Wang, and Mellisa Xie) and second (Eric Albanese, Yono Bulis, Edgar Hsieh, Shaariq Khan, Andre Mijacika, Sean Parker, Rohan Ramdeholl, Annalise Weber, and Kelly Yoon) remote pilots, as well as the 25 BIOL 264 students who participated in the CURE. We also thank Karen Resendes, Kelsey Gray, Jennifer Gresham, Ethan Rundell, and Michaelyn Hartmann for critically reviewing this manuscript. This work was supported by K12GM00068 to CAS and HSC; F32GM140778 to CAS; T32GM00008490 and F31HD105452 to LJH; and R00HD092625 and R35GM142724 to LER.

## Appendix 1

**Appendix 1:**
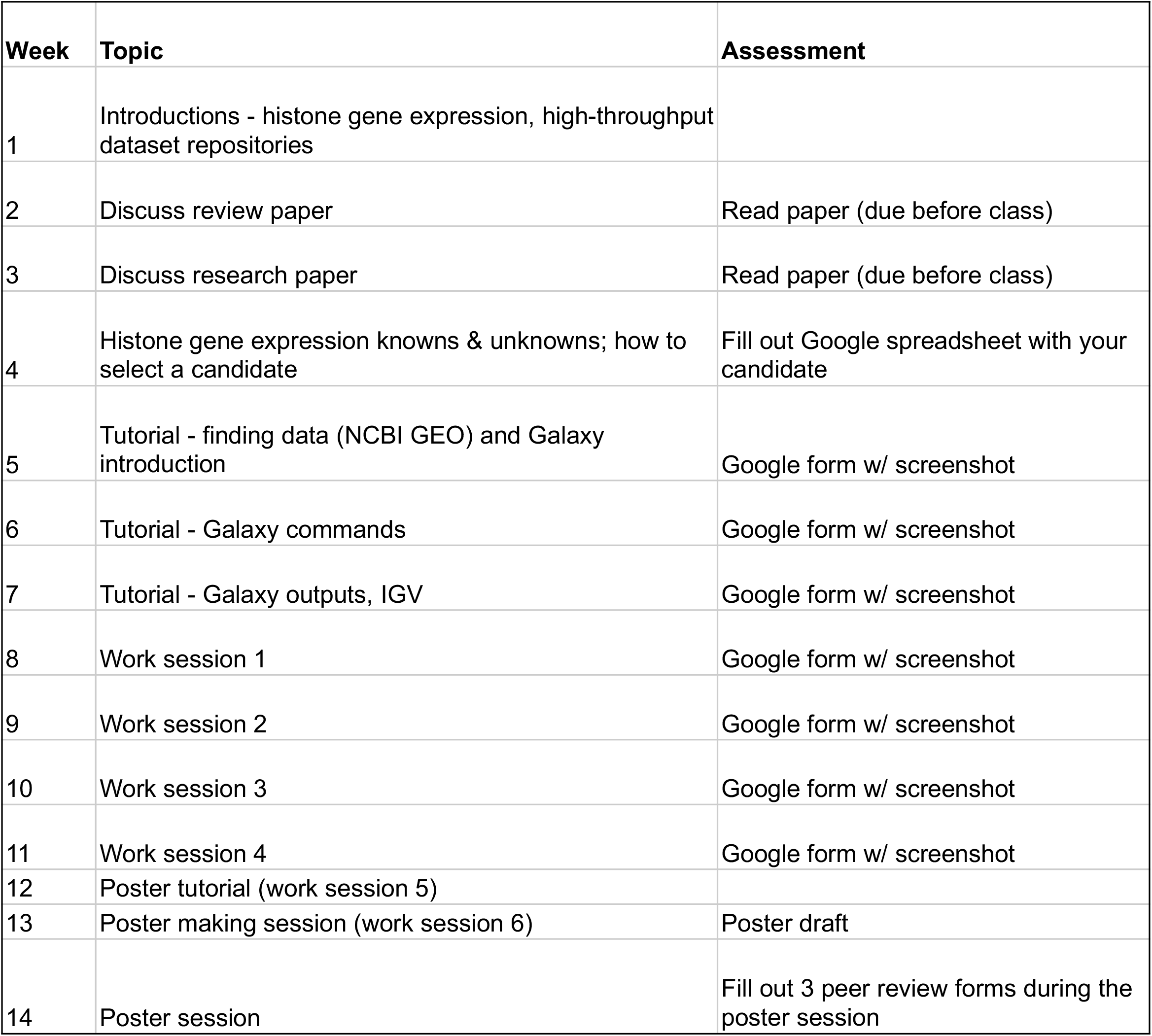
Weekly class schedule for the CURE

| Week | Topic | Assessment |
| --- | --- | --- |
| 1 | Introductions - histone gene expression, high-throughput dataset repositories |  |
| 2 | Discuss review paper | Read paper (due before class) |
| 3 | Discuss research paper | Read paper (due before class) |
| 4 | Histone gene expression knowns & unknowns; how to select a candidate | Fill out Google spreadsheet with your candidate |
| 5 | Tutorial - finding data (NCBI GEO) and Galaxy introduction | Google form w/ screenshot |
| 6 | Tutorial - Galaxy commands | Google form w/ screenshot |
| 7 | Tutorial - Galaxy outputs, IGV | Google form w/ screenshot |
| 8 | Work session 1 | Google form w/ screenshot |
| 9 | Work session 2 | Google form w/ screenshot |
| 10 | Work session 3 | Google form w/ screenshot |
| 11 | Work session 4 | Google form w/ screenshot |
| 12 | Poster tutorial (work session 5) |  |
| 13 | Poster making session (work session 6) | Poster draft |
| 14 | Poster session | Fill out 3 peer review forms during the poster session |

## Appendix 2

**Appendix 2:**
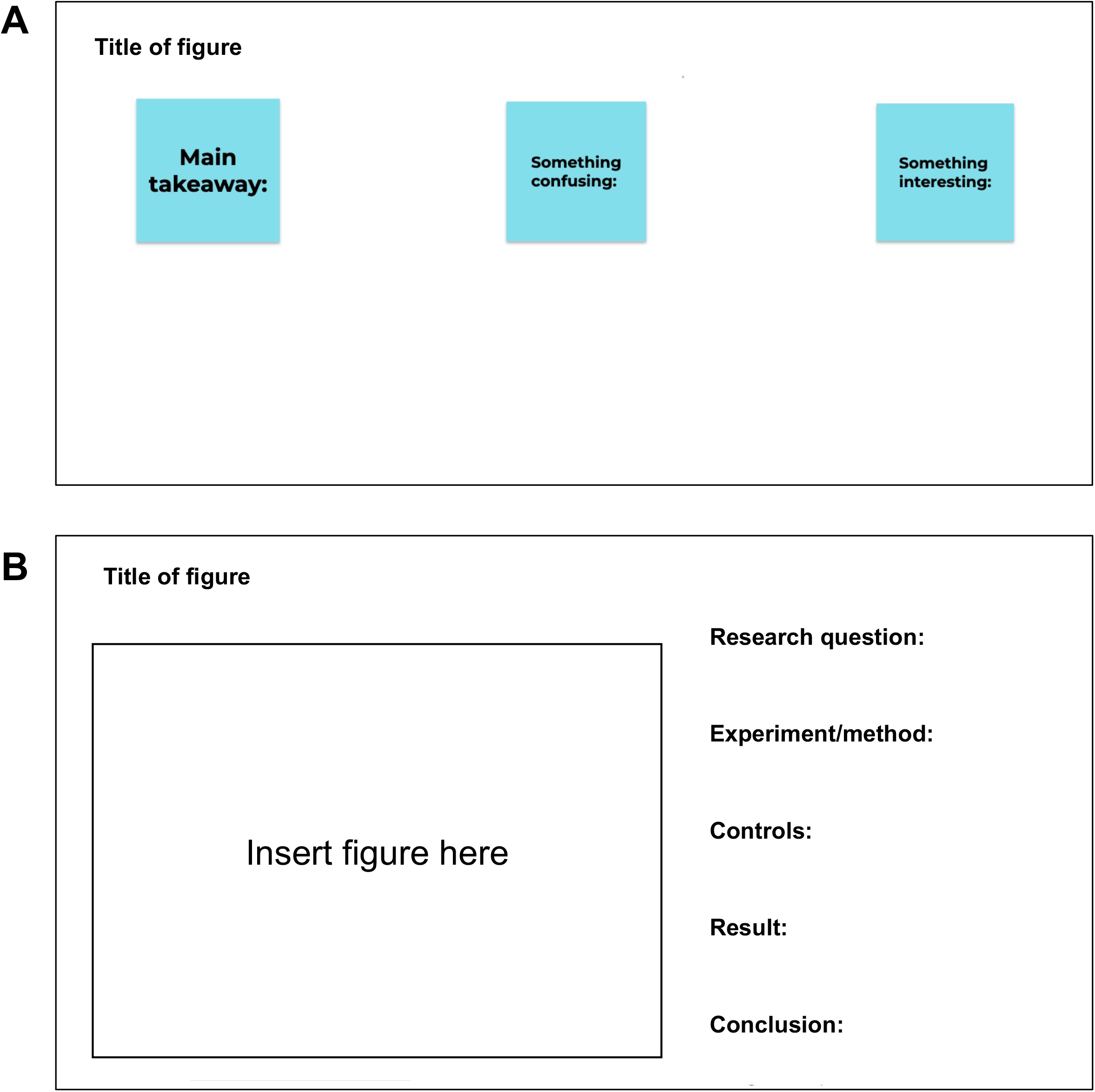
Template for student annotation of review and research articles, via Jamboard (A) and Google Slides (B). (A) Students responded to the the prompts by virtual “sticky notes” on Jamboard (https://jamboard.google.com/). (B) Students responded to the prompts by creating text boxes and typing in answers on Google Slides.

## Appendix 3

Tutorials (separate document)

## Appendix 4

**Appendix 4:** Poster rubric given to students

|  | Satisfactory | Unsatisfactory |
| --- | --- | --- |
| Background (8 points) | -Includes 1 figure drawn in BioRender<br>-Includes prior knowledge on your specific factor<br>-Includes background on the *Drosophila* histone gene locus<br>-Includes citations for background information | -Does not include 1 figure drawn in BioRender<br>-Does not include prior knowledge on your specific factor<br>-Does not include background on the *Drosophila* histone gene locus<br>-Does not include citations for background information |
| Research question (2 points) | -Accurately summarizes your project/data | -Does not accurately summarize your project/data |
| Methods (12 points) | -Cites paper that data came from<br>-Lists all Galaxy tools used and their purpose | -Does not cite paper that data came from<br>-Does not list all Galaxy tools used and their purpose |
| Data (12 points) | -Changes default font to larger size<br>-Comparisons have same scale<br>-IGV tracks are re-labeled to short, descriptive titles | -Font not changed from default<br>-Comparisons have different scales<br>-IGV tracks are not re-labeled |
| Conclusion (8 points) | -Conclusion is appropriate for the data presented and addresses limitations (i.e. cell type, time point, etc.) | -Conclusion is not appropriate for the data presented or does not address limitations |
| Future directions (8 points) | -Includes at least 2 future directions | -Does not include at least 2 future directions |

## Appendix 5

**Appendix 5:** Peer review form that students completed during the poster session. Each student filled out 3 forms, which counted toward their own grade.

|  |
| --- |
| Your name: _____ |
| Name of poster presenter: _____ |
| What factor or factors are being studied? _____ |
| What is the main finding? _____ |
| _____ |
| _____ |
| What future directions are proposed? _____ |
| _____ |
| _____ |
| What is one thing you liked about the poster? _____ |
| _____ |
| _____ |

