## Supplementary material for "A cost-free CURE: Using bioinformatics to identify DNA-binding factors at a specific genomic locus": Tutorial

### Galaxy and IGV tutorial

To use this tutorial: create a Galaxy account and log in (usegalaxy.org)

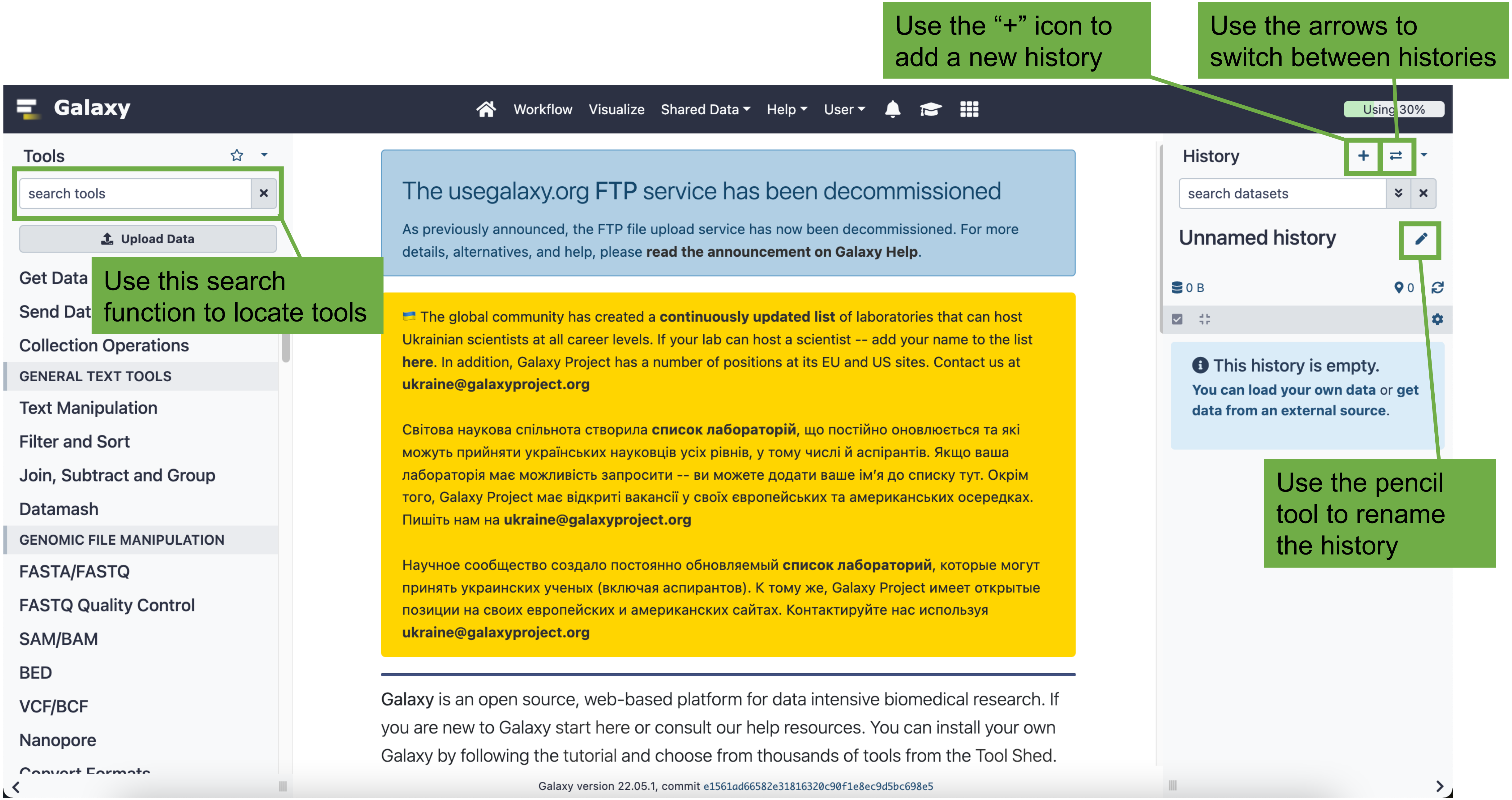
Use the "+" icon to add a new history
Use the arrows to switch between histories
Galaxy
Workflow
Visualize
Shared Data
Help
User
Using 30%
Tools
search tools
Upload Data
Use this search function to locate tools
Get Data
Send Dat
Collection Operations
GENERAL TEXT TOOLS
Text Manipulation
Filter and Sort
Join, Subtract and Group
Datamash
GENOMIC FILE MANIPULATION
FASTA/FASTQ
FASTQ Quality Control
SAM/BAM
BED
VCF/BCF
Nanopore
The usegalaxy.org FTP service has been decommissioned
As previously announced, the FTP file upload service has now been decommissioned. For more details, alternatives, and help, please read the announcement on Galaxy Help.
The global community has created a continuously updated list of laboratories that can host Ukrainian scientists at all career levels. If your lab can host a scientist -- add your name to the list here. In addition, Galaxy Project has a number of positions at its EU and US sites. Contact us at
Світова наукова спільнота створила список лабораторій, що постійно оновлюється та які можуть прийняти українських науковців усіх рівнів, у тому числі й аспірантів. Якщо ваша лабораторія має можливість запросити -- ви можете додати ваше ім'я до списку тут. Окрім того, Galaxy Project має відкриті вакансії у своїх європейських та американських осередках. Пишіть нам на
Научное сообщество создало постоянно обновляемый список лабораторий, которые могут принять украинских ученых (включая аспирантов). К тому же, Galaxy Project имеет открытые позиции на своих европейских и американских сайтах. Контактируйте нас используя
Galaxy is an open source, web-based platform for data intensive biomedical research. If you are new to Galaxy start here or consult our help resources. You can install your own Galaxy by following the tutorial and choose from thousands of tools from the Tool Shed.
Galaxy version 22.05.1, commit e1561ad66582e31816320c90f1e8ec9d5bc698e5
History
search datasets
Unnamed history
0 B
0
This history is empty.
You can load your own data or get data from an external source.
Use the pencil tool to rename the history

### Getting data

#### Option 1: manual upload

Use this option if files are stored locally

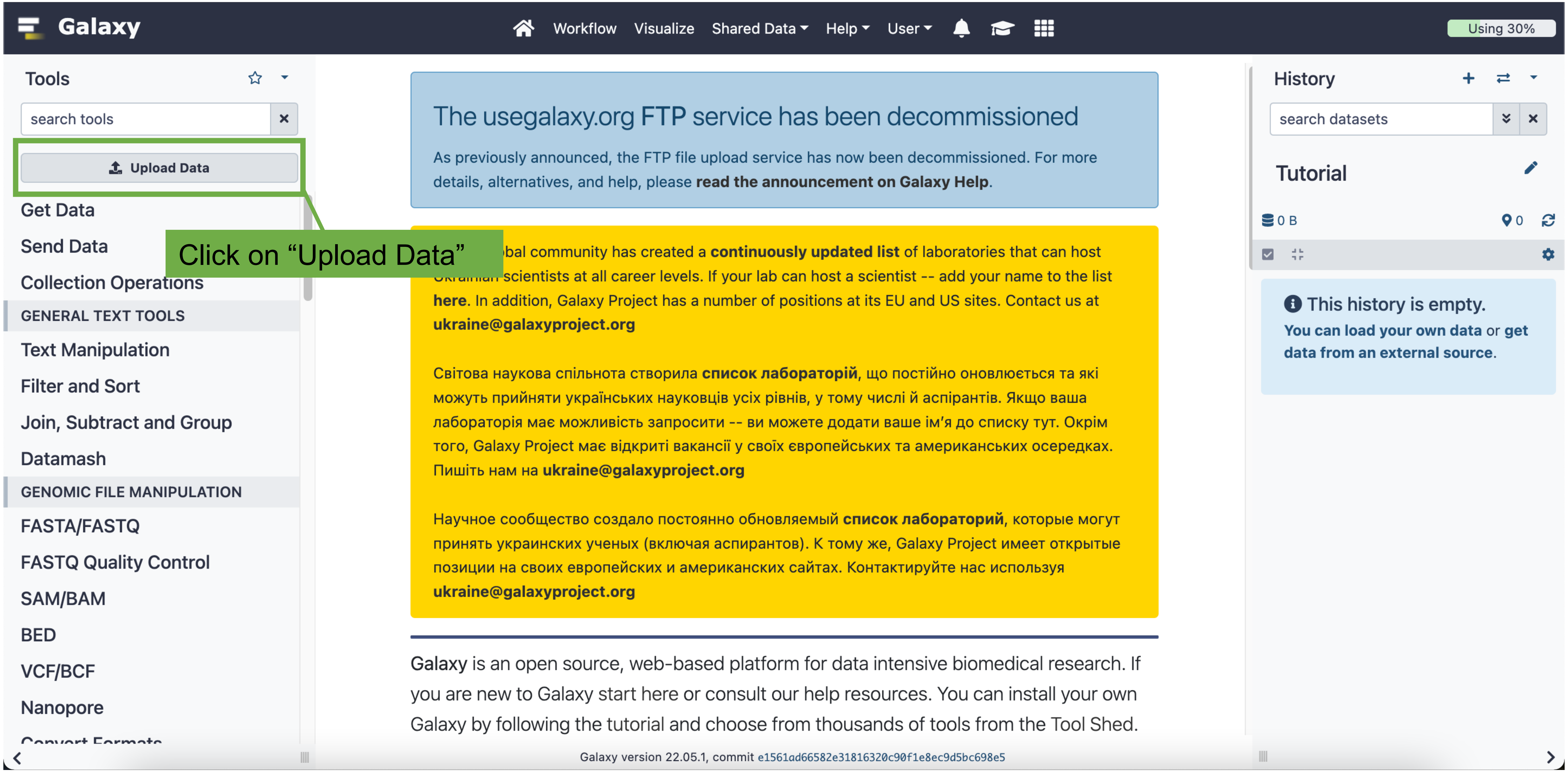
Galaxy
Workflow
Visualize
Shared Data
Help
User
Using 30%
Tools
search tools
Upload Data
Get Data
Send Data
Collection Operations
GENERAL TEXT TOOLS
Text Manipulation
Filter and Sort
Join, Subtract and Group
Datamash
GENOMIC FILE MANIPULATION
FASTA/FASTQ
FASTQ Quality Control
SAM/BAM
BED
VCF/BCF
Nanopore
Click on “Upload Data”
The usegalaxy.org FTP service has been decommissioned
As previously announced, the FTP file upload service has now been decommissioned. For more details, alternatives, and help, please read the announcement on Galaxy Help.
bal community has created a continuously updated list of laboratories that can host
scientists at all career levels. If your lab can host a scientist -- add your name to the list here. In addition, Galaxy Project has a number of positions at its EU and US sites. Contact us at
Світова наукова спільнота створила список лабораторій, що постійно оновлюється та які можуть прийняти українських науковців усіх рівнів, у тому числі й аспірантів. Якщо ваша лабораторія має можливість запросити -- ви можете додати ваше ім’я до списку тут. Окрім того, Galaxy Project має відкриті вакансії у своїх європейських та американських осередках. Пишіть нам на
Научное сообщество создало постоянно обновляемый список лабораторий, которые могут принять украинских ученых (включая аспирантов). К тому же, Galaxy Project имеет открытые позиции на своих европейских и американских сайтах. Контактируйте нас используя
Galaxy is an open source, web-based platform for data intensive biomedical research. If you are new to Galaxy start here or consult our help resources. You can install your own Galaxy by following the tutorial and choose from thousands of tools from the Tool Shed.
Galaxy version 22.05.1, commit e1561ad66582e31816320c90f1e8ec9d5bc698e5
History
search datasets
Tutorial
0 B
0
This history is empty.
You can load your own data or get data from an external source.

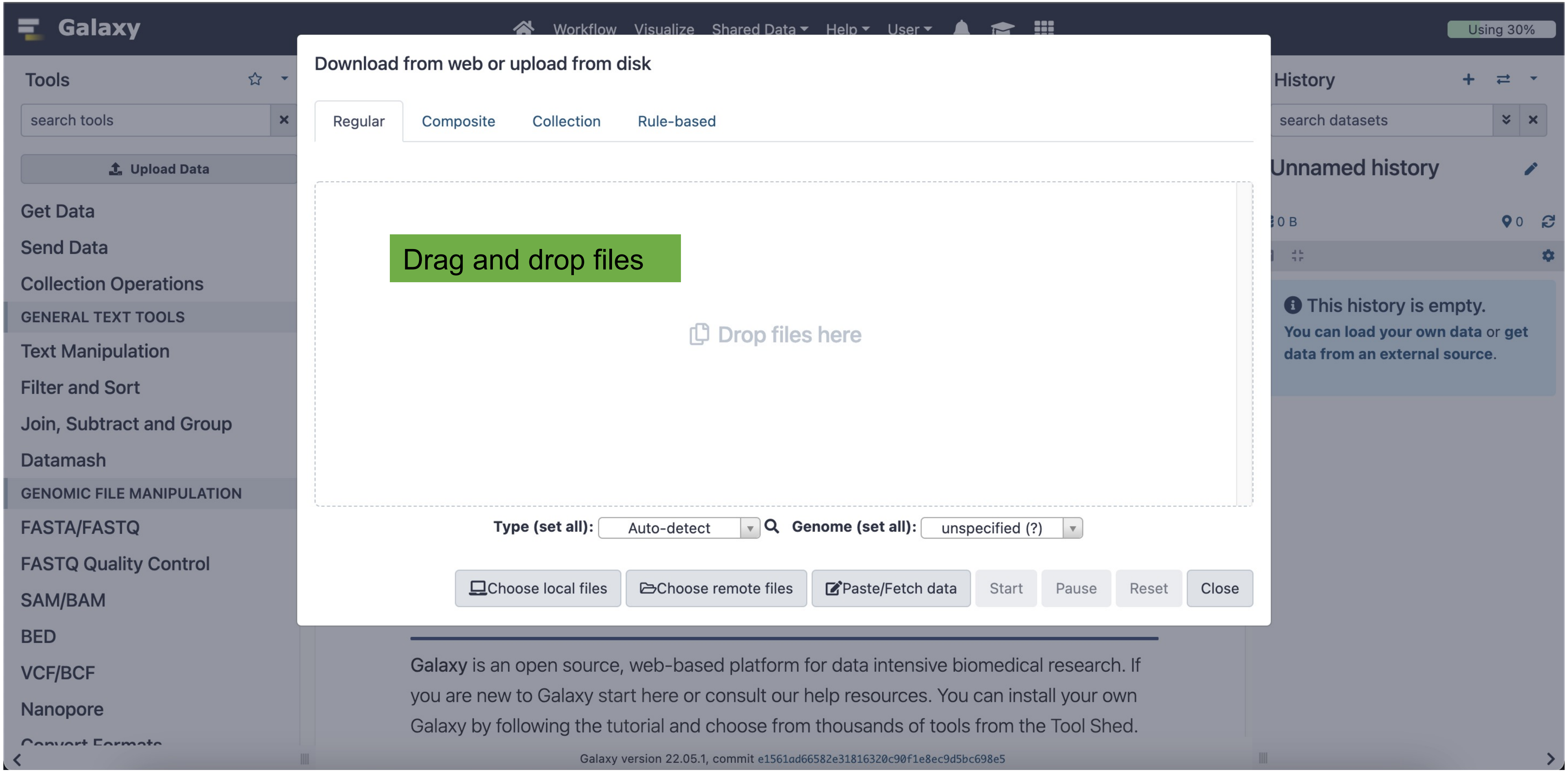
Galaxy
Workflow
Visualize
Shared Data
Help
User
Using 30%
Tools
search tools
Upload Data
Get Data
Send Data
Collection Operations
GENERAL TEXT TOOLS
Text Manipulation
Filter and Sort
Join, Subtract and Group
Datamash
GENOMIC FILE MANIPULATION
FASTA/FASTQ
FASTQ Quality Control
SAM/BAM
BED
VCF/BCF
Nanopore
Download from web or upload from disk
Regular
Composite
Collection
Rule-based
Drag and drop files
Drop files here
Type (set all):
Auto-detect
Genome (set all):
unspecified (?)
Choose local files
Choose remote files
Paste/Fetch data
Start
Pause
Reset
Close
History
search datasets
Unnamed history
0 B
0
This history is empty.
You can load your own data or get data from an external source.
Galaxy is an open source, web-based platform for data intensive biomedical research. If you are new to Galaxy start here or consult our help resources. You can install your own Galaxy by following the tutorial and choose from thousands of tools from the Tool Shed.
Galaxy version 22.05.1, commit e1561ad66582e31816320c90f1e8ec9d5bc698e5

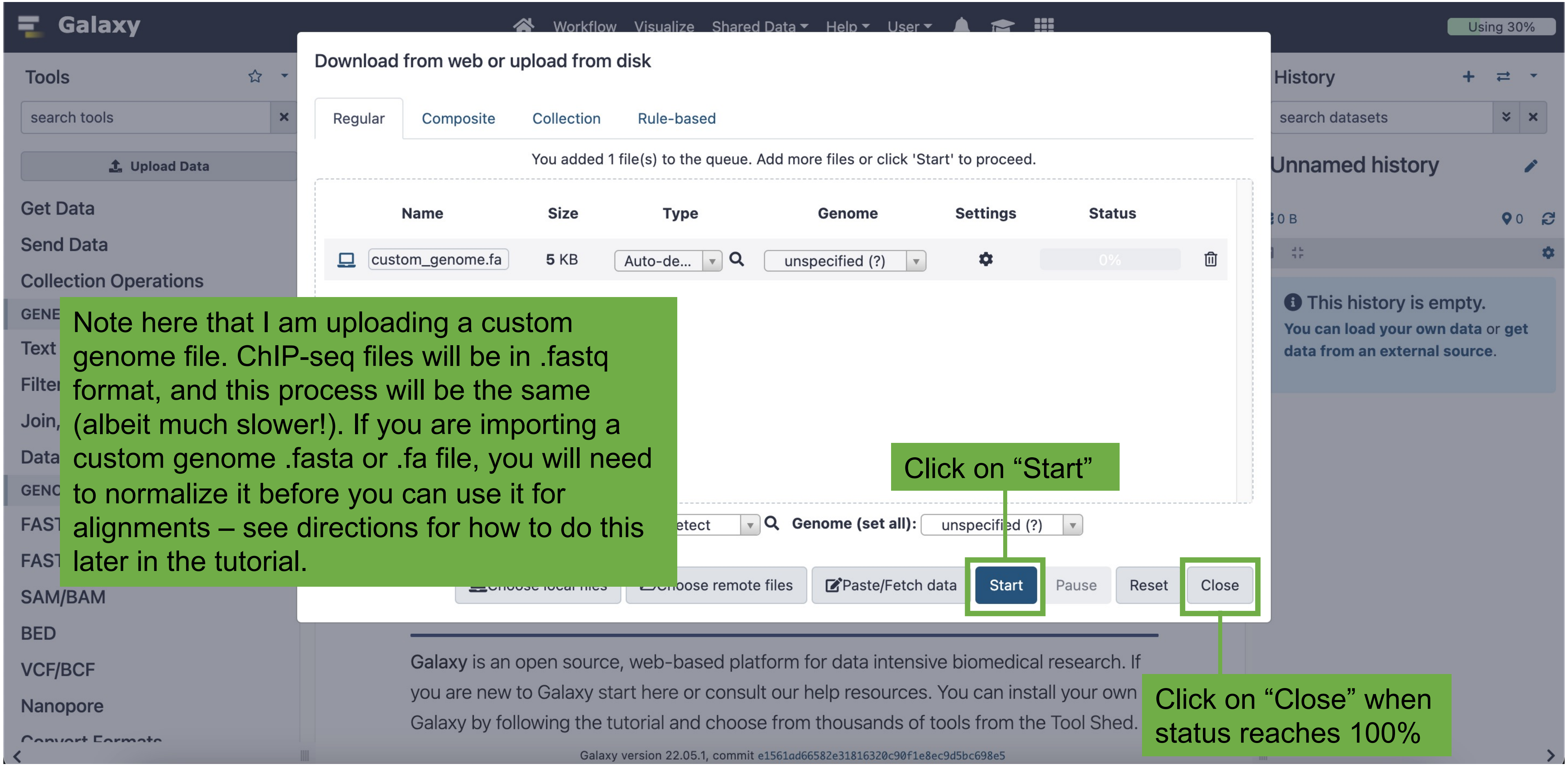
Galaxy
Workflow Visualize Shared Data Help User
Using 30%
Tools
search tools
Upload Data
Get Data
Send Data
Collection Operations
SAM/BAM
BED
VCF/BCF
Nanopore
Download from web or upload from disk
Regular Composite Collection Rule-based
You added 1 file(s) to the queue. Add more files or click 'Start' to proceed.
Name Size Type Genome Settings Status
custom_genome.fa 5 KB Auto-de... unspecified (?) 0%
Genome (set all): unspecified (?)
Choose remote files Paste/Fetch data Start Pause Reset Close
History
search datasets
Unnamed history
This history is empty.
You can load your own data or get data from an external source.
Galaxy is an open source, web-based platform for data intensive biomedical research. If you are new to Galaxy start here or consult our help resources. You can install your own Galaxy by following the tutorial and choose from thousands of tools from the Tool Shed.
Galaxy version 22.05.1, commit e1561ad66582e31816320c90f1e8ec9d5bc698e5
Note here that I am uploading a custom genome file. ChIP-seq files will be in .fastq format, and this process will be the same (albeit much slower!). If you are importing a custom genome .fasta or .fa file, you will need to normalize it before you can use it for alignments – see directions for how to do this later in the tutorial.
Click on “Start”
Click on “Close” when status reaches 100%

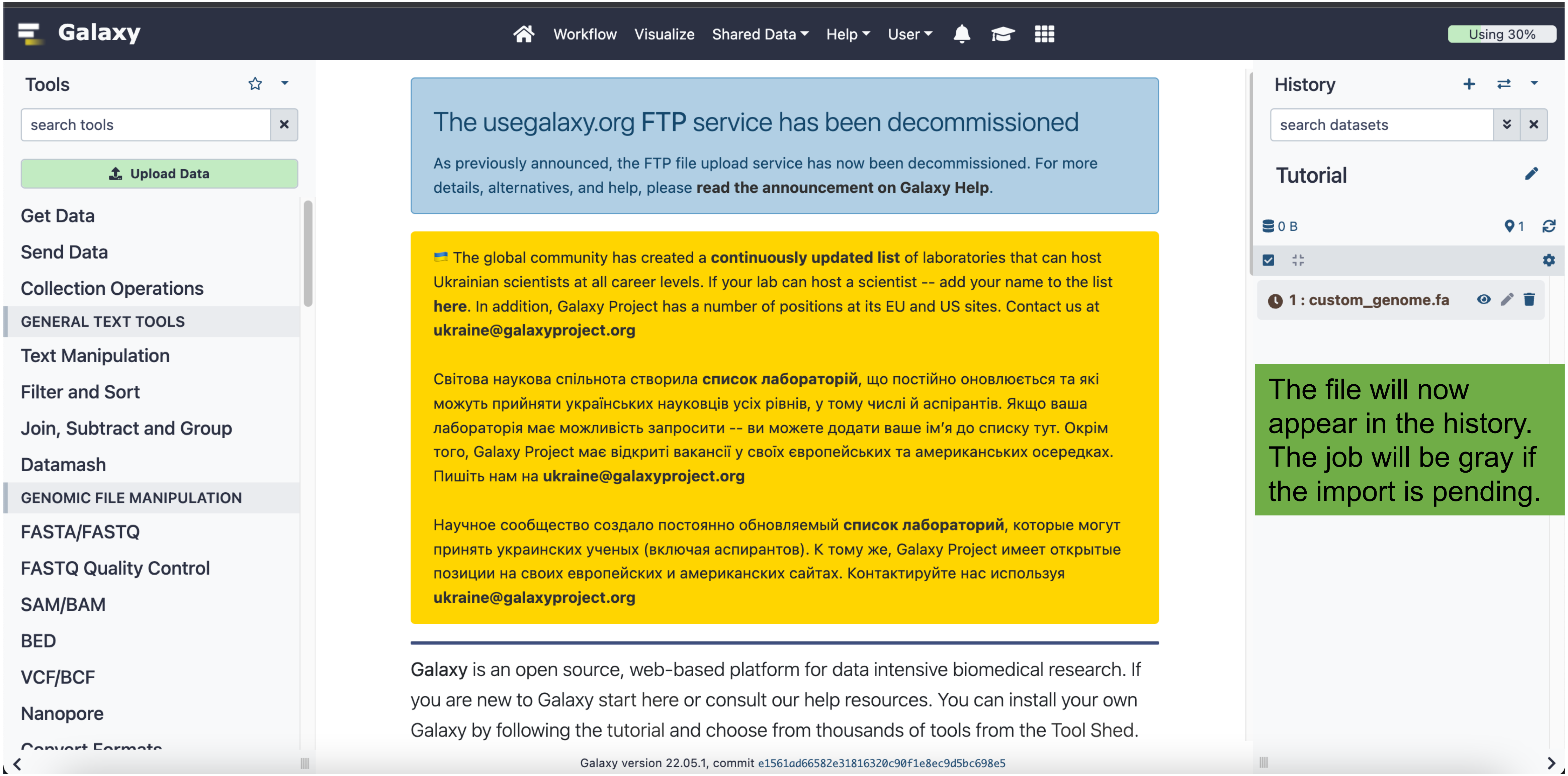
Galaxy
Workflow
Visualize
Shared Data
Help
User
Using 30%
Tools
search tools
Upload Data
Get Data
Send Data
Collection Operations
GENERAL TEXT TOOLS
Text Manipulation
Filter and Sort
Join, Subtract and Group
Datamash
GENOMIC FILE MANIPULATION
FASTA/FASTQ
FASTQ Quality Control
SAM/BAM
BED
VCF/BCF
Nanopore
The usegalaxy.org FTP service has been decommissioned
As previously announced, the FTP file upload service has now been decommissioned. For more details, alternatives, and help, please read the announcement on Galaxy Help.
The global community has created a continuously updated list of laboratories that can host Ukrainian scientists at all career levels. If your lab can host a scientist -- add your name to the list here. In addition, Galaxy Project has a number of positions at its EU and US sites. Contact us at
Світова наукова спільнота створила список лабораторій, що постійно оновлюється та які можуть прийняти українських науковців усіх рівнів, у тому числі й аспірантів. Якщо ваша лабораторія має можливість запросити -- ви можете додати ваше ім’я до списку тут. Окрім того, Galaxy Project має відкриті вакансії у своїх європейських та американських осередках. Пишіть нам на
Научное сообщество создало постоянно обновляемый список лабораторий, которые могут принять украинских ученых (включая аспирантов). К тому же, Galaxy Project имеет открытые позиции на своих европейских и американских сайтах. Контактируйте нас используя
Galaxy is an open source, web-based platform for data intensive biomedical research. If you are new to Galaxy start here or consult our help resources. You can install your own Galaxy by following the tutorial and choose from thousands of tools from the Tool Shed.
Galaxy version 22.05.1, commit e1561ad66582e31816320c90f1e8ec9d5bc698e5
History
search datasets
Tutorial
0 B
1
1 : custom_genome.fa
The file will now appear in the history. The job will be gray if the import is pending.

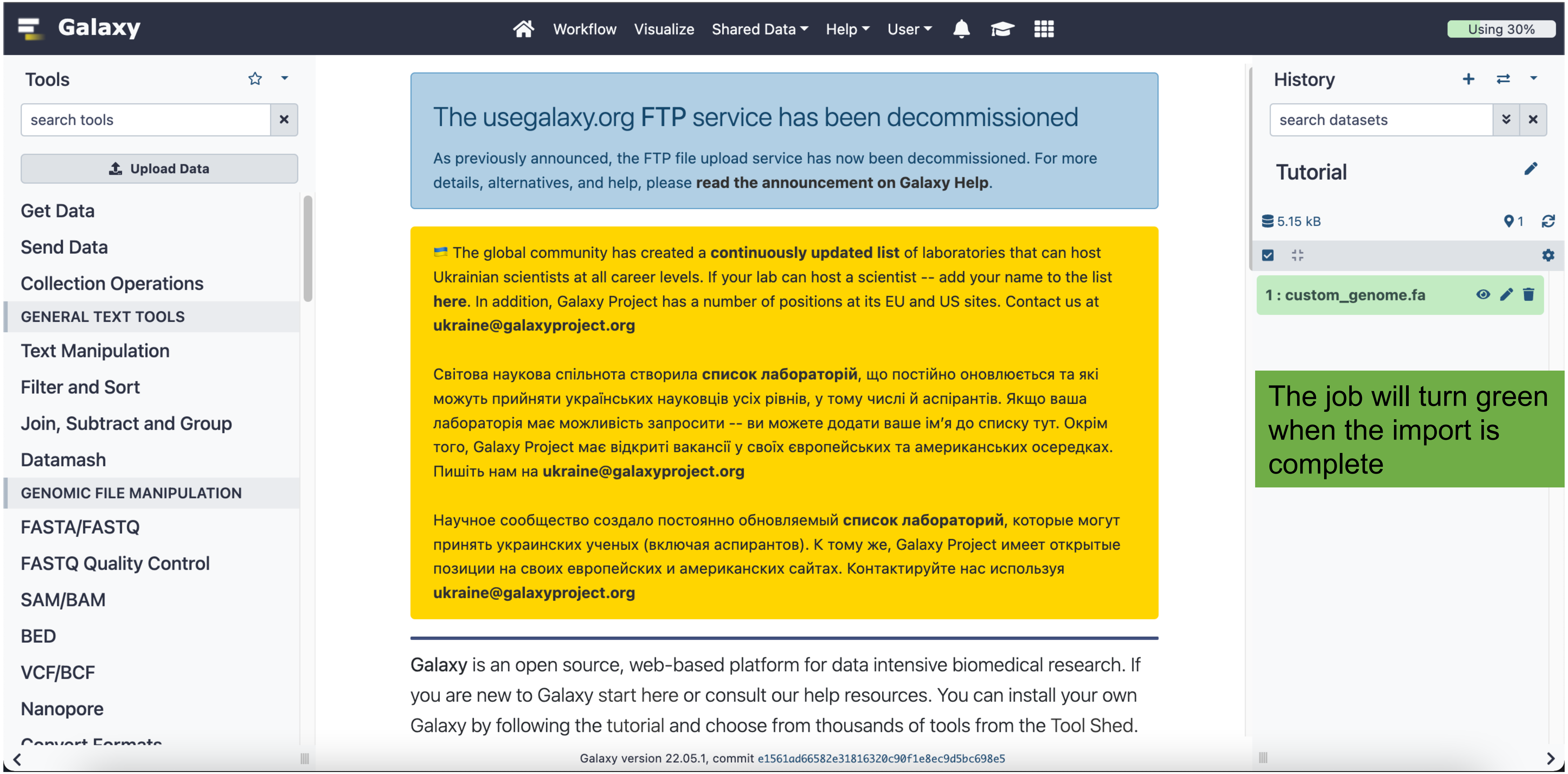
Galaxy
Workflow
Visualize
Shared Data
Help
User
Using 30%
Tools
search tools
Upload Data
Get Data
Send Data
Collection Operations
GENERAL TEXT TOOLS
Text Manipulation
Filter and Sort
Join, Subtract and Group
Datamash
GENOMIC FILE MANIPULATION
FASTA/FASTQ
FASTQ Quality Control
SAM/BAM
BED
VCF/BCF
Nanopore
The usegalaxy.org FTP service has been decommissioned
As previously announced, the FTP file upload service has now been decommissioned. For more details, alternatives, and help, please read the announcement on Galaxy Help.
The global community has created a continuously updated list of laboratories that can host Ukrainian scientists at all career levels. If your lab can host a scientist -- add your name to the list here. In addition, Galaxy Project has a number of positions at its EU and US sites. Contact us at
Світова наукова спільнота створила список лабораторій, що постійно оновлюється та які можуть прийняти українських науковців усіх рівнів, у тому числі й аспірантів. Якщо ваша лабораторія має можливість запросити -- ви можете додати ваше ім’я до списку тут. Окрім того, Galaxy Project має відкриті вакансії у своїх європейських та американських осередках. Пишіть нам на
Научное сообщество создало постоянно обновляемый список лабораторий, которые могут принять украинских ученых (включая аспирантов). К тому же, Galaxy Project имеет открытые позиции на своих европейских и американских сайтах. Контактируйте нас используя
Galaxy is an open source, web-based platform for data intensive biomedical research. If you are new to Galaxy start here or consult our help resources. You can install your own Galaxy by following the tutorial and choose from thousands of tools from the Tool Shed.
Galaxy version 22.05.1, commit e1561ad66582e31816320c90f1e8ec9d5bc698e5
History
search datasets
Tutorial
5.15 kB
1
1 : custom_genome.fa
The job will turn green when the import is complete

### Getting data

#### Option 2: SRA run selector import

Use this option if your data have a GEO accession number

**Preparation of Illumina sequencing libraries**

Data will be available at NCBI GEO Short Read Archive (SRA). GEO release number GSE39271.

Soruco et al., *Genes & Development* 2013

If a research article performs a high-throughput experiment, the authors will typically list the accession number in the Methods or Supplemental Information section. Copy the accession number, and then paste into the search box at NCBI GEO. Alternatively, you can directly search GEO for your factor of interest.

https://www.ncbi.nlm.nih.gov/geo/

#### Gene Expression Omnibus

GEO is a public functional genomics data repository supporting MIAME-compliant data submissions. Array- and sequence-based data are accepted. Tools are provided to help users query and download experiments and curated gene expression profiles.

GSE39271 **Search**

NCBI | Gene Expression Omnibus

HOME | SEARCH | SITE MAP | GEO Publications | FAQ | MIAME | Email GEO

NCBI > GEO > **Accession Display** | Not logged in | Login

Scope: Self | Format: HTML | Amount: Quick | GEO accession: GSE39271 | GO

| **Series GSE39271** | Query DataSets for GSE39271 |
|---|---|
| Status | Public on Jul 15, 2013 |
| Title | Synergistic interactions between CLAMP and MSL complex facilitate Drosophila dosage compensation |
| Organism | Drosophila melanogaster |
| Experiment type | Genome binding/occupancy profiling by high throughput sequencing<br>Expression profiling by high throughput sequencing |
| Summary | ChIP-seq and mRNA-seq experiments were performed to understand the role of the CLAMP protein in dosage compensation |
| Overall design | ChIP-seq experiments compared the binding profiles of CLAMP in male and female cells and mRNA-seq data to define the role of CLAMP in regulating genes on the X-chromosome |
| Contributor(s) | Larschan E, Bishop E |
| Citation(s) | Soruco MM, Chery J, Bishop EP, Siggers T et al. The CLAMP protein links the MSL complex to the X chromosome during Drosophila dosage compensation. *Genes Dev* 2013 Jul 15;27(14):1551-6. PMID: 23873939 |
| Submission date | Jul 11, 2012 |
| Last update date | May 15, 2019 |
| Contact name | Erica Larschan |
| E-mail(s) | |
| Organization name | Brown University |
| Street address | 185 Meeting St. |
| City | Providence |
| ZIP/Postal code | 02912 |
| Country | USA |

The landing page will have information about the authors and the experiment

Country USA

Platforms (1) GPL13304 Illumina HiSeq 2000 (Drosophila melanogaster)

Samples (15)
More...

GSM959255 CLAMP ChIP Kc cells control RNAi replicate 1 Input
GSM959256 CLAMP ChIP Kc cells control RNAi replicate 1
GSM959257 CLAMP ChIP Kc cells control RNAi replicate 2 Input

**Relations**

BioProject PRJNA170445
SRA SRP014195

| Download family | Format |
|---|---|
| SOFT formatted family file(s) | SOFT |
| MINiML formatted family file(s) | MINiML |
| Series Matrix File(s) | TXT |

| Supplementary file | Size | Download | File type/resource |
|---|---|---|---|
| GSE39271_RAW.tar | 1.0 Mb | (http) (custom) | TAR (of RPKM) |
| GSE39271_chip_kccells_clamp_gfp_rnai_combined.wig.gz | 13.0 Mb | (ftp)(http) | WIG |
| GSE39271_chip_s2cells_clamp_gfp_rnai_combined.wig.gz | 10.2 Mb | (ftp)(http) | WIG |
| GSE39271_chip_s2cells_clamp_msl2_rnai_combined.wig.gz | 13.2 Mb | (ftp)(http) | WIG |

SRA Run Selector

*Raw data are available in SRA*
*Processed data provided as supplementary file*
*Processed data are available on Series record*

Scroll to the bottom of the landing page and click on the "SRA Run Selector" link

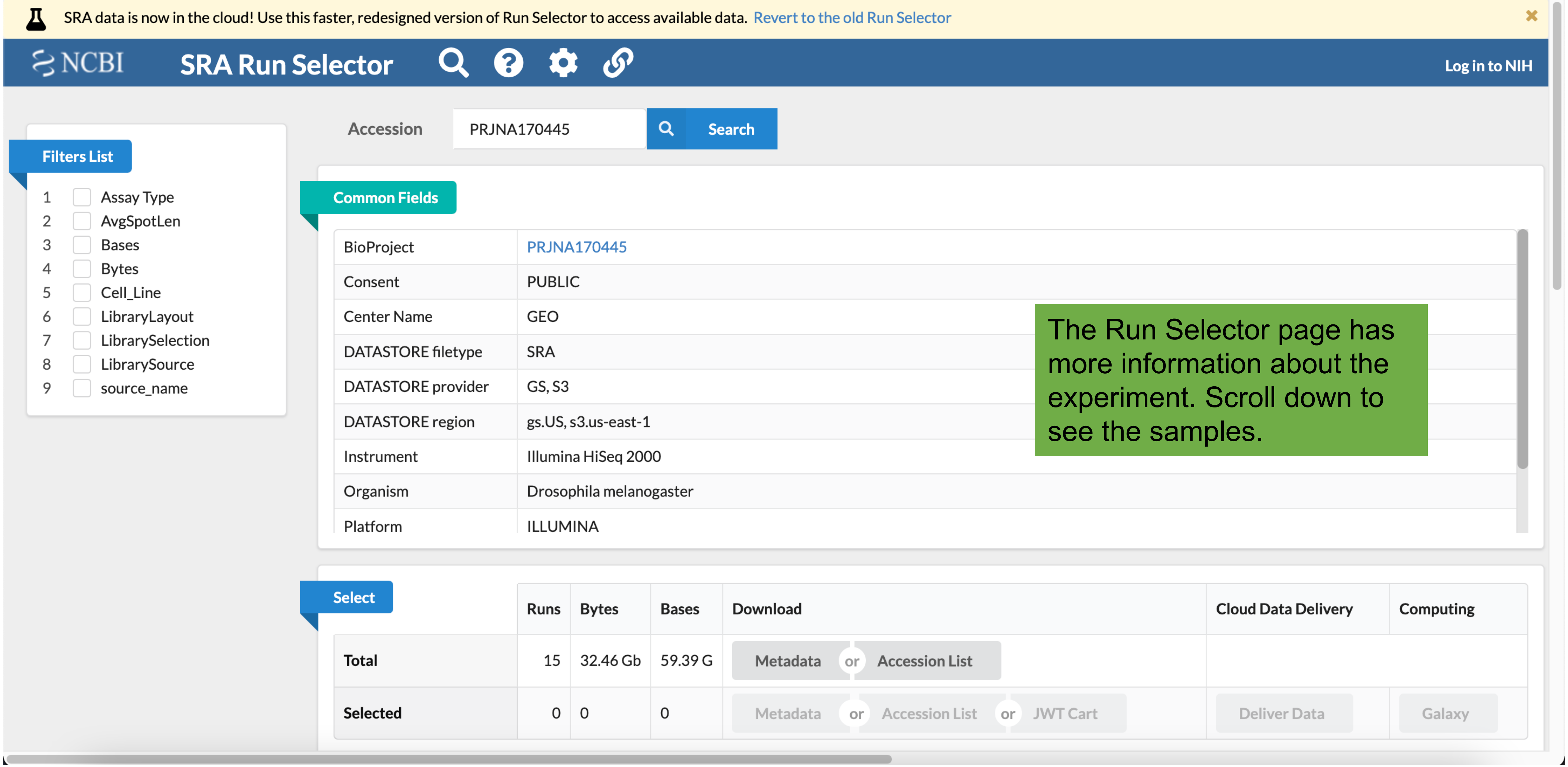


The Run Selector page has more information about the experiment. Scroll down to see the samples.

NCBI SRA Run Selector

NCBI SRA Run Selector Log in to NIH

Found 15 Items

| | Run | BioSample | Assay Type | AvgSpotLen | Bases | Bytes | Cell_Line | Experiment | GEO_Accession | Library Name | LibraryLayout | LibrarySelection | LibrarySource | Sample Name | source_name |
|---|---|---|---|---|---|---|---|---|---|---|---|---|---|---|---|
| 1 | SRR520444 | SAMN01087358 | ChIP-Seq | 50 | 3.73 G | 1.99 Gb | Kc | SRX159163 | GSM959255 | GSM959255: CLAMP ChIP Kc cells control RNAi replicate 1 Input | SINGLE | ChIP | GENOMIC | GSM959255 | CLAMP ChIP Kc cells control RNAi |
| 2 | SRR520445 | SAMN01087359 | ChIP-Seq | [illegible] | [illegible] | [illegible] | [illegible] | [illegible]9164 | GSM959256 | GSM959256: CLAMP ChIP Kc cells control RNAi replicate 1 | SINGLE | ChIP | GENOMIC | GSM959256 | CLAMP ChIP Kc cells control RNAi |
| 3 | SRR520446 | SAMN01087360 | ChIP-Seq | [illegible] | [illegible] | [illegible] | [illegible] | [illegible]9165 | GSM959257 | GSM959257: CLAMP ChIP Kc cells control RNAi replicate 2 Input | SINGLE | ChIP | GENOMIC | GSM959257 | CLAMP ChIP Kc cells control RNAi |
| 4 | SRR520447 | SAMN01087361 | ChIP-Seq | [illegible] | [illegible] | [illegible] | [illegible] | [illegible]9166 | GSM959258 | GSM959258: CLAMP ChIP Kc cells control RNAi replicate 2 | SINGLE | ChIP | GENOMIC | GSM959258 | CLAMP ChIP Kc cells control RNAi |
| 5 | SRR520448 | SAMN01087362 | ChIP-Seq | 50 | 3.45 G | 1.88 Gb | SL2 | SRX159167 | GSM959259 | GSM959259: CLAMP ChIP SL2 cells MSL2 RNAi replicate 1 Input | SINGLE | ChIP | GENOMIC | GSM959259 | CLAMP ChIP SL2 cells MSL2 RNAi |
| 6 | SRR520449 | SAMN01087363 | ChIP-Seq | 50 | 3.24 G | 1.69 Gb | SL2 | SRX159168 | GSM959260 | GSM959260: CLAMP ChIP SL2 cells MSL2 RNAi replicate 1 | SINGLE | ChIP | GENOMIC | GSM959260 | CLAMP ChIP SL2 cells MSL2 RNAi |
| 7 | SRR520450 | SAMN01087364 | ChIP-Seq | 50 | 2.19 G | 1.21 Gb | SL2 | SRX159169 | GSM959261 | GSM959261: CLAMP ChIP SL2 cells MSL2 RNAi replicate 2 Input | SINGLE | ChIP | GENOMIC | GSM959261 | CLAMP ChIP SL2 cells MSL2 RNAi |
| 8 | SRR520451 | SAMN01087365 | ChIP-Seq | 50 | 1.89 G | 1.06 Gb | SL2 | SRX159170 | GSM959262 | GSM959262: CLAMP ChIP SL2 cells MSL2 RNAi replicate 2 | SINGLE | ChIP | GENOMIC | GSM959262 | CLAMP ChIP SL2 cells MSL2 RNAi |
| 9 | SRR520452 | SAMN01087366 | ChIP-Seq | 36 | 1.06 G | 569.88 Mb | SL2 | SRX159171 | GSM959263 | GSM959263: CLAMP ChIP SL2 cells control RNAi replicate 1 Input | SINGLE | ChIP | GENOMIC | GSM959263 | CLAMP ChIP SL2 cells control RNAi |
| 10 | SRR520453 | SAMN01087367 | ChIP-Seq | 36 | 1.00 G | 531.81 Mb | SL2 | SRX159172 | GSM959264 | GSM959264: CLAMP ChIP SL2 cells control RNAi replicate 1 | SINGLE | ChIP | GENOMIC | GSM959264 | CLAMP ChIP SL2 cells control RNAi |
| 11 | SRR520454 | SAMN01087368 | ChIP-Seq | 42 | 944.92 M | 446.79 Mb | SL2 | SRX159173 | GSM959265 | GSM959265: CLAMP ChIP SL2 cells control RNAi replicate 2 Input | SINGLE | ChIP | GENOMIC | GSM959265 | CLAMP ChIP SL2 cells control RNAi |
| 12 | SRR520455 | SAMN01087369 | ChIP-Seq | 42 | 1.19 G | 586.17 Mb | SL2 | SRX159174 | GSM959266 | GSM959266: CLAMP ChIP SL2 cells control RNAi replicate 2 | SINGLE | ChIP | GENOMIC | GSM959266 | CLAMP ChIP SL2 cells control RNAi |
| 13 | SRR520456 | SAMN01087370 | RNA-Seq | 100 | 11.09 G | 6.17 Gb | SL2 | SRX159175 | GSM959267 | GSM959267: mRNA-seq CLAMP RNAi SL2 | PAIRED | cDNA | TRANSCRIPTOMIC | GSM959267 | mRNA-seq CLAMP RNAi SL2 |
| 14 | SRR520457 | SAMN01087371 | RNA-Seq | 100 | 11.45 G | 6.33 Gb | SL2 | SRX159176 | GSM959268 | GSM959268: mRNA-seq Control RNAi SL2 | PAIRED | cDNA | TRANSCRIPTOMIC | GSM959268 | mRNA-seq Control RNAi SL2 |
| 15 | SRR520458 | SAMN01087372 | RNA-Seq | 100 | 11.07 G | 6.10 Gb | SL2 | SRX159177 | GSM959269 | GSM959269: mRNA-seq MSL2 RNAi SL2 | PAIRED | cDNA | TRANSCRIPTOMIC | GSM959269 | mRNA-seq MSL2 RNAi SL2 |

ChIP-seq samples from an RNAi background

RNA-seq samples

There may be a lot of samples, and not every run might be useful. This is a large dataset with many samples, including RNA-seq (which must be analyzed by different bioinformatic tools) and ChIP-seq in the background of RNAi for a different factor (which will not be informative for wild-type situations).

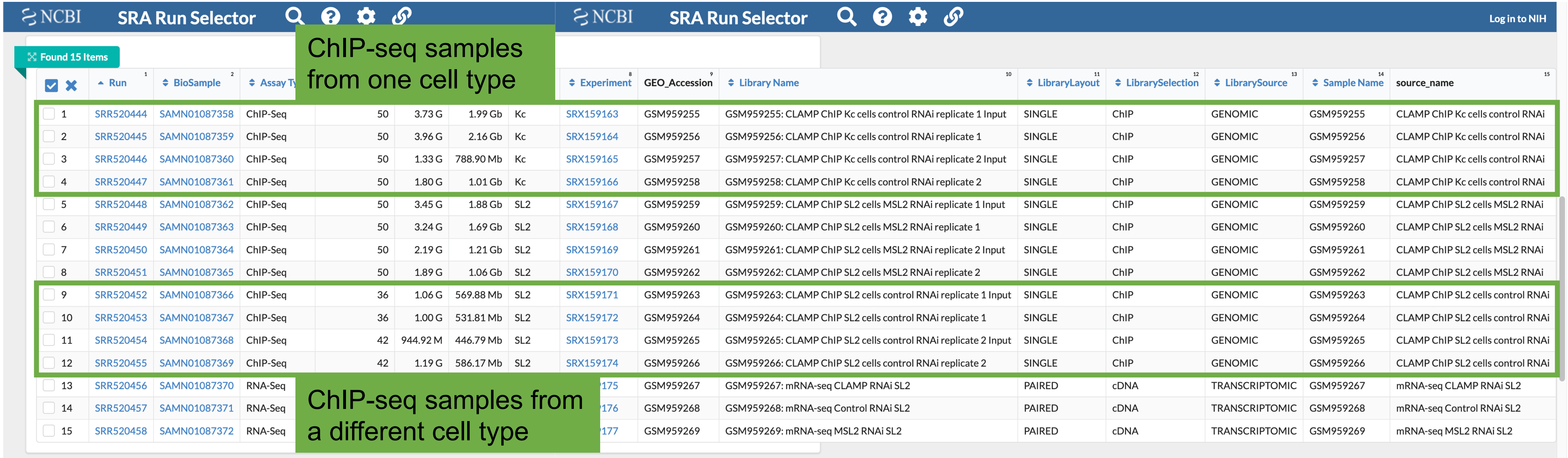


| | Run | BioSample | Assay Ty | [illegible] | [illegible] | [illegible] | [illegible] | Experiment | GEO_Accession | Library Name | LibraryLayout | LibrarySelection | LibrarySource | Sample Name | source_name |
|---|---|---|---|---|---|---|---|---|---|---|---|---|---|---|---|
| 1 | SRR520444 | SAMN01087358 | ChIP-Seq | 50 | 3.73 G | 1.99 Gb | Kc | SRX159163 | GSM959255 | GSM959255: CLAMP ChIP Kc cells control RNAi replicate 1 Input | SINGLE | ChIP | GENOMIC | GSM959255 | CLAMP ChIP Kc cells control RNAi |
| 2 | SRR520445 | SAMN01087359 | ChIP-Seq | 50 | 3.96 G | 2.16 Gb | Kc | SRX159164 | GSM959256 | GSM959256: CLAMP ChIP Kc cells control RNAi replicate 1 | SINGLE | ChIP | GENOMIC | GSM959256 | CLAMP ChIP Kc cells control RNAi |
| 3 | SRR520446 | SAMN01087360 | ChIP-Seq | 50 | 1.33 G | 788.90 Mb | Kc | SRX159165 | GSM959257 | GSM959257: CLAMP ChIP Kc cells control RNAi replicate 2 Input | SINGLE | ChIP | GENOMIC | GSM959257 | CLAMP ChIP Kc cells control RNAi |
| 4 | SRR520447 | SAMN01087361 | ChIP-Seq | 50 | 1.80 G | 1.01 Gb | Kc | SRX159166 | GSM959258 | GSM959258: CLAMP ChIP Kc cells control RNAi replicate 2 | SINGLE | ChIP | GENOMIC | GSM959258 | CLAMP ChIP Kc cells control RNAi |
| 5 | SRR520448 | SAMN01087362 | ChIP-Seq | 50 | 3.45 G | 1.88 Gb | SL2 | SRX159167 | GSM959259 | GSM959259: CLAMP ChIP SL2 cells MSL2 RNAi replicate 1 Input | SINGLE | ChIP | GENOMIC | GSM959259 | CLAMP ChIP SL2 cells MSL2 RNAi |
| 6 | SRR520449 | SAMN01087363 | ChIP-Seq | 50 | 3.24 G | 1.69 Gb | SL2 | SRX159168 | GSM959260 | GSM959260: CLAMP ChIP SL2 cells MSL2 RNAi replicate 1 | SINGLE | ChIP | GENOMIC | GSM959260 | CLAMP ChIP SL2 cells MSL2 RNAi |
| 7 | SRR520450 | SAMN01087364 | ChIP-Seq | 50 | 2.19 G | 1.21 Gb | SL2 | SRX159169 | GSM959261 | GSM959261: CLAMP ChIP SL2 cells MSL2 RNAi replicate 2 Input | SINGLE | ChIP | GENOMIC | GSM959261 | CLAMP ChIP SL2 cells MSL2 RNAi |
| 8 | SRR520451 | SAMN01087365 | ChIP-Seq | 50 | 1.89 G | 1.06 Gb | SL2 | SRX159170 | GSM959262 | GSM959262: CLAMP ChIP SL2 cells MSL2 RNAi replicate 2 | SINGLE | ChIP | GENOMIC | GSM959262 | CLAMP ChIP SL2 cells MSL2 RNAi |
| 9 | SRR520452 | SAMN01087366 | ChIP-Seq | 36 | 1.06 G | 569.88 Mb | SL2 | SRX159171 | GSM959263 | GSM959263: CLAMP ChIP SL2 cells control RNAi replicate 1 Input | SINGLE | ChIP | GENOMIC | GSM959263 | CLAMP ChIP SL2 cells control RNAi |
| 10 | SRR520453 | SAMN01087367 | ChIP-Seq | 36 | 1.00 G | 531.81 Mb | SL2 | SRX159172 | GSM959264 | GSM959264: CLAMP ChIP SL2 cells control RNAi replicate 1 | SINGLE | ChIP | GENOMIC | GSM959264 | CLAMP ChIP SL2 cells control RNAi |
| 11 | SRR520454 | SAMN01087368 | ChIP-Seq | 42 | 944.92 M | 446.79 Mb | SL2 | SRX159173 | GSM959265 | GSM959265: CLAMP ChIP SL2 cells control RNAi replicate 2 Input | SINGLE | ChIP | GENOMIC | GSM959265 | CLAMP ChIP SL2 cells control RNAi |
| 12 | SRR520455 | SAMN01087369 | ChIP-Seq | 42 | 1.19 G | 586.17 Mb | SL2 | SRX159174 | GSM959266 | GSM959266: CLAMP ChIP SL2 cells control RNAi replicate 2 | SINGLE | ChIP | GENOMIC | GSM959266 | CLAMP ChIP SL2 cells control RNAi |
| 13 | SRR520456 | SAMN01087370 | RNA-Seq | [illegible] | [illegible] | [illegible] | [illegible] | [illegible]175 | GSM959267 | GSM959267: mRNA-seq CLAMP RNAi SL2 | PAIRED | cDNA | TRANSCRIPTOMIC | GSM959267 | mRNA-seq CLAMP RNAi SL2 |
| 14 | SRR520457 | SAMN01087371 | RNA-Seq | [illegible] | [illegible] | [illegible] | [illegible] | [illegible]176 | GSM959268 | GSM959268: mRNA-seq Control RNAi SL2 | PAIRED | cDNA | TRANSCRIPTOMIC | GSM959268 | mRNA-seq Control RNAi SL2 |
| 15 | SRR520458 | SAMN01087372 | RNA-Seq | [illegible] | [illegible] | [illegible] | [illegible] | [illegible]177 | GSM959269 | GSM959269: mRNA-seq MSL2 RNAi SL2 | PAIRED | cDNA | TRANSCRIPTOMIC | GSM959269 | mRNA-seq MSL2 RNAi SL2 |

Here are ChIP-seq runs for the factor of interest in two different cell types. Note that there are 2 replicates and input runs, which allows for normalization of the data.

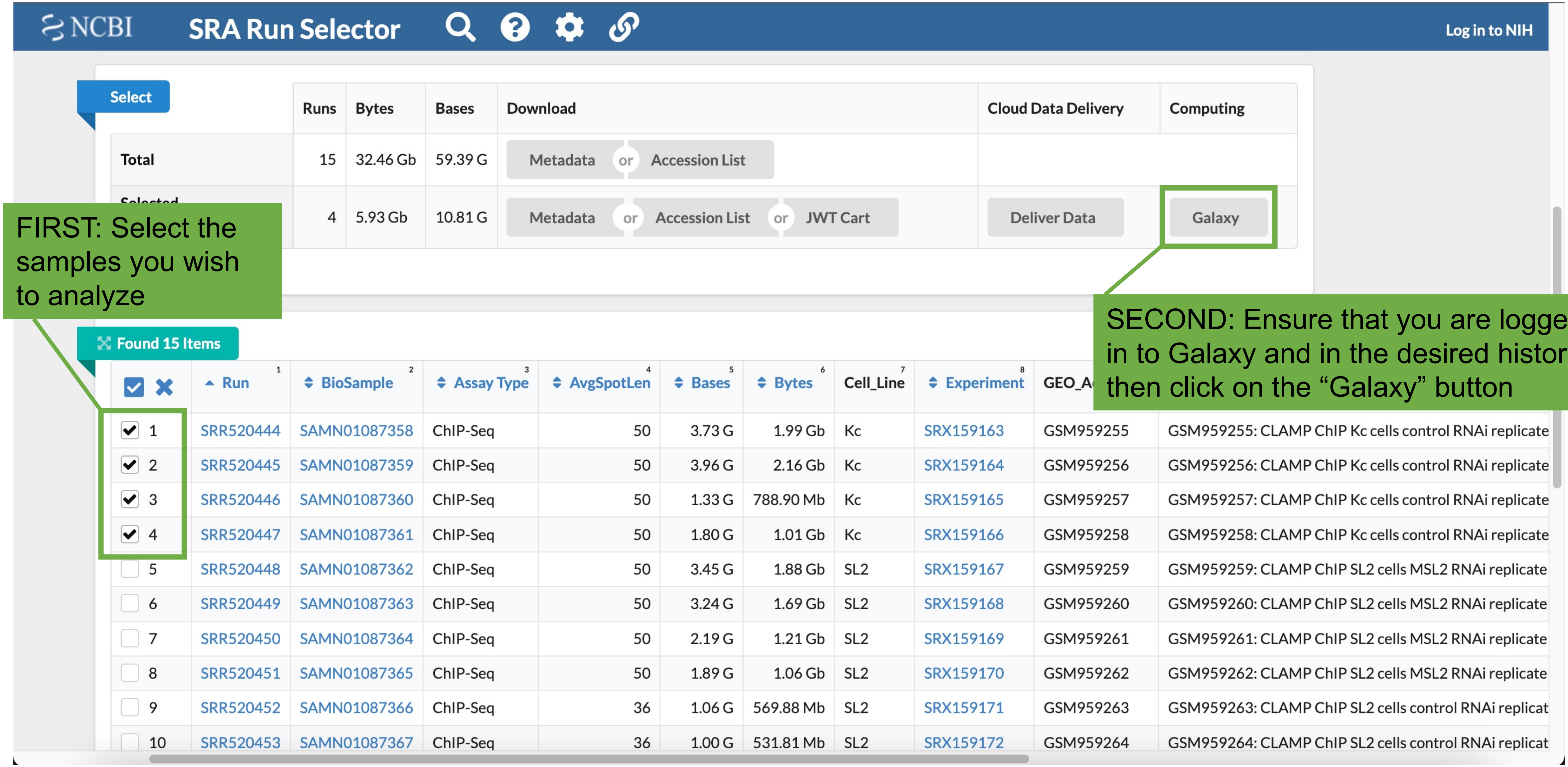


| | Runs | Bytes | Bases | Download | Cloud Data Delivery | Computing |
|---|---|---|---|---|---|---|
| Total | 15 | 32.46 Gb | 59.39 G | Metadata or Accession List | | |
| Selected | 4 | 5.93 Gb | 10.81 G | Metadata or Accession List or JWT Cart | Deliver Data | Galaxy |

| | Run | BioSample | Assay Type | AvgSpotLen | Bases | Bytes | Cell_Line | Experiment | GEO_A[illegible] | [illegible] |
|---|---|---|---|---|---|---|---|---|---|---|
| ☑ 1 | SRR520444 | SAMN01087358 | ChIP-Seq | 50 | 3.73 G | 1.99 Gb | Kc | SRX159163 | GSM959255 | GSM959255: CLAMP ChIP Kc cells control RNAi replicate |
| ☑ 2 | SRR520445 | SAMN01087359 | ChIP-Seq | 50 | 3.96 G | 2.16 Gb | Kc | SRX159164 | GSM959256 | GSM959256: CLAMP ChIP Kc cells control RNAi replicate |
| ☑ 3 | SRR520446 | SAMN01087360 | ChIP-Seq | 50 | 1.33 G | 788.90 Mb | Kc | SRX159165 | GSM959257 | GSM959257: CLAMP ChIP Kc cells control RNAi replicate |
| ☑ 4 | SRR520447 | SAMN01087361 | ChIP-Seq | 50 | 1.80 G | 1.01 Gb | Kc | SRX159166 | GSM959258 | GSM959258: CLAMP ChIP Kc cells control RNAi replicate |
| ☐ 5 | SRR520448 | SAMN01087362 | ChIP-Seq | 50 | 3.45 G | 1.88 Gb | SL2 | SRX159167 | GSM959259 | GSM959259: CLAMP ChIP SL2 cells MSL2 RNAi replicate |
| ☐ 6 | SRR520449 | SAMN01087363 | ChIP-Seq | 50 | 3.24 G | 1.69 Gb | SL2 | SRX159168 | GSM959260 | GSM959260: CLAMP ChIP SL2 cells MSL2 RNAi replicate |
| ☐ 7 | SRR520450 | SAMN01087364 | ChIP-Seq | 50 | 2.19 G | 1.21 Gb | SL2 | SRX159169 | GSM959261 | GSM959261: CLAMP ChIP SL2 cells MSL2 RNAi replicate |
| ☐ 8 | SRR520451 | SAMN01087365 | ChIP-Seq | 50 | 1.89 G | 1.06 Gb | SL2 | SRX159170 | GSM959262 | GSM959262: CLAMP ChIP SL2 cells MSL2 RNAi replicate |
| ☐ 9 | SRR520452 | SAMN01087366 | ChIP-Seq | 36 | 1.06 G | 569.88 Mb | SL2 | SRX159171 | GSM959263 | GSM959263: CLAMP ChIP SL2 cells control RNAi replicat |
| ☐ 10 | SRR520453 | SAMN01087367 | ChIP-Seq | 36 | 1.00 G | 531.81 Mb | SL2 | SRX159172 | GSM959264 | GSM959264: CLAMP ChIP SL2 cells control RNAi replicat |

FIRST: Select the samples you wish to analyze

SECOND: Ensure that you are logged in to Galaxy and in the desired history, then click on the “Galaxy” button

A job has been successfully added to the queue - resulting in the following dataset:

**2 : SRA**

You can check the status of queued jobs and view the resulting data by refreshing the **History** pane. When the job has been run the status will change from 'running' to 'finished' if completed successfully or 'error' if problems were encountered. You are now being redirected back to **Galaxy**.

A new tab will open and this message will briefly appear

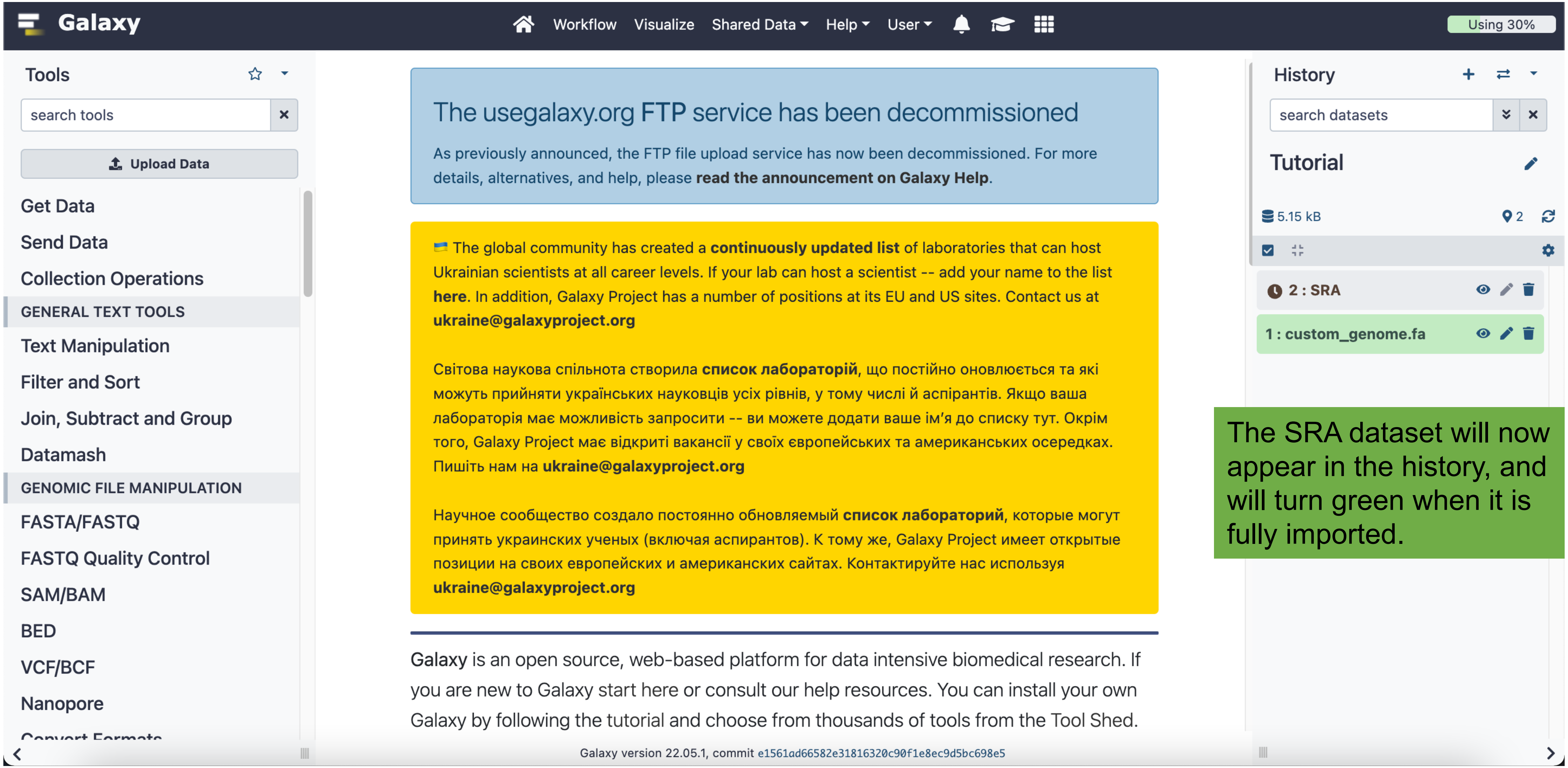


The SRA dataset will now appear in the history, and will turn green when it is fully imported.

Now the SRA reads need to be extracted. Click on “Get Data”, which will open the available options

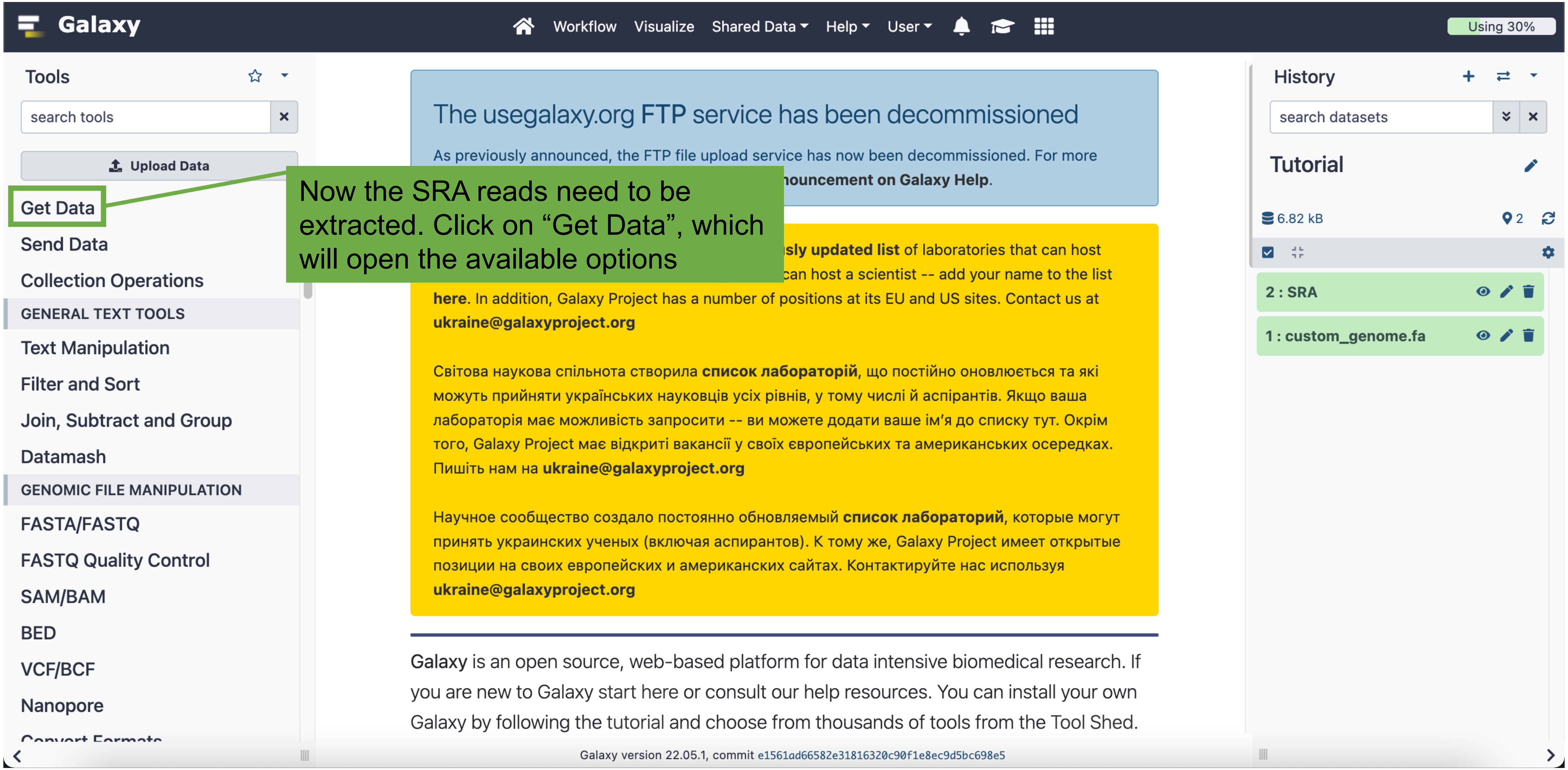

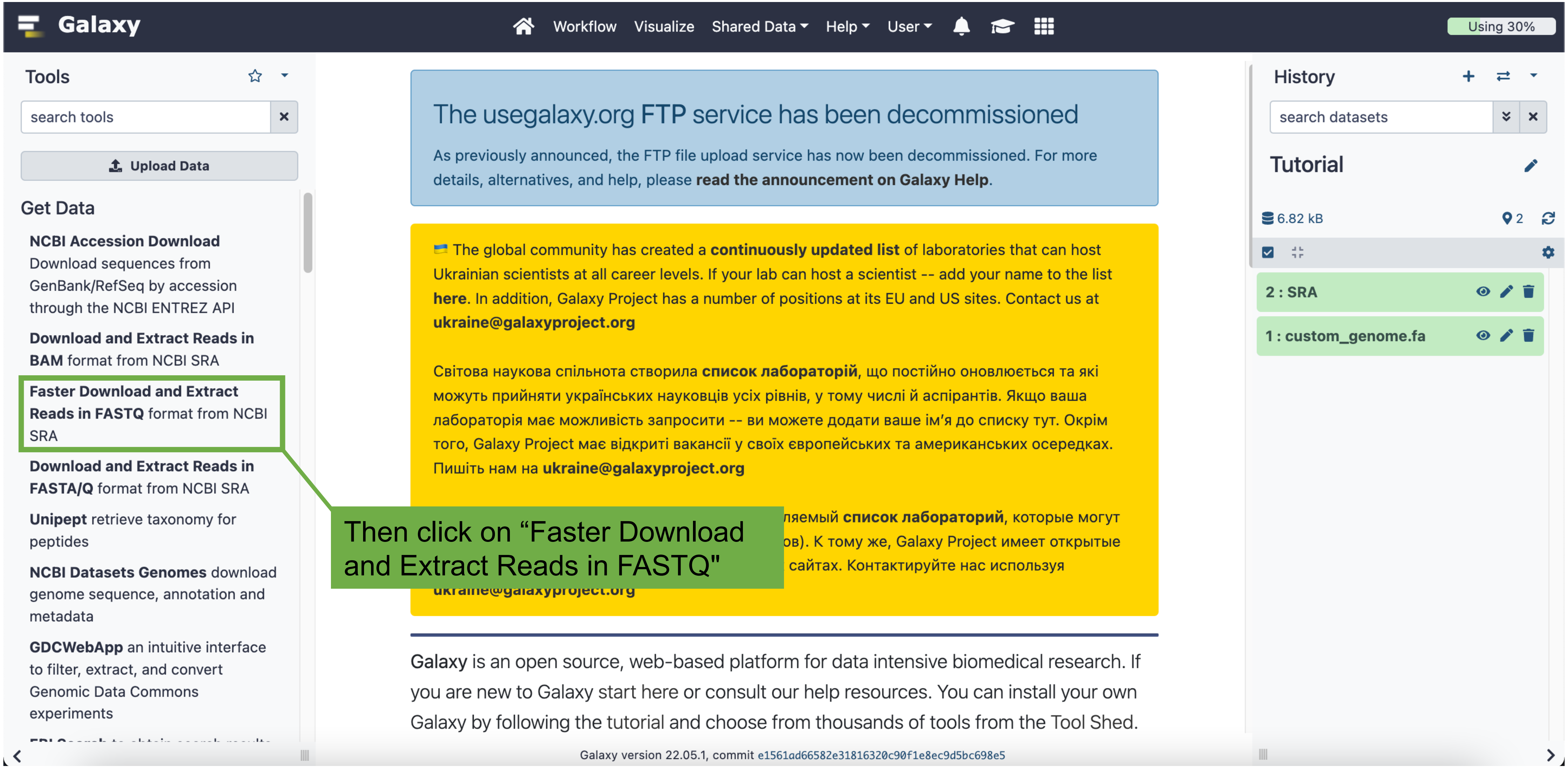
Galaxy
Workflow
Visualize
Shared Data
Help
User
Using 30%
Tools
search tools
Upload Data
Get Data
NCBI Accession Download Download sequences from GenBank/RefSeq by accession through the NCBI ENTREZ API
Download and Extract Reads in BAM format from NCBI SRA
Faster Download and Extract Reads in FASTQ format from NCBI SRA
Download and Extract Reads in FASTA/Q format from NCBI SRA
Unipept retrieve taxonomy for peptides
NCBI Datasets Genomes download genome sequence, annotation and metadata
GDCWebApp an intuitive interface to filter, extract, and convert Genomic Data Commons experiments
The usegalaxy.org FTP service has been decommissioned
As previously announced, the FTP file upload service has now been decommissioned. For more details, alternatives, and help, please read the announcement on Galaxy Help.
The global community has created a continuously updated list of laboratories that can host Ukrainian scientists at all career levels. If your lab can host a scientist -- add your name to the list here. In addition, Galaxy Project has a number of positions at its EU and US sites. Contact us at
Світова наукова спільнота створила список лабораторій, що постійно оновлюється та які можуть прийняти українських науковців усіх рівнів, у тому числі й аспірантів. Якщо ваша лабораторія має можливість запросити -- ви можете додати ваше ім'я до списку тут. Окрім того, Galaxy Project має відкриті вакансії у своїх європейських та американських осередках. Пишіть нам на
Then click on “Faster Download and Extract Reads in FASTQ"
Galaxy is an open source, web-based platform for data intensive biomedical research. If you are new to Galaxy start here or consult our help resources. You can install your own Galaxy by following the tutorial and choose from thousands of tools from the Tool Shed.
Galaxy version 22.05.1, commit e1561ad66582e31816320c90f1e8ec9d5bc698e5
History
search datasets
Tutorial
6.82 kB
2
2 : SRA
1 : custom_genome.fa

Select the blocks icon to change the version to “2.10.9+galaxy0”

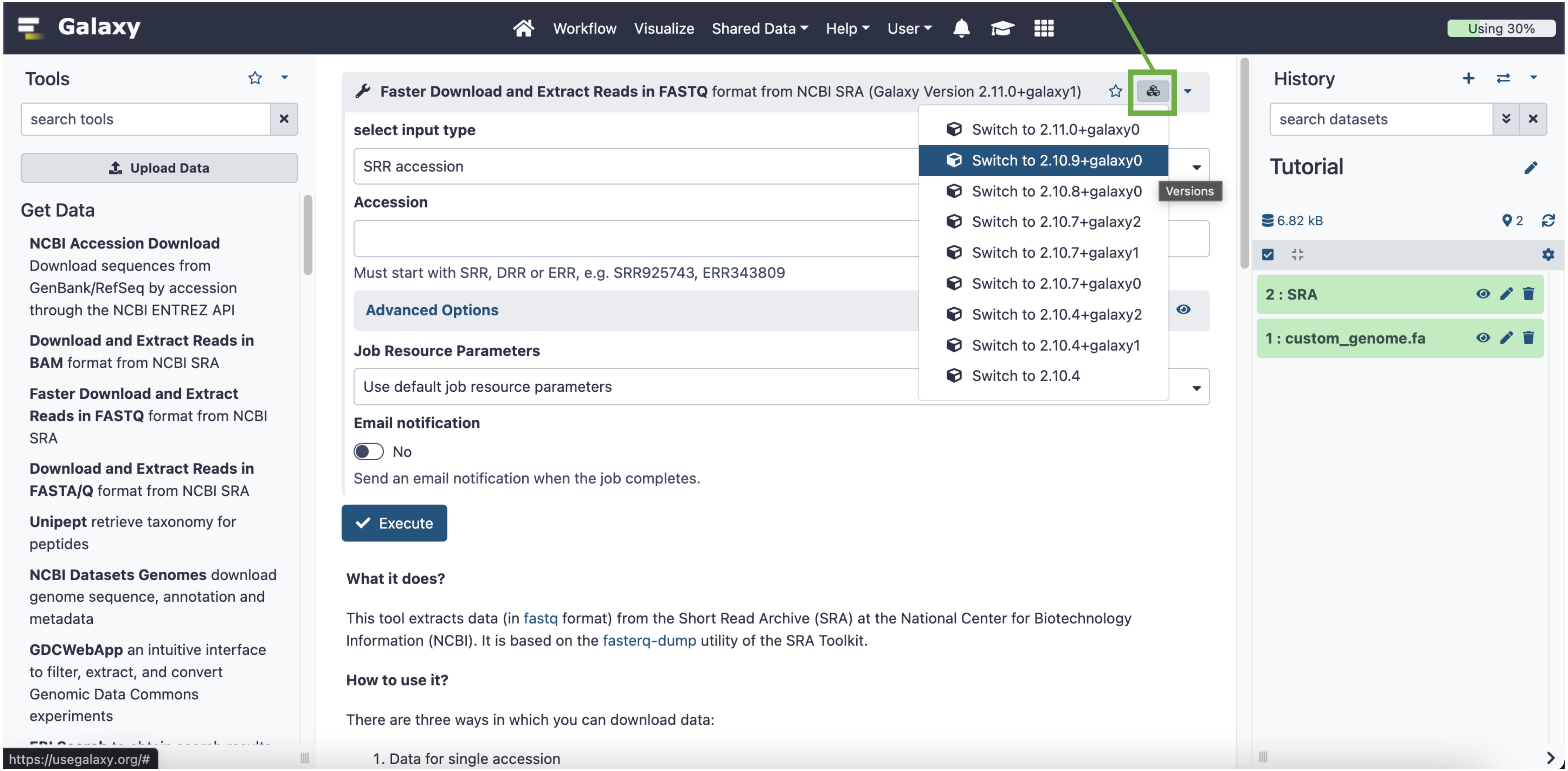

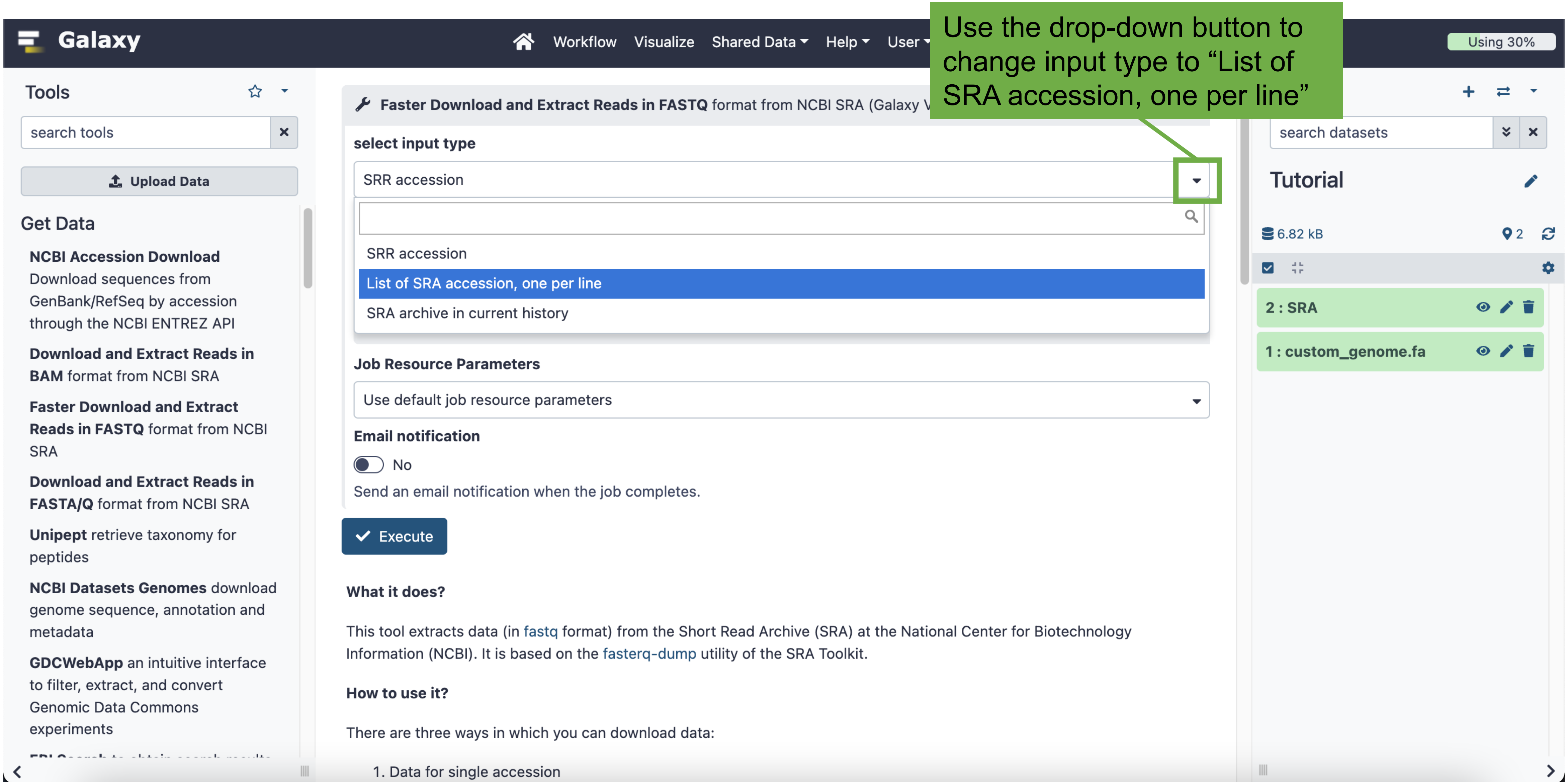
Use the drop-down button to change input type to "List of SRA accession, one per line"
Galaxy
Workflow
Visualize
Shared Data
Help
User
Using 30%
Tools
search tools
Upload Data
Get Data
NCBI Accession Download Download sequences from GenBank/RefSeq by accession through the NCBI ENTREZ API
Download and Extract Reads in BAM format from NCBI SRA
Faster Download and Extract Reads in FASTQ format from NCBI SRA
Download and Extract Reads in FASTA/Q format from NCBI SRA
Unipept retrieve taxonomy for peptides
NCBI Datasets Genomes download genome sequence, annotation and metadata
GDCWebApp an intuitive interface to filter, extract, and convert Genomic Data Commons experiments
Faster Download and Extract Reads in FASTQ format from NCBI SRA (Galaxy V
select input type
SRR accession
SRR accession
List of SRA accession, one per line
SRA archive in current history
Job Resource Parameters
Use default job resource parameters
Email notification
No
Send an email notification when the job completes.
Execute
What it does?
This tool extracts data (in fastq format) from the Short Read Archive (SRA) at the National Center for Biotechnology Information (NCBI). It is based on the fasterq-dump utility of the SRA Toolkit.
How to use it?
There are three ways in which you can download data:
1. Data for single accession
search datasets
Tutorial
6.82 kB
2
2 : SRA
1 : custom_genome.fa

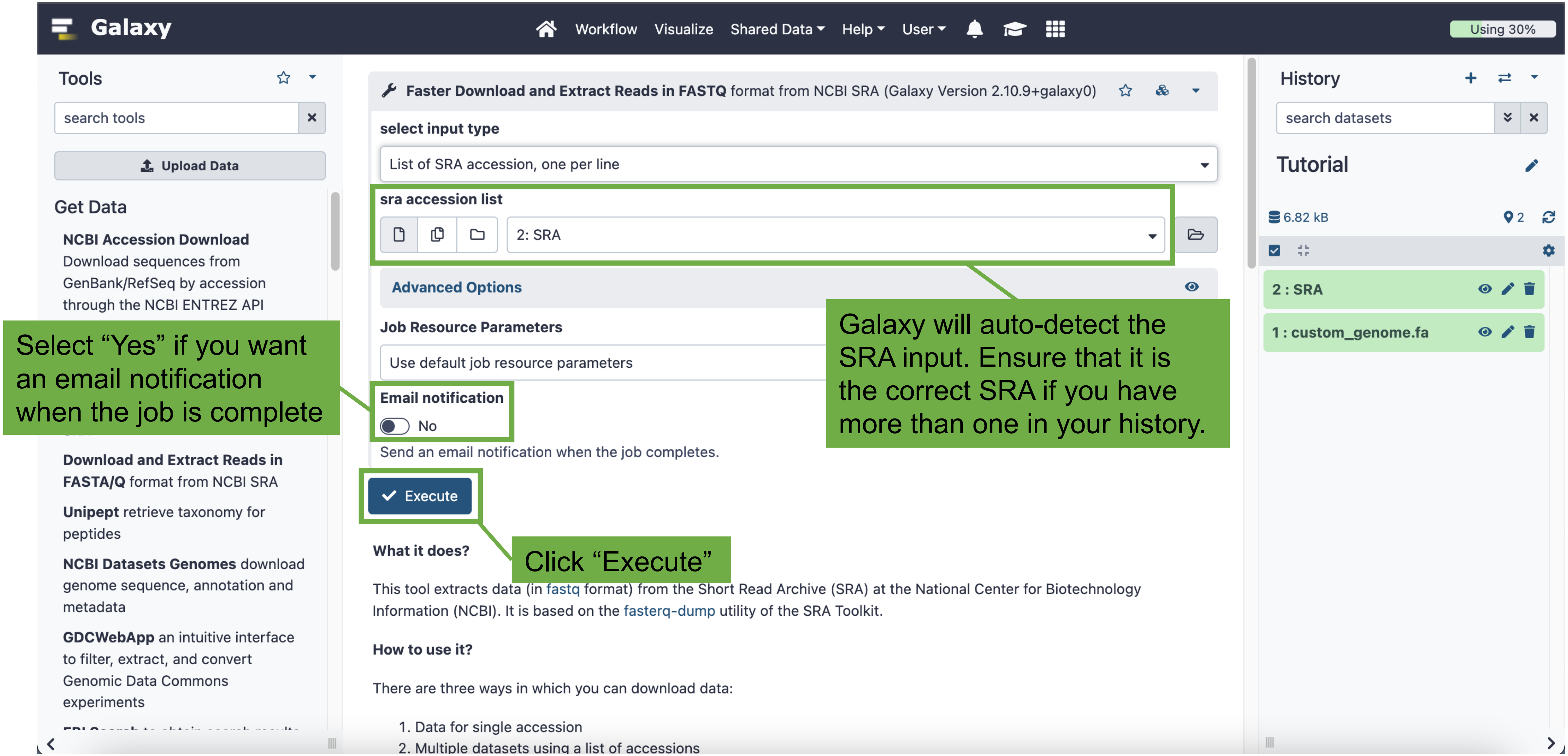

Galaxy
Workflow
Visualize
Shared Data
Help
User
Using 30%
Tools
search tools
Upload Data
Get Data
NCBI Accession Download Download sequences from GenBank/RefSeq by accession through the NCBI ENTREZ API
Download and Extract Reads in FASTA/Q format from NCBI SRA
Unipept retrieve taxonomy for peptides
NCBI Datasets Genomes download genome sequence, annotation and metadata
GDCWebApp an intuitive interface to filter, extract, and convert Genomic Data Commons experiments
Faster Download and Extract Reads in FASTQ format from NCBI SRA (Galaxy Version 2.10.9+galaxy0)
select input type
List of SRA accession, one per line
sra accession list
2: SRA
Advanced Options
Job Resource Parameters
Use default job resource parameters
Email notification
No
Send an email notification when the job completes.
Execute
What it does?
This tool extracts data (in fastq format) from the Short Read Archive (SRA) at the National Center for Biotechnology Information (NCBI). It is based on the fasterq-dump utility of the SRA Toolkit.
How to use it?
There are three ways in which you can download data:
1. Data for single accession
2. Multiple datasets using a list of accessions
History
search datasets
Tutorial
6.82 kB
2
2 : SRA
1 : custom_genome.fa
Select “Yes” if you want an email notification when the job is complete
Galaxy will auto-detect the SRA input. Ensure that it is the correct SRA if you have more than one in your history.
Click “Execute”

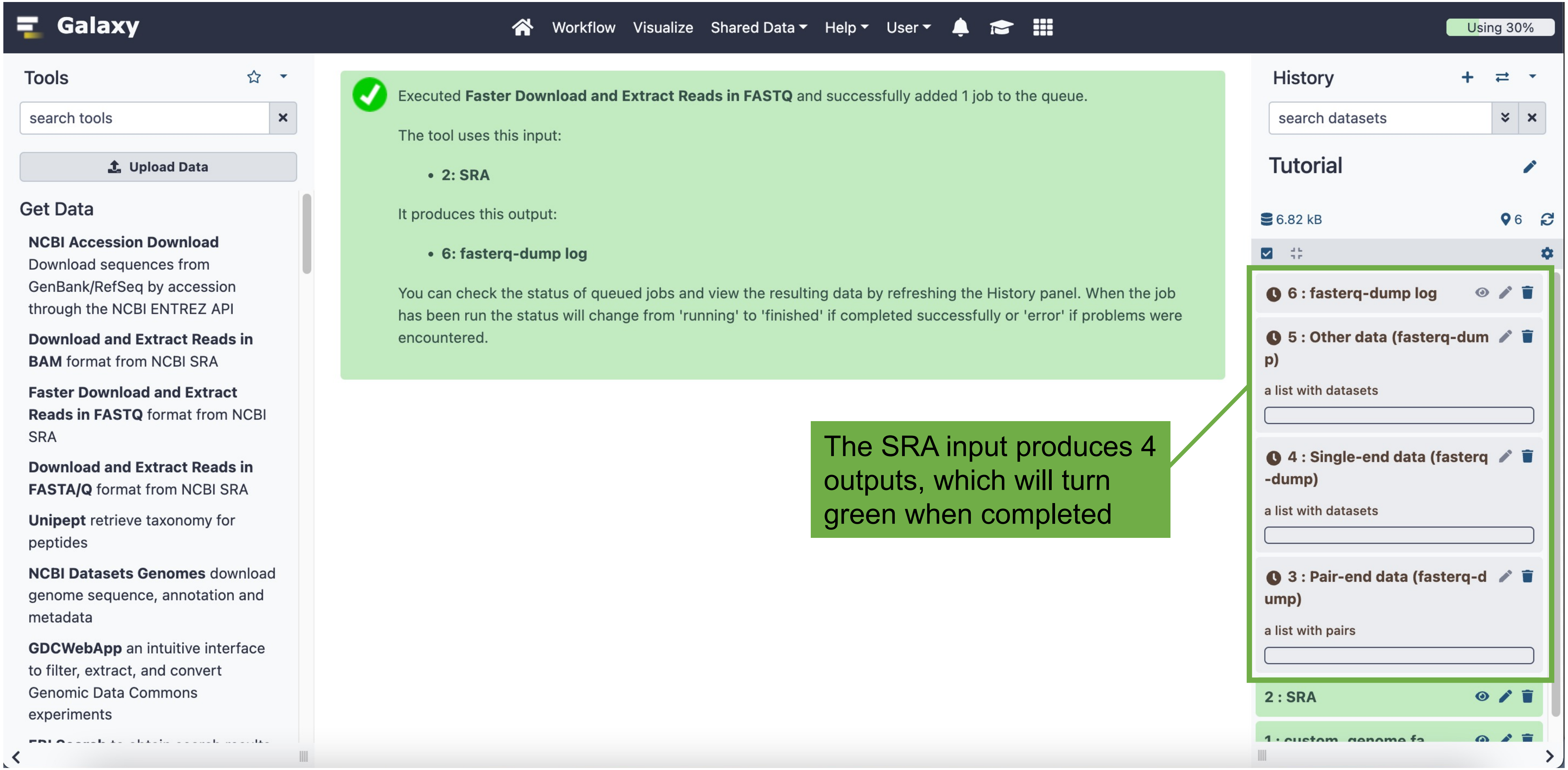

Galaxy
Workflow
Visualize
Shared Data
Help
User
Using 30%
Tools
search tools
Upload Data
Get Data
NCBI Accession Download Download sequences from GenBank/RefSeq by accession through the NCBI ENTREZ API
Download and Extract Reads in BAM format from NCBI SRA
Faster Download and Extract Reads in FASTQ format from NCBI SRA
Download and Extract Reads in FASTA/Q format from NCBI SRA
Unipept retrieve taxonomy for peptides
NCBI Datasets Genomes download genome sequence, annotation and metadata
GDCWebApp an intuitive interface to filter, extract, and convert Genomic Data Commons experiments
Executed Faster Download and Extract Reads in FASTQ and successfully added 1 job to the queue.
The tool uses this input:
2: SRA
It produces this output:
6: fasterq-dump log
You can check the status of queued jobs and view the resulting data by refreshing the History panel. When the job has been run the status will change from 'running' to 'finished' if completed successfully or 'error' if problems were encountered.
The SRA input produces 4 outputs, which will turn green when completed
History
search datasets
Tutorial
6.82 kB
6
6 : fasterq-dump log
5 : Other data (fasterq-dump)
a list with datasets
4 : Single-end data (fasterq-dump)
a list with datasets
3 : Pair-end data (fasterq-dump)
a list with pairs
2 : SRA

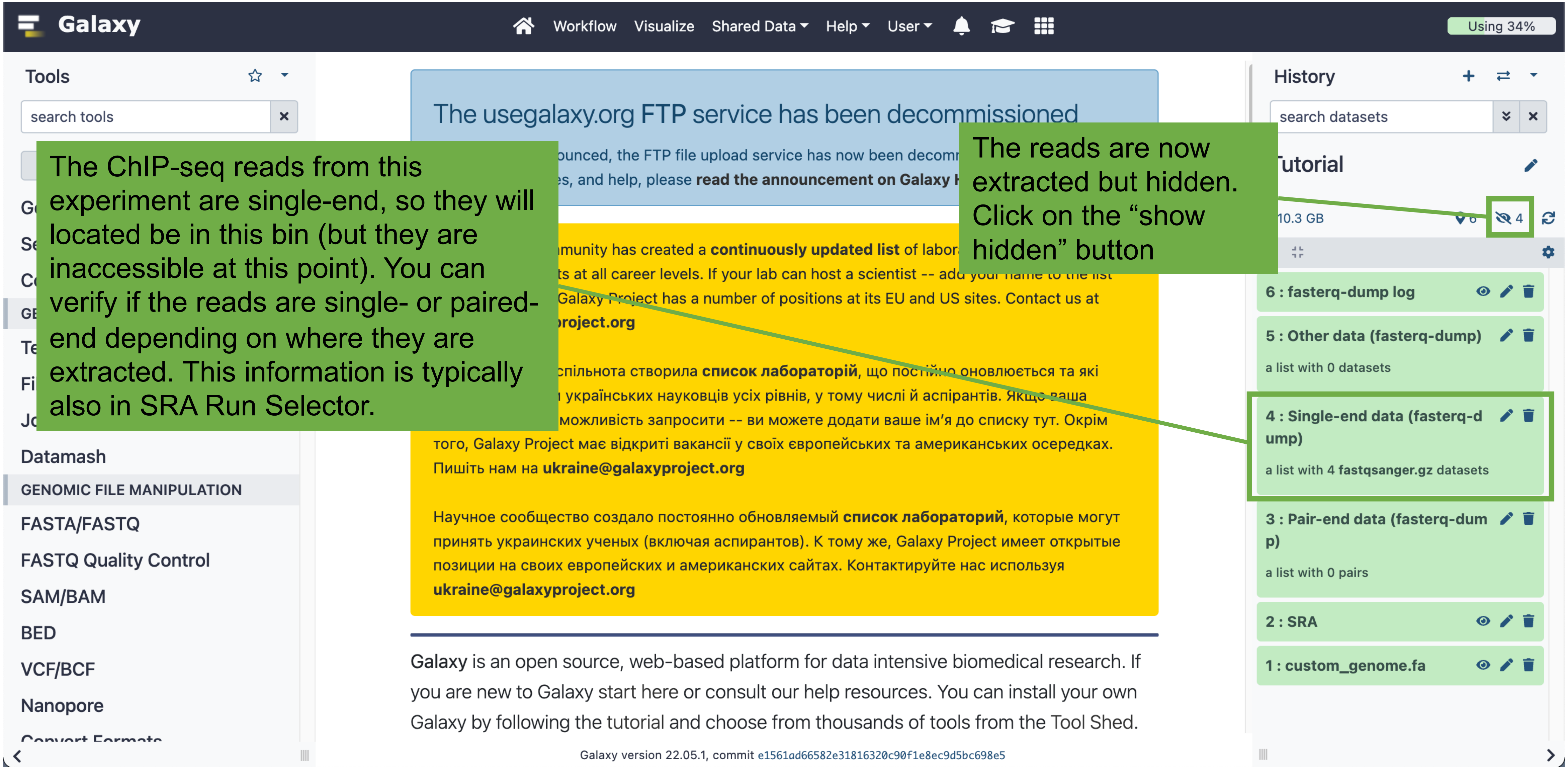

The ChIP-seq reads from this experiment are single-end, so they will located be in this bin (but they are inaccessible at this point). You can verify if the reads are single- or paired-end depending on where they are extracted. This information is typically also in SRA Run Selector.
The reads are now extracted but hidden. Click on the “show hidden” button
Galaxy
Workflow
Visualize
Shared Data
Help
User
Using 34%
Tools
search tools
Datamash
GENOMIC FILE MANIPULATION
FASTA/FASTQ
FASTQ Quality Control
SAM/BAM
BED
VCF/BCF
Nanopore
The usegalaxy.org FTP service has been decommissioned
History
search datasets
10.3 GB
6 : fasterq-dump log
5 : Other data (fasterq-dump)
a list with 0 datasets
4 : Single-end data (fasterq-dump)
a list with 4 fastqsanger.gz datasets
3 : Pair-end data (fasterq-dump)
a list with 0 pairs
2 : SRA
1 : custom_genome.fa
Galaxy is an open source, web-based platform for data intensive biomedical research. If you are new to Galaxy start here or consult our help resources. You can install your own Galaxy by following the tutorial and choose from thousands of tools from the Tool Shed.
Galaxy version 22.05.1, commit e1561ad66582e31816320c90f1e8ec9d5bc698e5

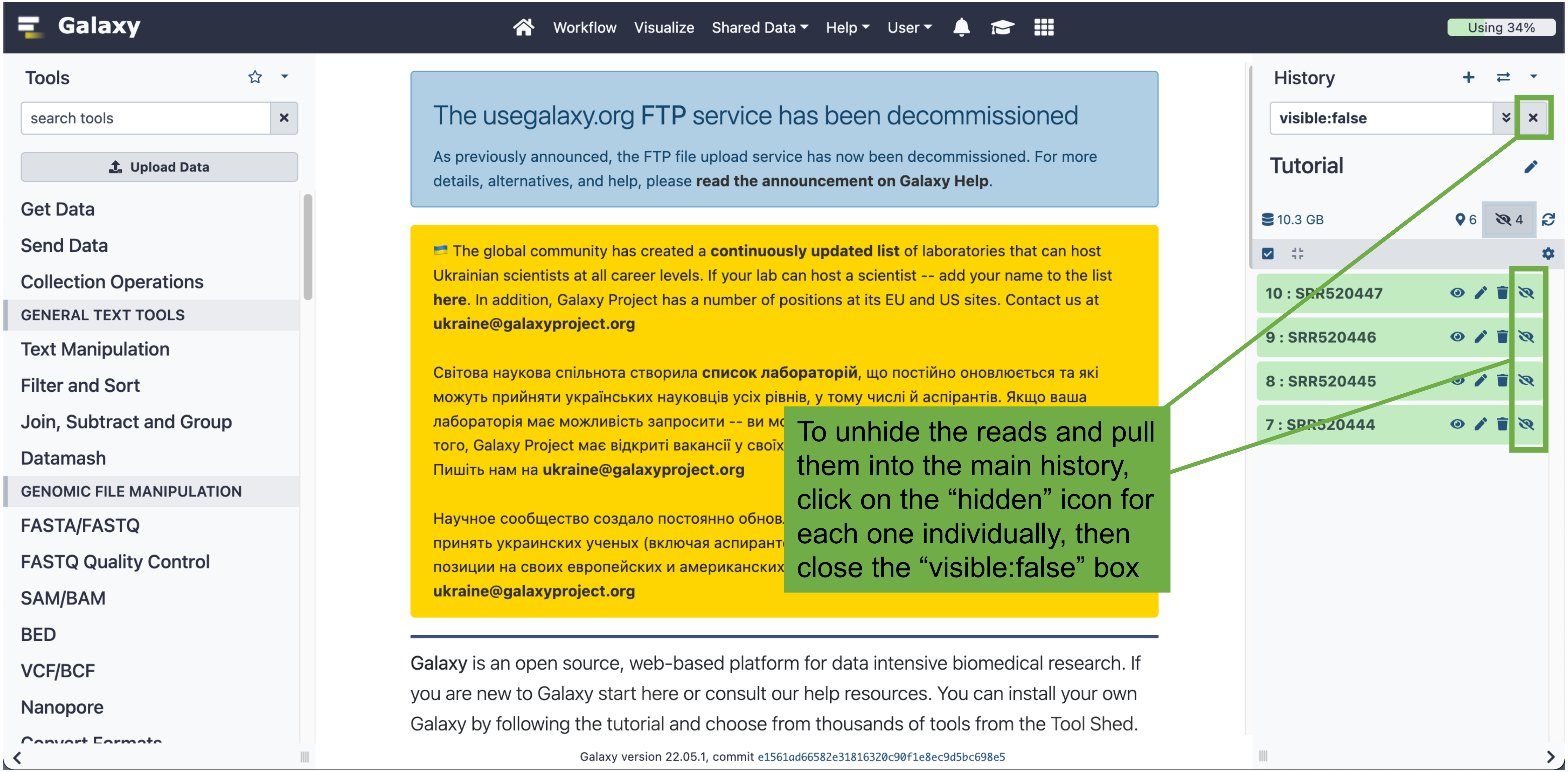
Galaxy
Workflow
Visualize
Shared Data
Help
User
Using 34%
Tools
search tools
Upload Data
Get Data
Send Data
Collection Operations
GENERAL TEXT TOOLS
Text Manipulation
Filter and Sort
Join, Subtract and Group
Datamash
GENOMIC FILE MANIPULATION
FASTA/FASTQ
FASTQ Quality Control
SAM/BAM
BED
VCF/BCF
Nanopore
The usegalaxy.org FTP service has been decommissioned
As previously announced, the FTP file upload service has now been decommissioned. For more details, alternatives, and help, please read the announcement on Galaxy Help.
The global community has created a continuously updated list of laboratories that can host Ukrainian scientists at all career levels. If your lab can host a scientist -- add your name to the list here. In addition, Galaxy Project has a number of positions at its EU and US sites. Contact us at
Світова наукова спільнота створила список лабораторій, що постійно оновлюється та які можуть прийняти українських науковців усіх рівнів, у тому числі й аспірантів. Якщо ваша лабораторія має можливість запросити -- ви мо
того, Galaxy Project має відкриті вакансії у своїх
Пишіть нам на
Научное сообщество создало постоянно обнов
принять украинских ученых (включая аспиранто
позиции на своих европейских и американских

Galaxy is an open source, web-based platform for data intensive biomedical research. If you are new to Galaxy start here or consult our help resources. You can install your own Galaxy by following the tutorial and choose from thousands of tools from the Tool Shed.
Galaxy version 22.05.1, commit e1561ad66582e31816320c90f1e8ec9d5bc698e5
History
visible:false
Tutorial
10.3 GB
6
4
10 : SRR520447
9 : SRR520446
8 : SRR520445
7 : SRR520444
To unhide the reads and pull them into the main history, click on the “hidden” icon for each one individually, then close the “visible:false” box

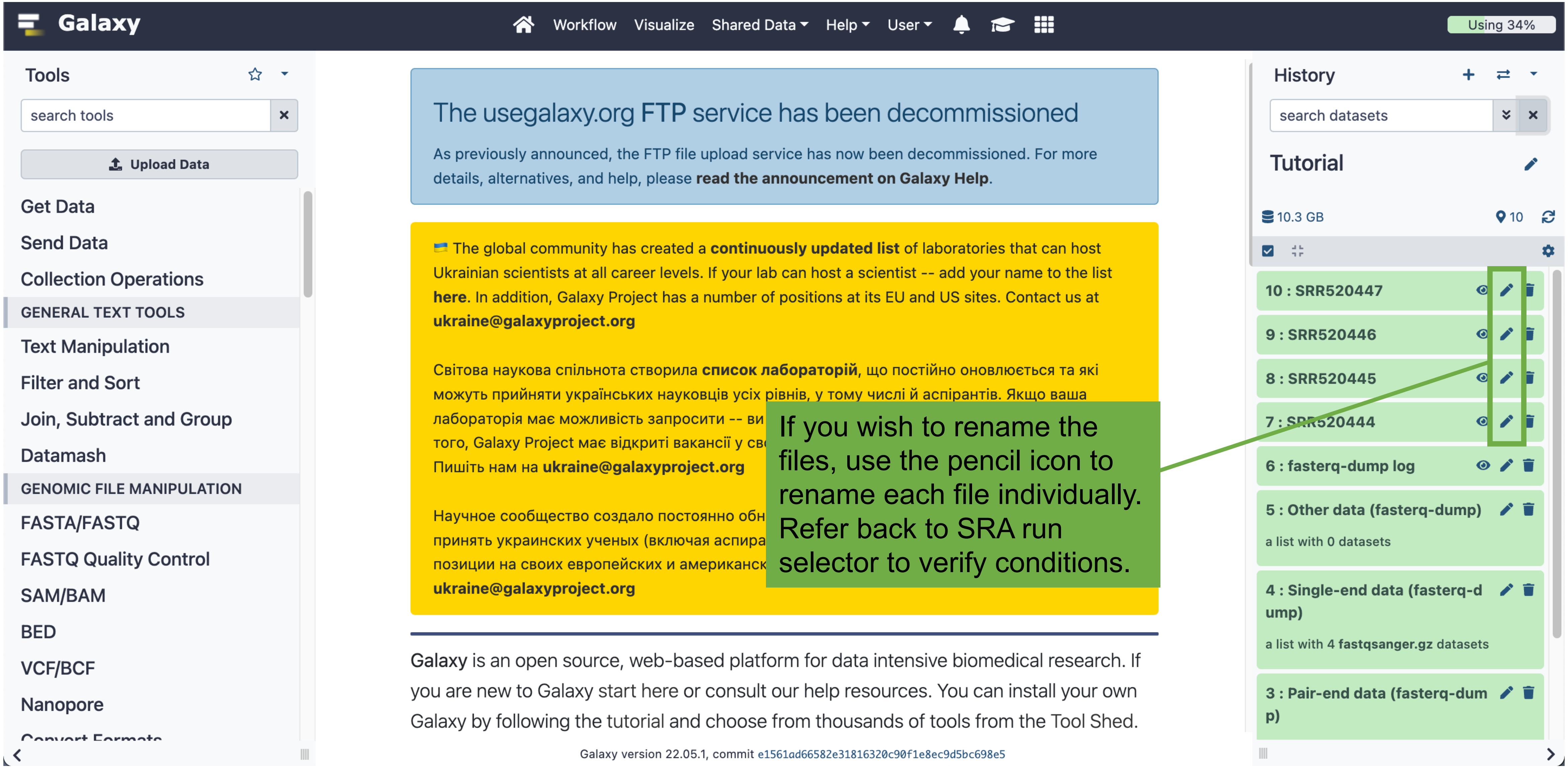

Galaxy
Workflow
Visualize
Shared Data
Help
User
Using 34%
Tools
search tools
Upload Data
Get Data
Send Data
Collection Operations
GENERAL TEXT TOOLS
Text Manipulation
Filter and Sort
Join, Subtract and Group
Datamash
GENOMIC FILE MANIPULATION
FASTA/FASTQ
FASTQ Quality Control
SAM/BAM
BED
VCF/BCF
Nanopore
The usegalaxy.org FTP service has been decommissioned
As previously announced, the FTP file upload service has now been decommissioned. For more details, alternatives, and help, please read the announcement on Galaxy Help.
The global community has created a continuously updated list of laboratories that can host Ukrainian scientists at all career levels. If your lab can host a scientist -- add your name to the list here. In addition, Galaxy Project has a number of positions at its EU and US sites. Contact us at
Світова наукова спільнота створила список лабораторій, що постійно оновлюється та які можуть прийняти українських науковців усіх рівнів, у тому числі й аспірантів. Якщо ваша лабораторія має можливість запросити -- ви
того, Galaxy Project має відкриті вакансії у св
Пишіть нам на
Научное сообщество создало постоянно обн
принять украинских ученых (включая аспира
позиции на своих европейских и американск

If you wish to rename the files, use the pencil icon to rename each file individually. Refer back to SRA run selector to verify conditions.
Galaxy is an open source, web-based platform for data intensive biomedical research. If you are new to Galaxy start here or consult our help resources. You can install your own Galaxy by following the tutorial and choose from thousands of tools from the Tool Shed.
Galaxy version 22.05.1, commit e1561ad66582e31816320c90f1e8ec9d5bc698e5
History
search datasets
Tutorial
10.3 GB
10
10 : SRR520447
9 : SRR520446
8 : SRR520445
7 : SRR520444
6 : fasterq-dump log
5 : Other data (fasterq-dump)
a list with 0 datasets
4 : Single-end data (fasterq-dump)
a list with 4 fastqsanger.gz datasets
3 : Pair-end data (fasterq-dump)

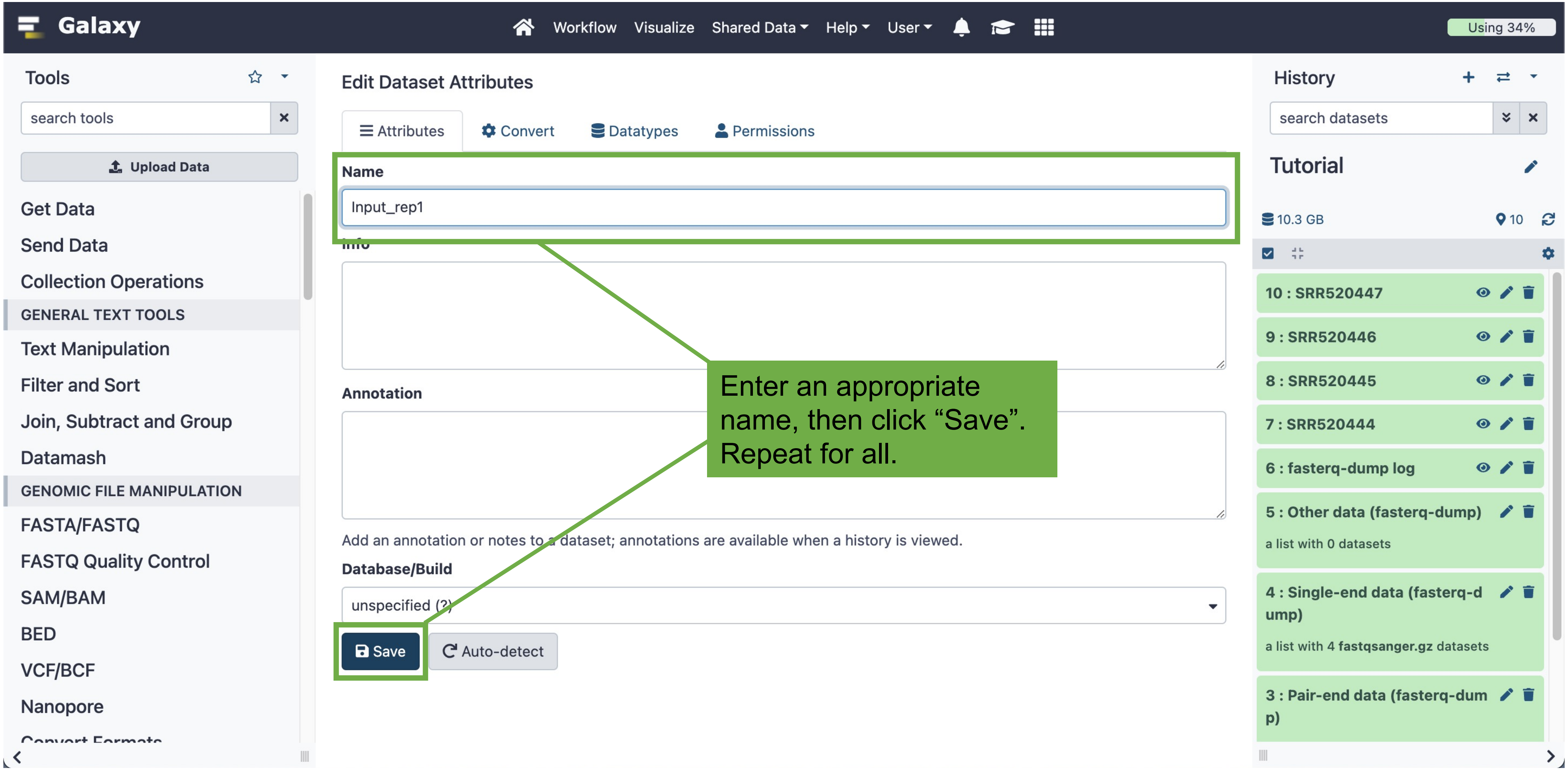
Galaxy
Workflow
Visualize
Shared Data
Help
User
Using 34%
Tools
search tools
Upload Data
Get Data
Send Data
Collection Operations
GENERAL TEXT TOOLS
Text Manipulation
Filter and Sort
Join, Subtract and Group
Datamash
GENOMIC FILE MANIPULATION
FASTA/FASTQ
FASTQ Quality Control
SAM/BAM
BED
VCF/BCF
Nanopore
Edit Dataset Attributes
Attributes
Convert
Datatypes
Permissions
Name
Input_rep1
Annotation
Add an annotation or notes to a dataset; annotations are available when a history is viewed.
Database/Build
unspecified (?)
Save
Auto-detect
Enter an appropriate name, then click “Save”. Repeat for all.
History
search datasets
Tutorial
10.3 GB
10
10 : SRR520447
9 : SRR520446
8 : SRR520445
7 : SRR520444
6 : fasterq-dump log
5 : Other data (fasterq-dump)
a list with 0 datasets
4 : Single-end data (fasterq-dump)
a list with 4 fastqsanger.gz datasets
3 : Pair-end data (fasterq-dump)

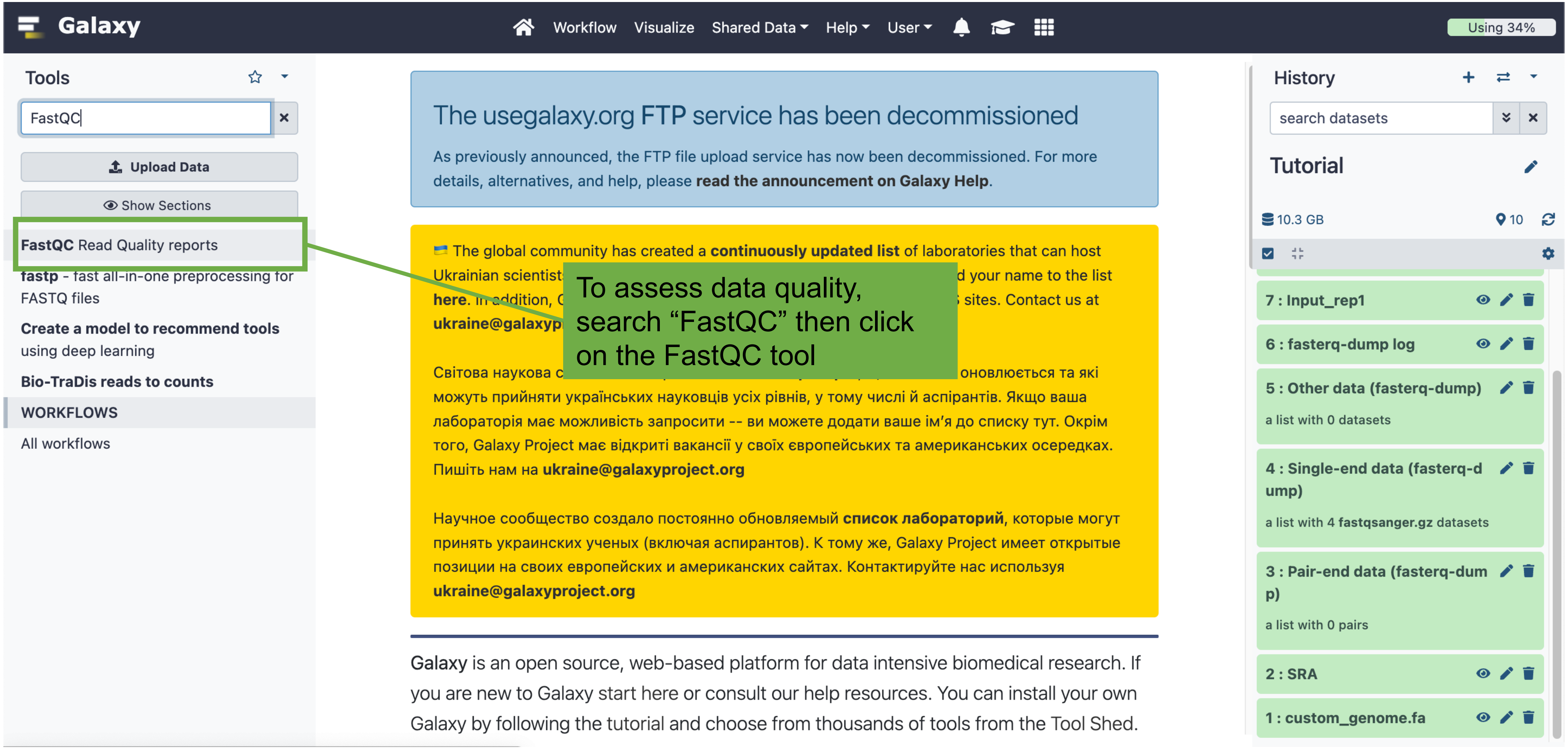
Galaxy
Workflow
Visualize
Shared Data
Help
User
Using 34%
Tools
FastQC
Upload Data
Show Sections
FastQC Read Quality reports
FASTQ files
Create a model to recommend tools
using deep learning
Bio-TraDis reads to counts
WORKFLOWS
All workflows
The usegalaxy.org FTP service has been decommissioned
As previously announced, the FTP file upload service has now been decommissioned. For more details, alternatives, and help, please read the announcement on Galaxy Help.
The global community has created a continuously updated list of laboratories that can host
можуть прийняти українських науковців усіх рівнів, у тому числі й аспірантів. Якщо ваша
лабораторія має можливість запросити -- ви можете додати ваше ім'я до списку тут. Окрім
того, Galaxy Project має відкриті вакансії у своїх європейських та американських осередках.
Пишіть нам на
Научное сообщество создало постоянно обновляемый список лабораторий, которые могут
принять украинских ученых (включая аспирантов). К тому же, Galaxy Project имеет открытые
позиции на своих европейских и американских сайтах. Контактируйте нас используя

Galaxy is an open source, web-based platform for data intensive biomedical research. If you are new to Galaxy start here or consult our help resources. You can install your own Galaxy by following the tutorial and choose from thousands of tools from the Tool Shed.
To assess data quality, search "FastQC" then click on the FastQC tool
History
search datasets
Tutorial
10.3 GB
10
7 : Input_rep1
6 : fasterq-dump log
5 : Other data (fasterq-dump)
a list with 0 datasets
4 : Single-end data (fasterq-dump)
a list with 4 fastqsanger.gz datasets
3 : Pair-end data (fasterq-dump)
a list with 0 pairs
2 : SRA
1 : custom_genome.fa

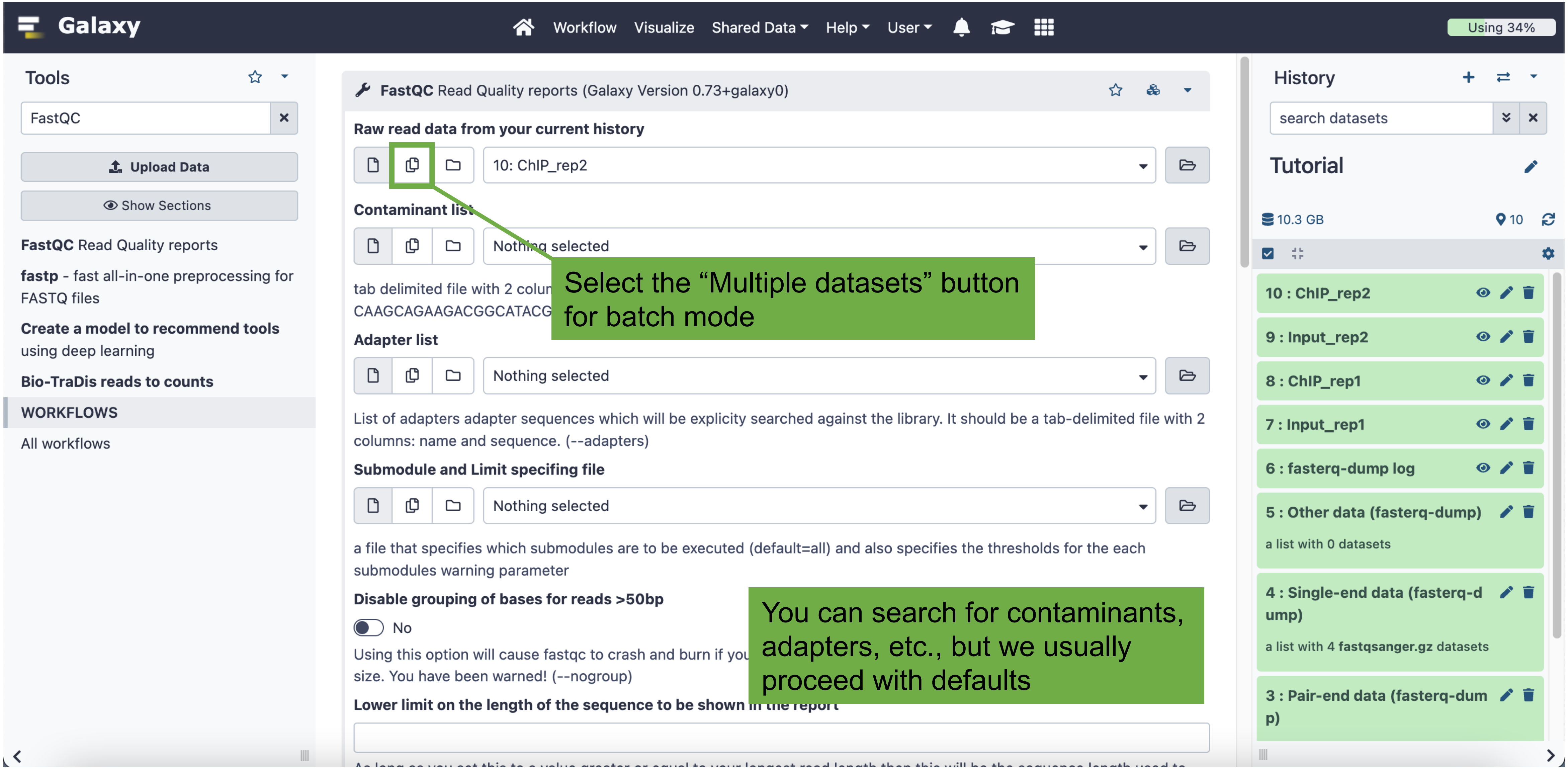
Galaxy
Workflow
Visualize
Shared Data
Help
User
Using 34%
Tools
FastQC
Upload Data
Show Sections
FastQC Read Quality reports
fastp - fast all-in-one preprocessing for FASTQ files
Create a model to recommend tools using deep learning
Bio-TraDis reads to counts
WORKFLOWS
All workflows
FastQC Read Quality reports (Galaxy Version 0.73+galaxy0)
Raw read data from your current history
10: ChIP_rep2
Contaminant list
Nothing selected
Select the "Multiple datasets" button for batch mode
Adapter list
Nothing selected
List of adapters adapter sequences which will be explicity searched against the library. It should be a tab-delimited file with 2 columns: name and sequence. (--adapters)
Submodule and Limit specifing file
Nothing selected
a file that specifies which submodules are to be executed (default=all) and also specifies the thresholds for the each submodules warning parameter
Disable grouping of bases for reads >50bp
No
You can search for contaminants, adapters, etc., but we usually proceed with defaults
History
search datasets
Tutorial
10.3 GB
10
10 : ChIP_rep2
9 : Input_rep2
8 : ChIP_rep1
7 : Input_rep1
6 : fasterq-dump log
5 : Other data (fasterq-dump)
a list with 0 datasets
4 : Single-end data (fasterq-dump)
a list with 4 fastqsanger.gz datasets
3 : Pair-end data (fasterq-dump)

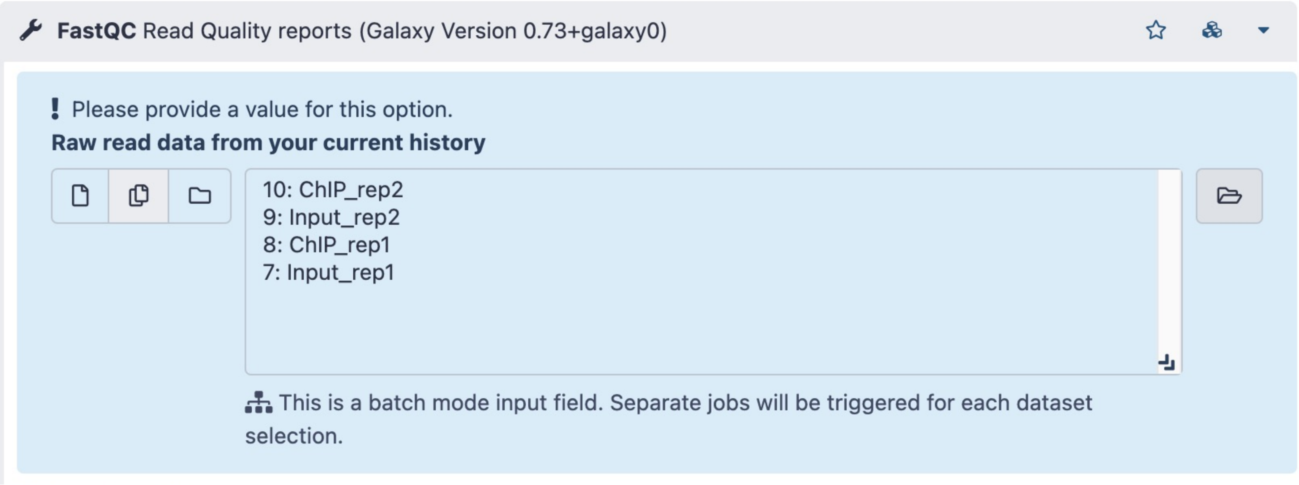


Hold “command” (Mac) or “control” (PC) and select all datasets you wish to analyze

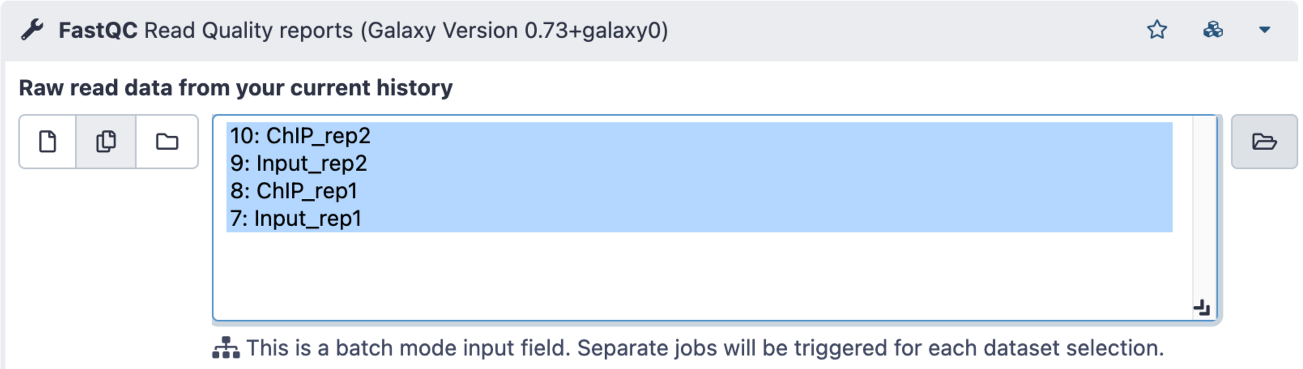


Then click “Execute” at the bottom of the page

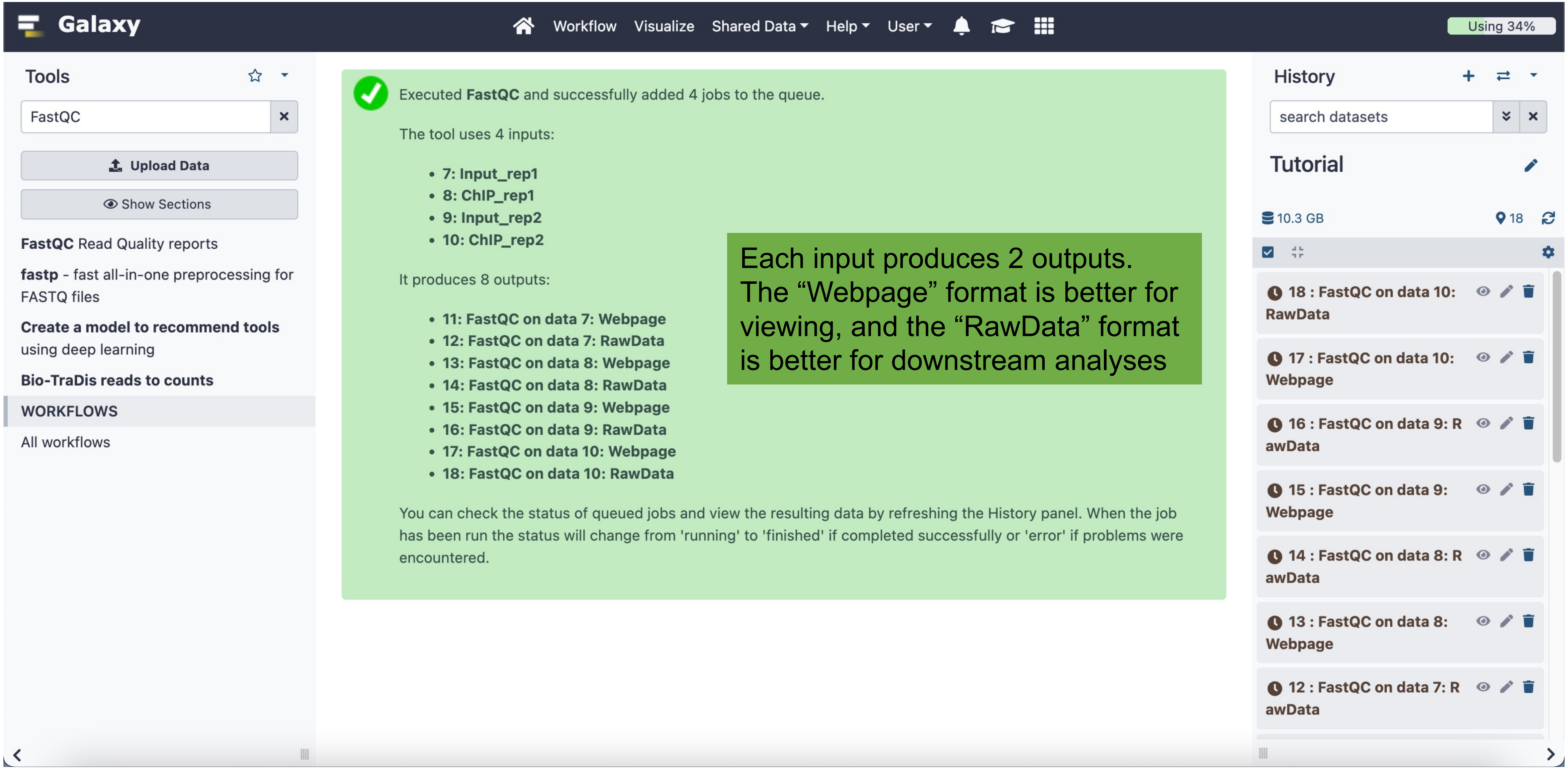
Galaxy
Workflow
Visualize
Shared Data
Help
User
Using 34%
Tools
FastQC
Upload Data
Show Sections
FastQC Read Quality reports
fastp - fast all-in-one preprocessing for FASTQ files
Create a model to recommend tools using deep learning
Bio-TraDis reads to counts
WORKFLOWS
All workflows
Executed FastQC and successfully added 4 jobs to the queue.
The tool uses 4 inputs:
7: Input_rep1
8: ChIP_rep1
9: Input_rep2
10: ChIP_rep2
It produces 8 outputs:
11: FastQC on data 7: Webpage
12: FastQC on data 7: RawData
13: FastQC on data 8: Webpage
14: FastQC on data 8: RawData
15: FastQC on data 9: Webpage
16: FastQC on data 9: RawData
17: FastQC on data 10: Webpage
18: FastQC on data 10: RawData
You can check the status of queued jobs and view the resulting data by refreshing the History panel. When the job has been run the status will change from 'running' to 'finished' if completed successfully or 'error' if problems were encountered.
Each input produces 2 outputs. The “Webpage” format is better for viewing, and the “RawData” format is better for downstream analyses
History
search datasets
Tutorial
10.3 GB
18
18 : FastQC on data 10: RawData
17 : FastQC on data 10: Webpage
16 : FastQC on data 9: RawData
15 : FastQC on data 9: Webpage
14 : FastQC on data 8: RawData
13 : FastQC on data 8: Webpage
12 : FastQC on data 7: RawData

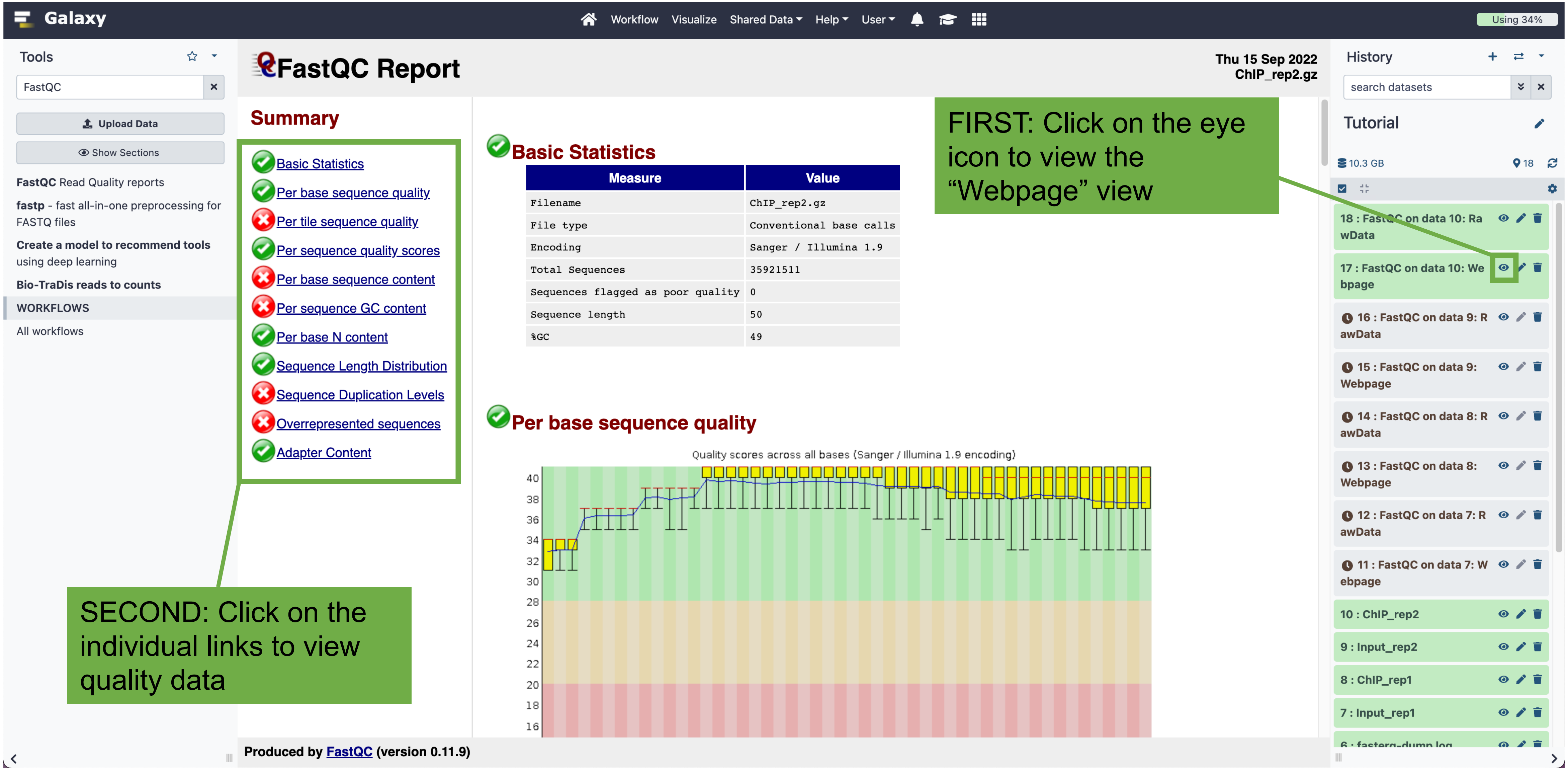
Galaxy
Workflow
Visualize
Shared Data
Help
User
Using 34%
Tools
FastQC
Upload Data
Show Sections
FastQC Read Quality reports
fastp - fast all-in-one preprocessing for FASTQ files
Create a model to recommend tools using deep learning
Bio-TraDis reads to counts
WORKFLOWS
All workflows
FastQC Report
Thu 15 Sep 2022
ChIP_rep2.gz
Summary
Basic Statistics
Per base sequence quality
Per tile sequence quality
Per sequence quality scores
Per base sequence content
Per sequence GC content
Per base N content
Sequence Length Distribution
Sequence Duplication Levels
Overrepresented sequences
Adapter Content
Basic Statistics
Measure | Value
Filename | ChIP_rep2.gz
File type | Conventional base calls
Encoding | Sanger / Illumina 1.9
Total Sequences | 35921511
Sequences flagged as poor quality | 0
Sequence length | 50
%GC | 49
Per base sequence quality
Quality scores across all bases (Sanger / Illumina 1.9 encoding)
40
38
36
34
32
30
28
26
24
22
20
18
16
Produced by FastQC (version 0.11.9)
History
search datasets
Tutorial
10.3 GB
18
18 : FastQC on data 10: RawData
17 : FastQC on data 10: Webpage
16 : FastQC on data 9: RawData
15 : FastQC on data 9: Webpage
14 : FastQC on data 8: RawData
13 : FastQC on data 8: Webpage
12 : FastQC on data 7: RawData
11 : FastQC on data 7: Webpage
10 : ChIP_rep2
9 : Input_rep2
8 : ChIP_rep1
7 : Input_rep1
6 : fasterq-dump log
FIRST: Click on the eye icon to view the “Webpage” view
SECOND: Click on the individual links to view quality data

### Aligning the ChIP-seq reads

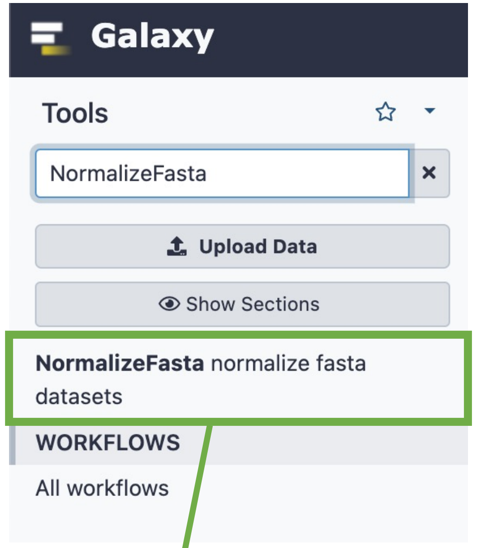


Click on the “NormalizeFasta” tool

If you are using a custom genome (that you imported as a .fasta or .fa file), you will need to “normalize” it so that it can be used with downstream tools. Skip these steps if you are using a built-in genome.

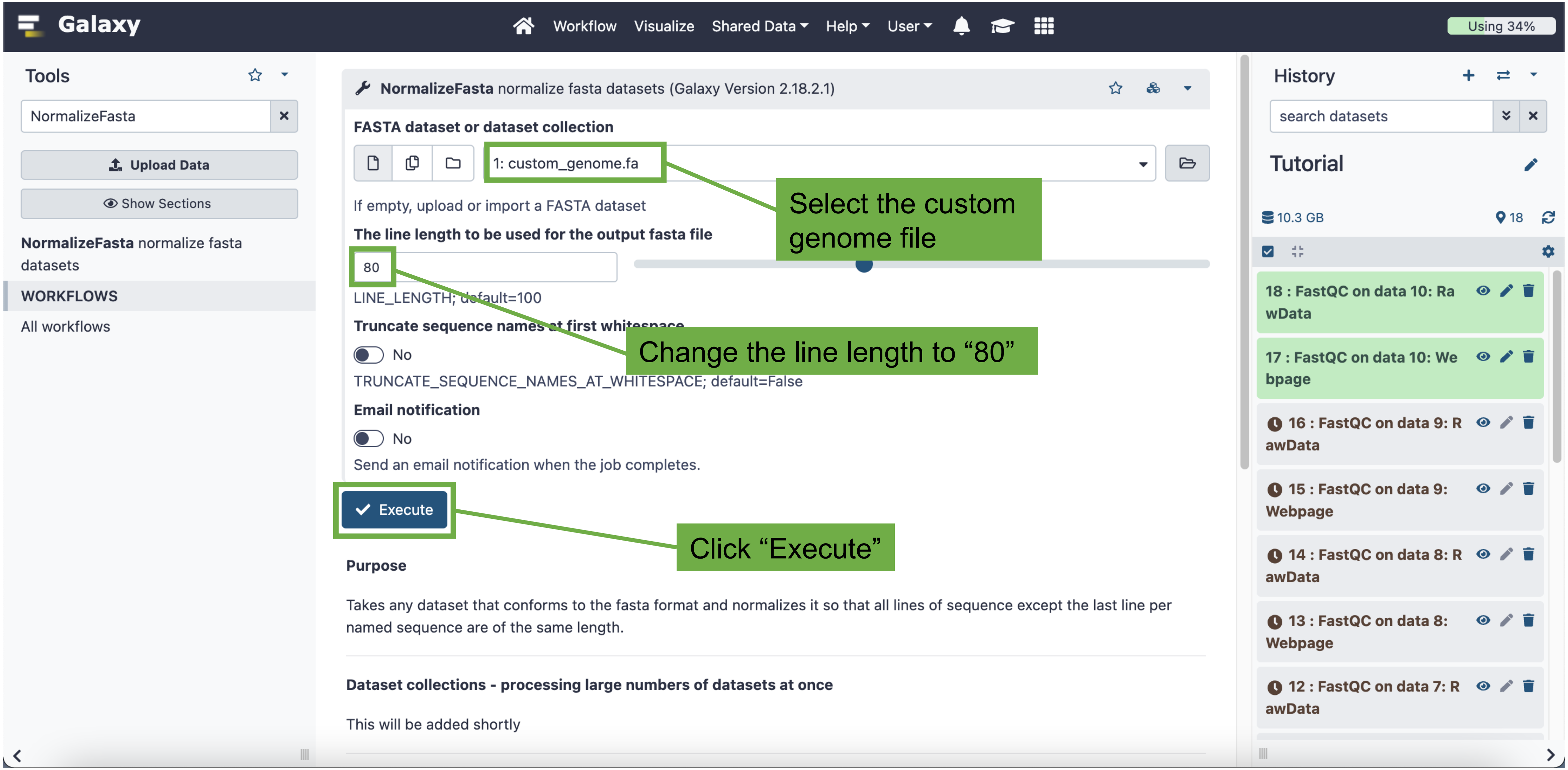
Galaxy
Workflow
Visualize
Shared Data
Help
User
Using 34%
Tools
NormalizeFasta
Upload Data
Show Sections
NormalizeFasta normalize fasta datasets
WORKFLOWS
All workflows
NormalizeFasta normalize fasta datasets (Galaxy Version 2.18.2.1)
FASTA dataset or dataset collection
1: custom_genome.fa
If empty, upload or import a FASTA dataset
The line length to be used for the output fasta file
80
LINE_LENGTH; default=100
Truncate sequence names at first whitespace
No
TRUNCATE_SEQUENCE_NAMES_AT_WHITESPACE; default=False
Email notification
No
Send an email notification when the job completes.
Execute
Purpose
Takes any dataset that conforms to the fasta format and normalizes it so that all lines of sequence except the last line per named sequence are of the same length.
Dataset collections - processing large numbers of datasets at once
This will be added shortly
Select the custom genome file
Change the line length to “80”
Click “Execute”
History
search datasets
Tutorial
10.3 GB
18
18 : FastQC on data 10: RawData
17 : FastQC on data 10: Webpage
16 : FastQC on data 9: RawData
15 : FastQC on data 9: Webpage
14 : FastQC on data 8: RawData
13 : FastQC on data 8: Webpage
12 : FastQC on data 7: RawData

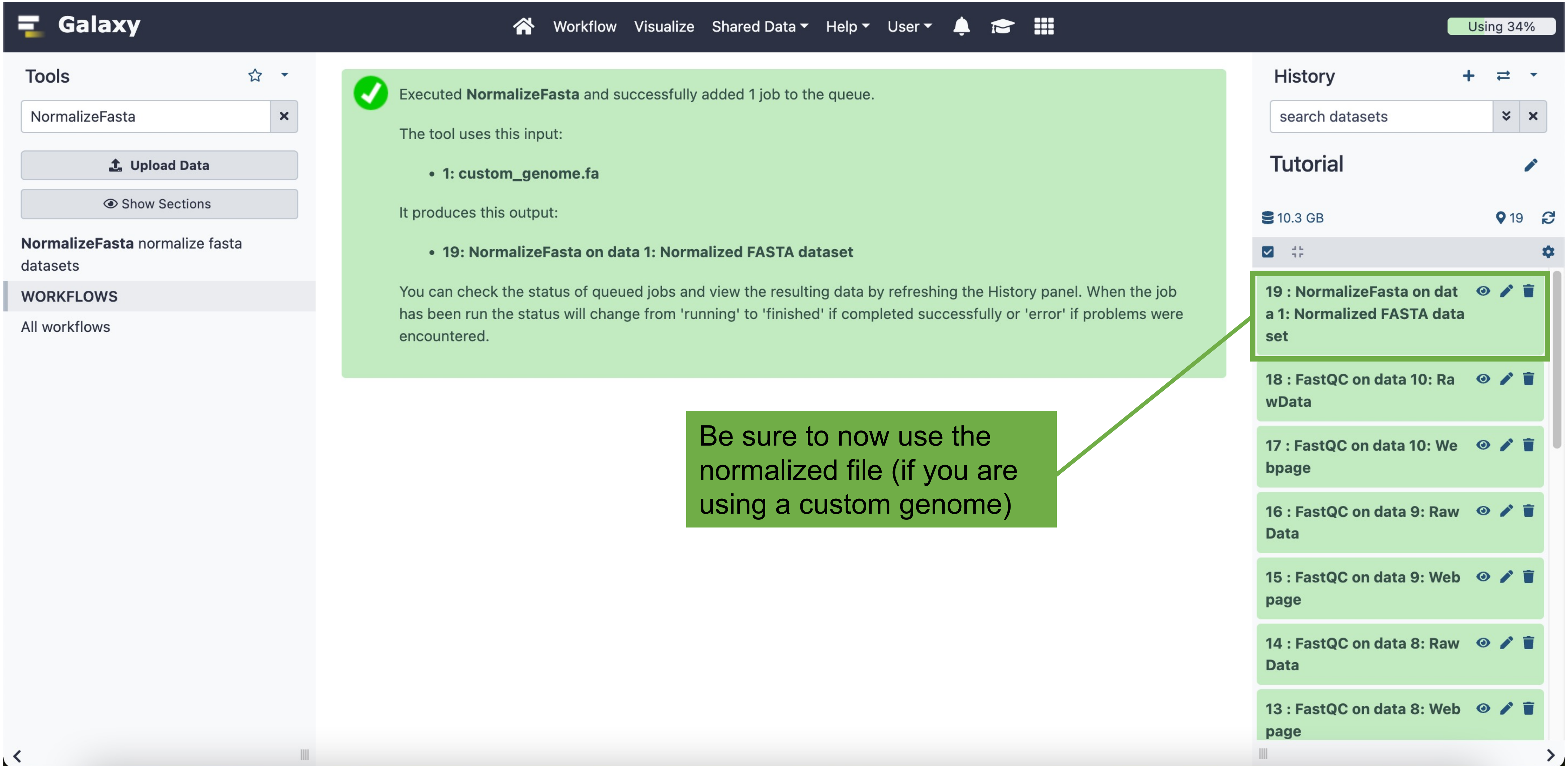
Galaxy
Workflow
Visualize
Shared Data
Help
User
Using 34%
Tools
NormalizeFasta
Upload Data
Show Sections
NormalizeFasta normalize fasta datasets
WORKFLOWS
All workflows
Executed NormalizeFasta and successfully added 1 job to the queue.
The tool uses this input:
• 1: custom_genome.fa
It produces this output:
• 19: NormalizeFasta on data 1: Normalized FASTA dataset
You can check the status of queued jobs and view the resulting data by refreshing the History panel. When the job has been run the status will change from 'running' to 'finished' if completed successfully or 'error' if problems were encountered.
Be sure to now use the normalized file (if you are using a custom genome)
History
search datasets
Tutorial
10.3 GB
19
19 : NormalizeFasta on data 1: Normalized FASTA dataset
18 : FastQC on data 10: RawData
17 : FastQC on data 10: Webpage
16 : FastQC on data 9: RawData
15 : FastQC on data 9: Webpage
14 : FastQC on data 8: RawData
13 : FastQC on data 8: Webpage

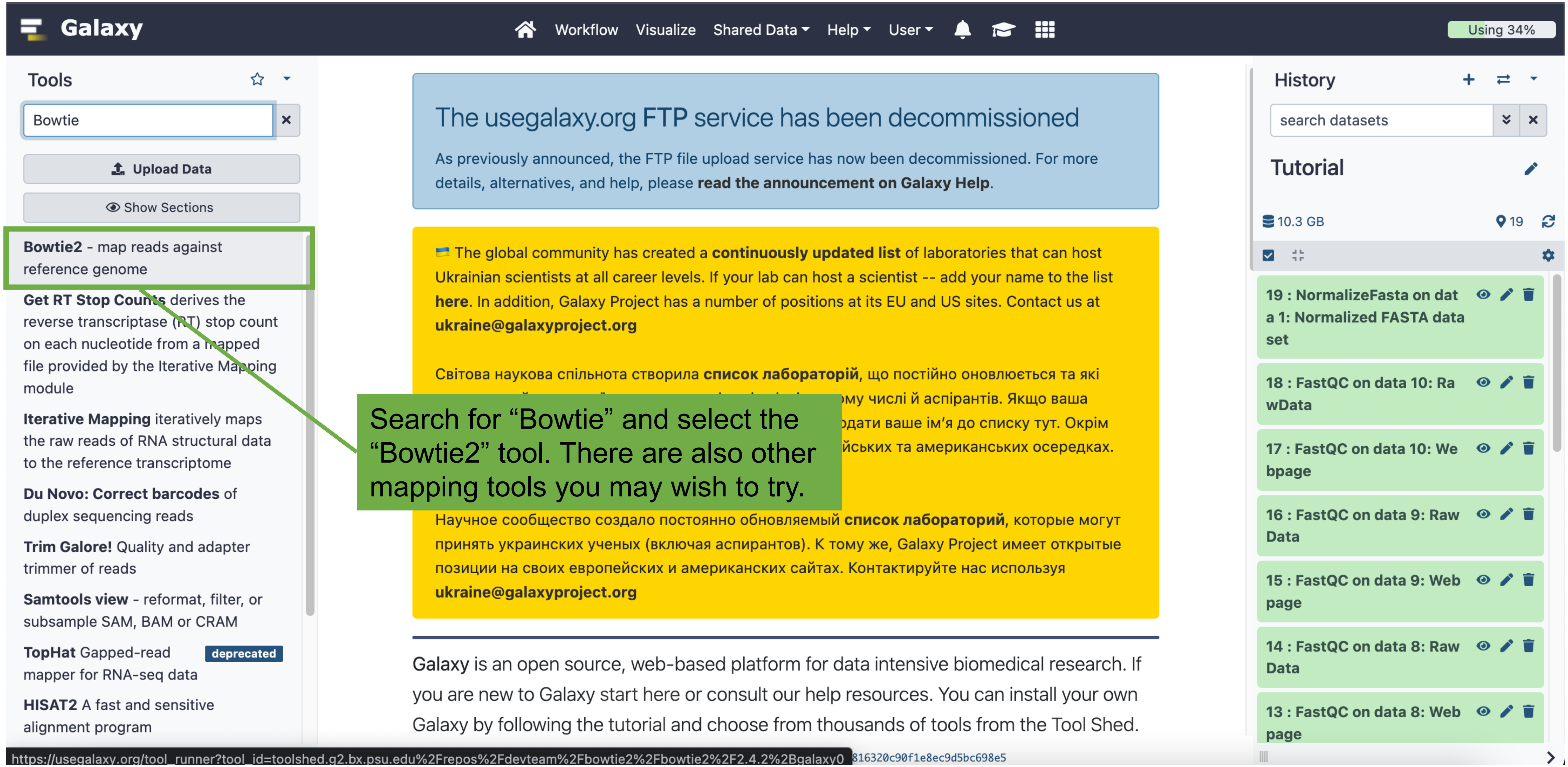
Search for “Bowtie” and select the “Bowtie2” tool. There are also other mapping tools you may wish to try.
Galaxy
Workflow
Visualize
Shared Data
Help
User
Using 34%
Tools
Bowtie
Upload Data
Show Sections
Bowtie2 - map reads against reference genome
Get RT Stop Counts derives the reverse transcriptase (RT) stop count on each nucleotide from a mapped file provided by the Iterative Mapping module
Iterative Mapping iteratively maps the raw reads of RNA structural data to the reference transcriptome
Du Novo: Correct barcodes of duplex sequencing reads
Trim Galore! Quality and adapter trimmer of reads
Samtools view - reformat, filter, or subsample SAM, BAM or CRAM
TopHat Gapped-read mapper for RNA-seq data
deprecated
HISAT2 A fast and sensitive alignment program
The usegalaxy.org FTP service has been decommissioned
As previously announced, the FTP file upload service has now been decommissioned. For more details, alternatives, and help, please read the announcement on Galaxy Help.
The global community has created a continuously updated list of laboratories that can host Ukrainian scientists at all career levels. If your lab can host a scientist -- add your name to the list here. In addition, Galaxy Project has a number of positions at its EU and US sites. Contact us at
Світова наукова спільнота створила список лабораторій, що постійно оновлюється та які
Научное сообщество создало постоянно обновляемый список лабораторий, которые могут принять украинских ученых (включая аспирантов). К тому же, Galaxy Project имеет открытые позиции на своих европейских и американских сайтах. Контактируйте нас используя
Galaxy is an open source, web-based platform for data intensive biomedical research. If you are new to Galaxy start here or consult our help resources. You can install your own Galaxy by following the tutorial and choose from thousands of tools from the Tool Shed.
History
search datasets
Tutorial
10.3 GB
19
19 : NormalizeFasta on data 1: Normalized FASTA dataset
18 : FastQC on data 10: RawData
17 : FastQC on data 10: Webpage
16 : FastQC on data 9: Raw Data
15 : FastQC on data 9: Web page
14 : FastQC on data 8: Raw Data
13 : FastQC on data 8: Web page
https://usegalaxy.org/tool_runner?tool_id=toolshed.g2.bx.psu.edu%2Frepos%2Fdevteam%2Fbowtie2%2Fbowtie2%2F2.4.2%2Bgalaxy0
816320c90f1e8ec9d5bc698e5

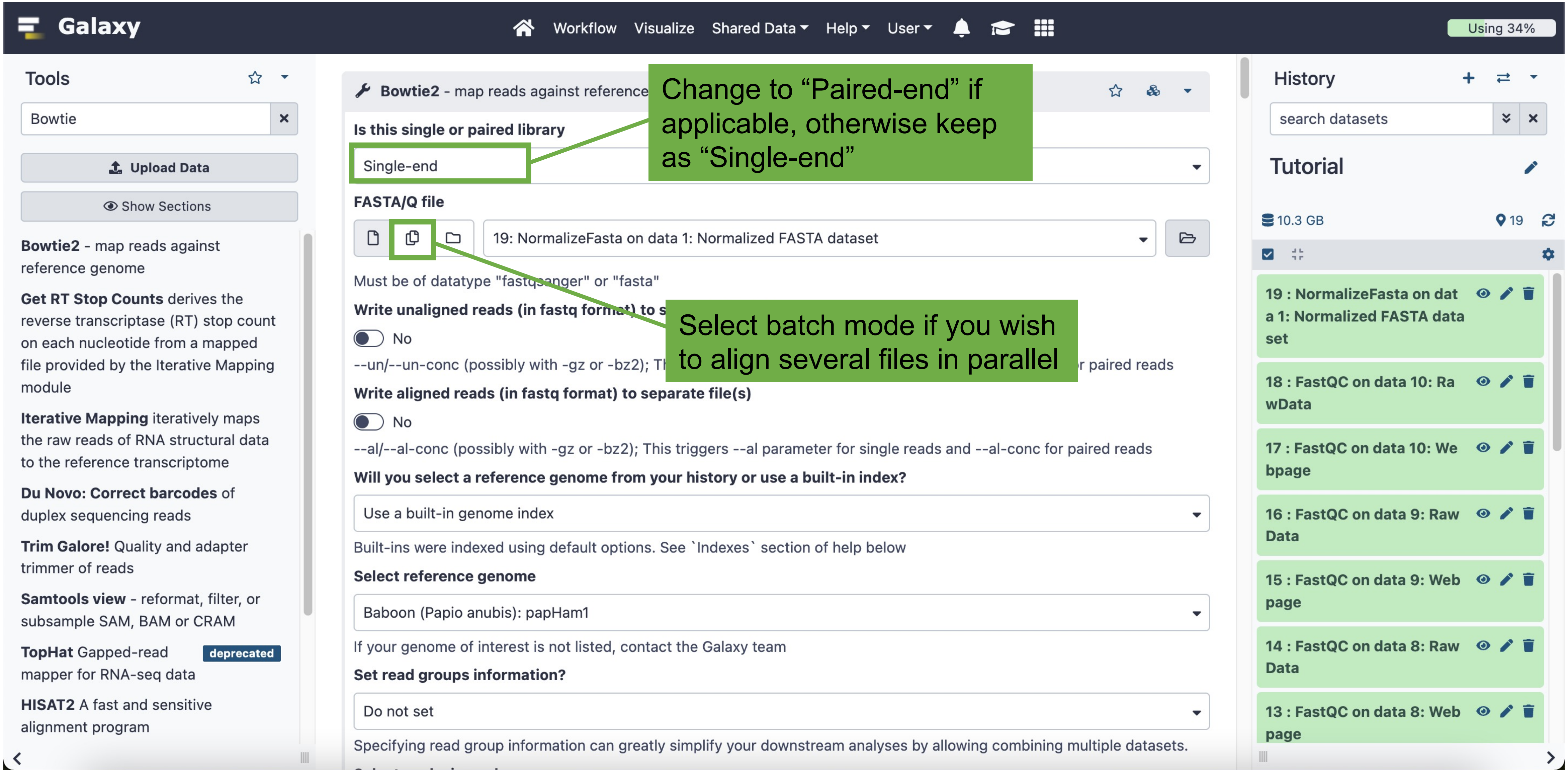

Galaxy
Workflow
Visualize
Shared Data
Help
User
Using 34%
Tools
Bowtie
Upload Data
Show Sections
Bowtie2 - map reads against reference genome
Get RT Stop Counts derives the reverse transcriptase (RT) stop count on each nucleotide from a mapped file provided by the Iterative Mapping module
Iterative Mapping iteratively maps the raw reads of RNA structural data to the reference transcriptome
Du Novo: Correct barcodes of duplex sequencing reads
Trim Galore! Quality and adapter trimmer of reads
Samtools view - reformat, filter, or subsample SAM, BAM or CRAM
TopHat Gapped-read mapper for RNA-seq data
deprecated
HISAT2 A fast and sensitive alignment program
Bowtie2 - map reads against reference
Is this single or paired library
Single-end
Change to “Paired-end” if applicable, otherwise keep as “Single-end”
FASTA/Q file
19: NormalizeFasta on data 1: Normalized FASTA dataset
Must be of datatype "fastqsanger" or "fasta"
Write unaligned reads (in fastq format) to s
No
--un/--un-conc (possibly with -gz or -bz2); Th
r paired reads
Select batch mode if you wish to align several files in parallel
Write aligned reads (in fastq format) to separate file(s)
No
--al/--al-conc (possibly with -gz or -bz2); This triggers --al parameter for single reads and --al-conc for paired reads
Will you select a reference genome from your history or use a built-in index?
Use a built-in genome index
Built-ins were indexed using default options. See `Indexes` section of help below
Select reference genome
Baboon (Papio anubis): papHam1
If your genome of interest is not listed, contact the Galaxy team
Set read groups information?
Do not set
Specifying read group information can greatly simplify your downstream analyses by allowing combining multiple datasets.
History
search datasets
Tutorial
10.3 GB
19
19 : NormalizeFasta on data 1: Normalized FASTA dataset
18 : FastQC on data 10: RawData
17 : FastQC on data 10: Webpage
16 : FastQC on data 9: Raw Data
15 : FastQC on data 9: Web page
14 : FastQC on data 8: Raw Data
13 : FastQC on data 8: Web page

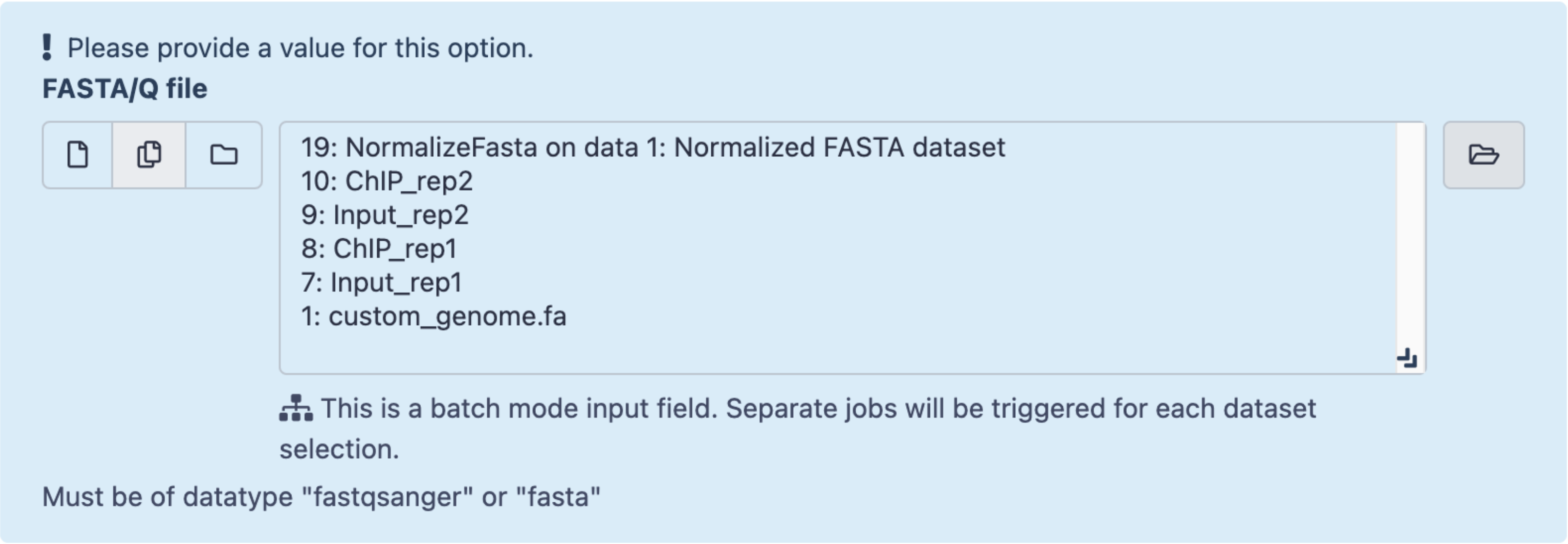


Hold “command” (Mac) or “control” (PC) and select all datasets you wish to analyze

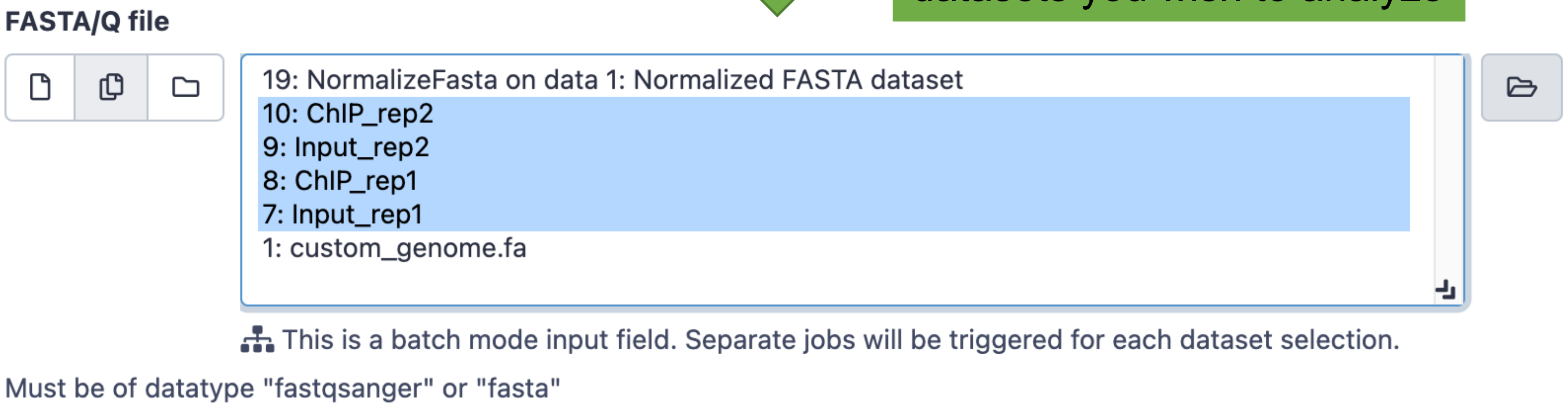


Note that we are only aligning the .fastq files (7-10) – I have not selected the custom genome files (original (1) and normalized (19))

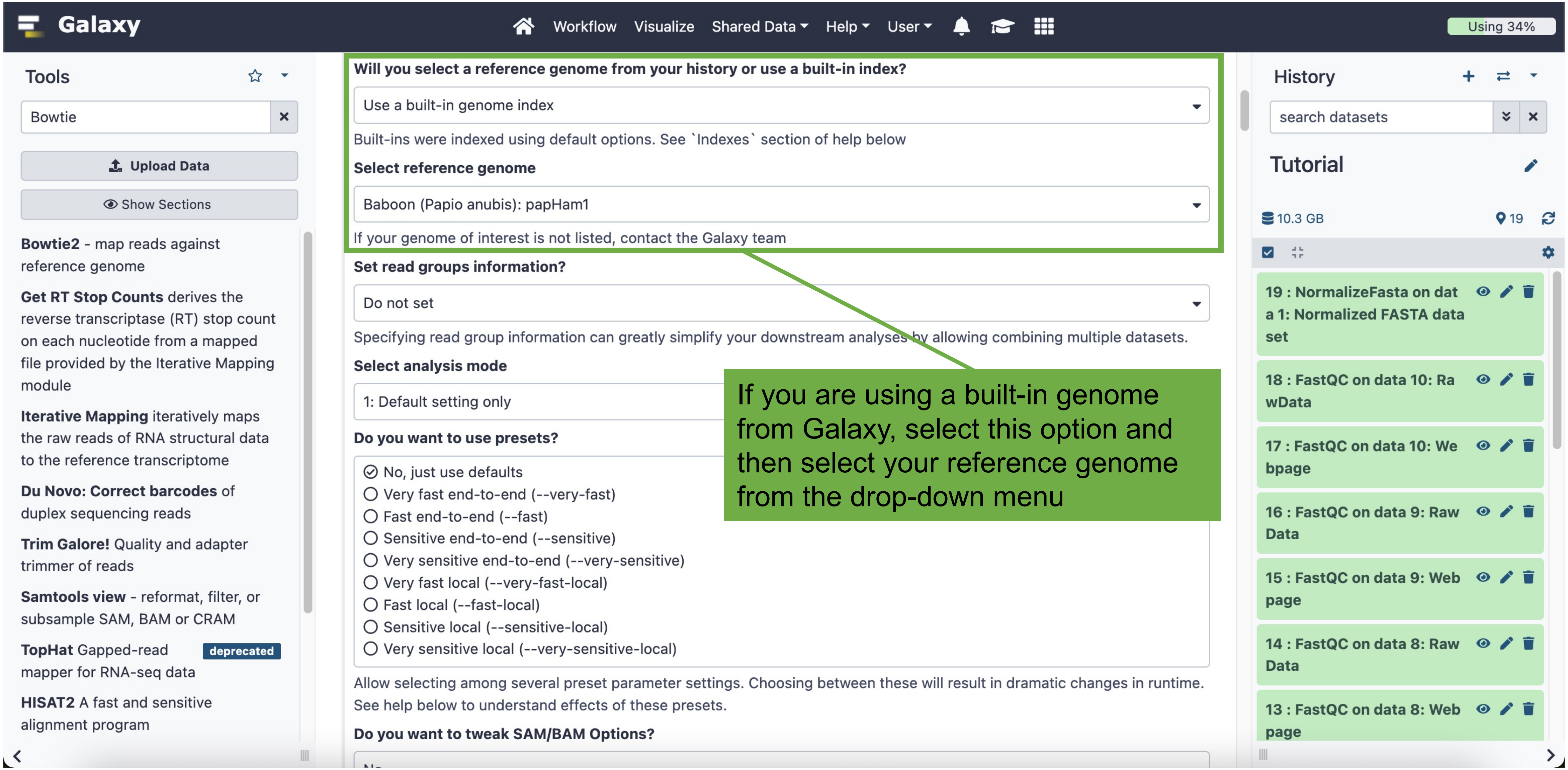
Galaxy
Workflow
Visualize
Shared Data
Help
User
Using 34%
Tools
Bowtie
Upload Data
Show Sections
Bowtie2 - map reads against reference genome
Get RT Stop Counts derives the reverse transcriptase (RT) stop count on each nucleotide from a mapped file provided by the Iterative Mapping module
Iterative Mapping iteratively maps the raw reads of RNA structural data to the reference transcriptome
Du Novo: Correct barcodes of duplex sequencing reads
Trim Galore! Quality and adapter trimmer of reads
Samtools view - reformat, filter, or subsample SAM, BAM or CRAM
TopHat Gapped-read mapper for RNA-seq data
deprecated
HISAT2 A fast and sensitive alignment program
Will you select a reference genome from your history or use a built-in index?
Use a built-in genome index
Built-ins were indexed using default options. See `Indexes` section of help below
Select reference genome
Baboon (Papio anubis): papHam1
If your genome of interest is not listed, contact the Galaxy team
Set read groups information?
Do not set
Specifying read group information can greatly simplify your downstream analyses by allowing combining multiple datasets.
Select analysis mode
1: Default setting only
Do you want to use presets?
No, just use defaults
Very fast end-to-end (--very-fast)
Fast end-to-end (--fast)
Sensitive end-to-end (--sensitive)
Very sensitive end-to-end (--very-sensitive)
Very fast local (--very-fast-local)
Fast local (--fast-local)
Sensitive local (--sensitive-local)
Very sensitive local (--very-sensitive-local)
Allow selecting among several preset parameter settings. Choosing between these will result in dramatic changes in runtime. See help below to understand effects of these presets.
Do you want to tweak SAM/BAM Options?
If you are using a built-in genome from Galaxy, select this option and then select your reference genome from the drop-down menu
History
search datasets
Tutorial
10.3 GB
19
19 : NormalizeFasta on data 1: Normalized FASTA data set
18 : FastQC on data 10: RawData
17 : FastQC on data 10: Webpage
16 : FastQC on data 9: Raw Data
15 : FastQC on data 9: Web page
14 : FastQC on data 8: Raw Data
13 : FastQC on data 8: Web page

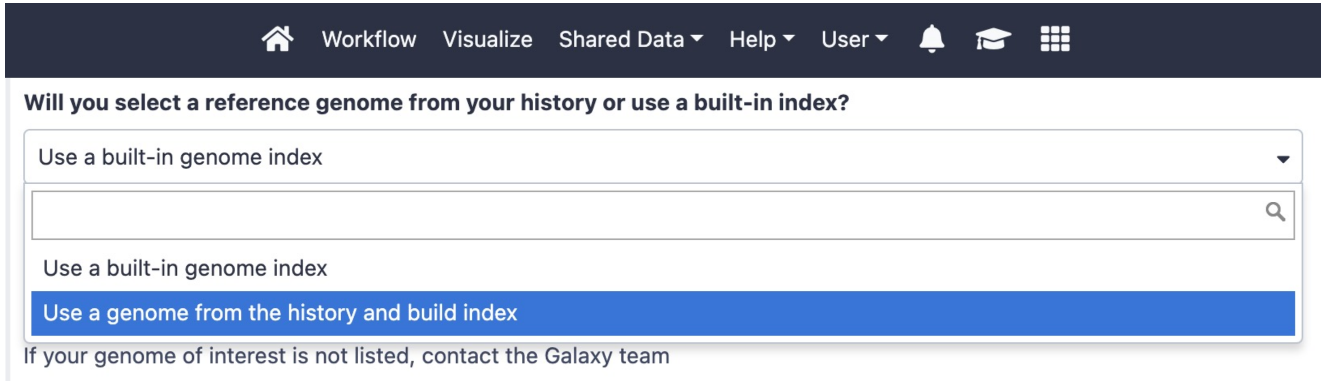


If you are using a custom genome, use the drop-down button to select that option.

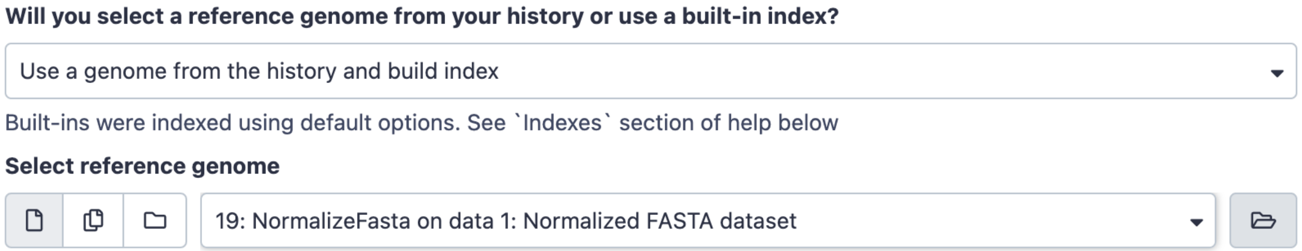


Be sure to select the normalized custom genome as your reference genome.

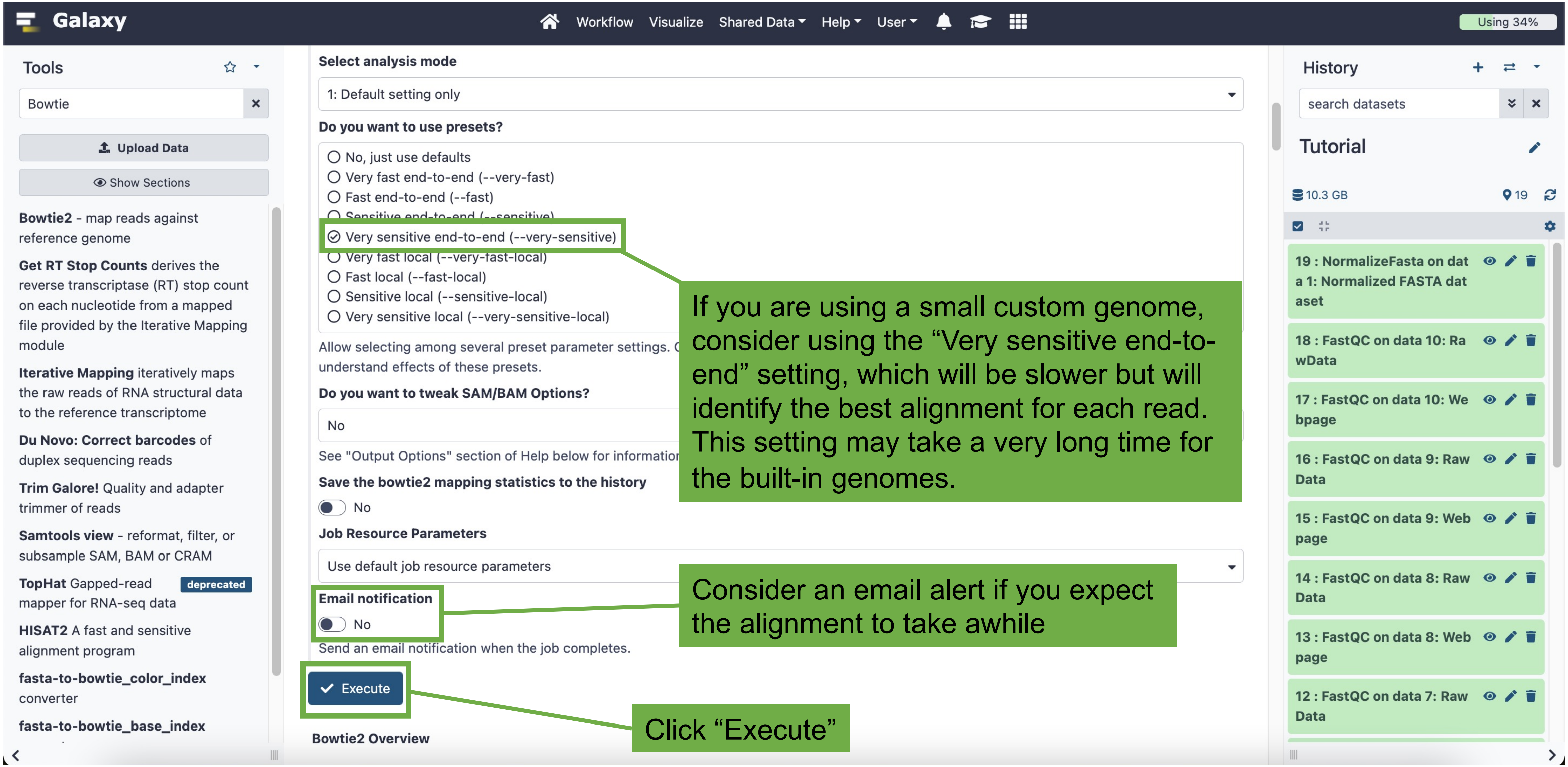
Galaxy
Workflow
Visualize
Shared Data
Help
User
Using 34%
Tools
Bowtie
Upload Data
Show Sections
Bowtie2 - map reads against reference genome
Get RT Stop Counts derives the reverse transcriptase (RT) stop count on each nucleotide from a mapped file provided by the Iterative Mapping module
Iterative Mapping iteratively maps the raw reads of RNA structural data to the reference transcriptome
Du Novo: Correct barcodes of duplex sequencing reads
Trim Galore! Quality and adapter trimmer of reads
Samtools view - reformat, filter, or subsample SAM, BAM or CRAM
TopHat Gapped-read mapper for RNA-seq data
deprecated
HISAT2 A fast and sensitive alignment program
fasta-to-bowtie_color_index converter
fasta-to-bowtie_base_index
Select analysis mode
1: Default setting only
Do you want to use presets?
No, just use defaults
Very fast end-to-end (--very-fast)
Fast end-to-end (--fast)
Sensitive end-to-end (--sensitive)
Very sensitive end-to-end (--very-sensitive)
Very fast local (--very-fast-local)
Fast local (--fast-local)
Sensitive local (--sensitive-local)
Very sensitive local (--very-sensitive-local)
Allow selecting among several preset parameter settings.
understand effects of these presets.
Do you want to tweak SAM/BAM Options?
No
See "Output Options" section of Help below for information
Save the bowtie2 mapping statistics to the history
No
Job Resource Parameters
Use default job resource parameters
Email notification
No
Send an email notification when the job completes.
Execute
Bowtie2 Overview
If you are using a small custom genome, consider using the “Very sensitive end-to-end” setting, which will be slower but will identify the best alignment for each read. This setting may take a very long time for the built-in genomes.
Consider an email alert if you expect the alignment to take awhile
Click “Execute”
History
search datasets
Tutorial
10.3 GB
19
19 : NormalizeFasta on data 1: Normalized FASTA dataset
18 : FastQC on data 10: RawData
17 : FastQC on data 10: Webpage
16 : FastQC on data 9: Raw Data
15 : FastQC on data 9: Webpage
14 : FastQC on data 8: Raw Data
13 : FastQC on data 8: Webpage
12 : FastQC on data 7: Raw Data

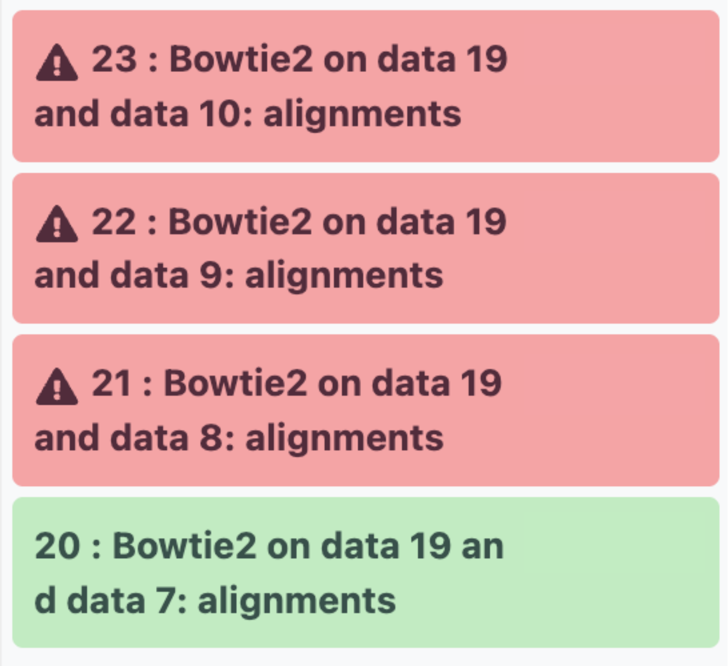


A quick note that even if everything is set up correctly, sometimes the jobs can fail due to Galaxy issues.

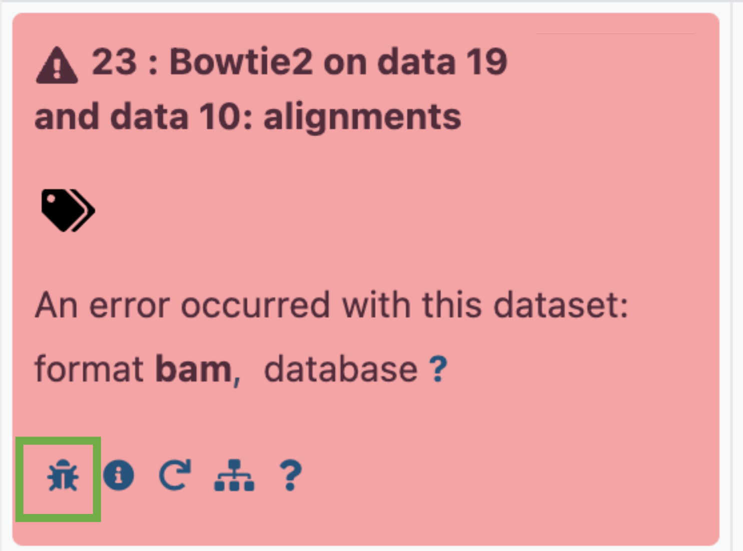


To diagnose a job failure, click on the dataset name to open, then click on the bug icon. A report will open in the main window with an error code, which you can use to search. These job failures were due to a Galaxy issue.

Because these jobs failed, I deleted them from my history (using the trash can icon). Note that even if you delete jobs, the numbering system that Galaxy uses will continue linearly. I deleted jobs 20-23, but when I repeated Bowtie, the successful runs were numbered 24-27.

Bowtie2 produces .bam files, which are large and can be difficult to view. We convert the .bam files to .bigwig files, which are much smaller and user-friendly.

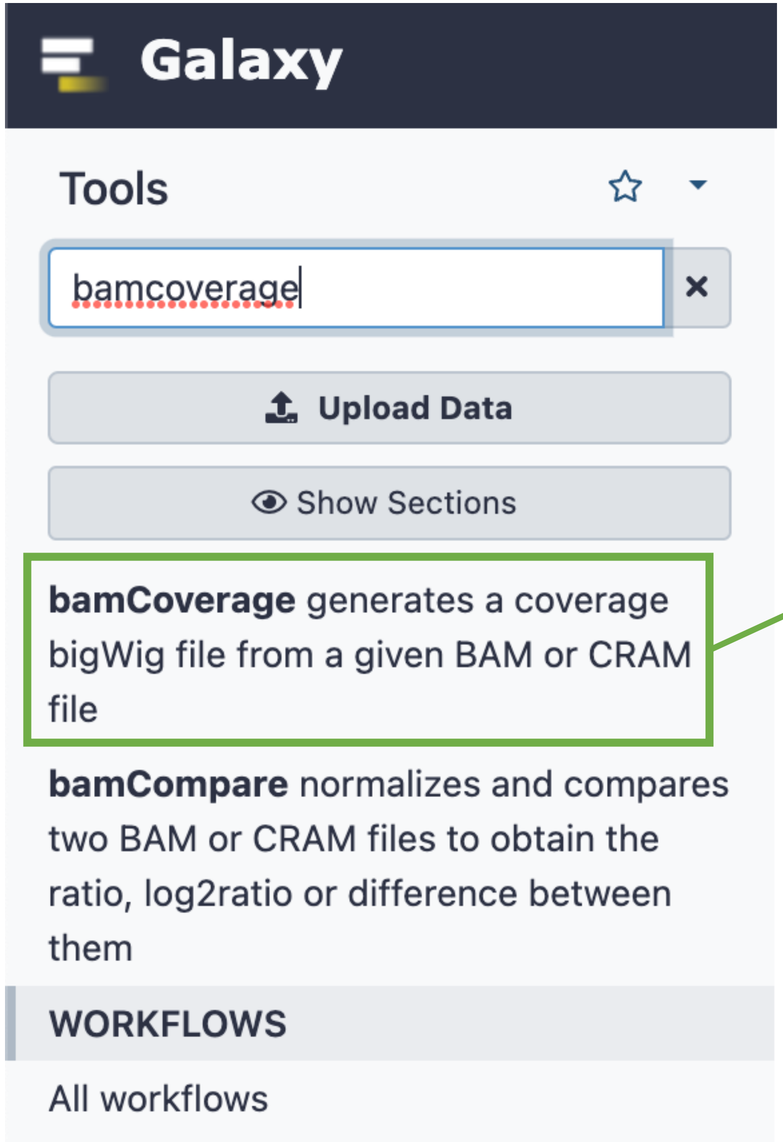


Search for “bamCoverage”, then click on the “bamCoverage” tool

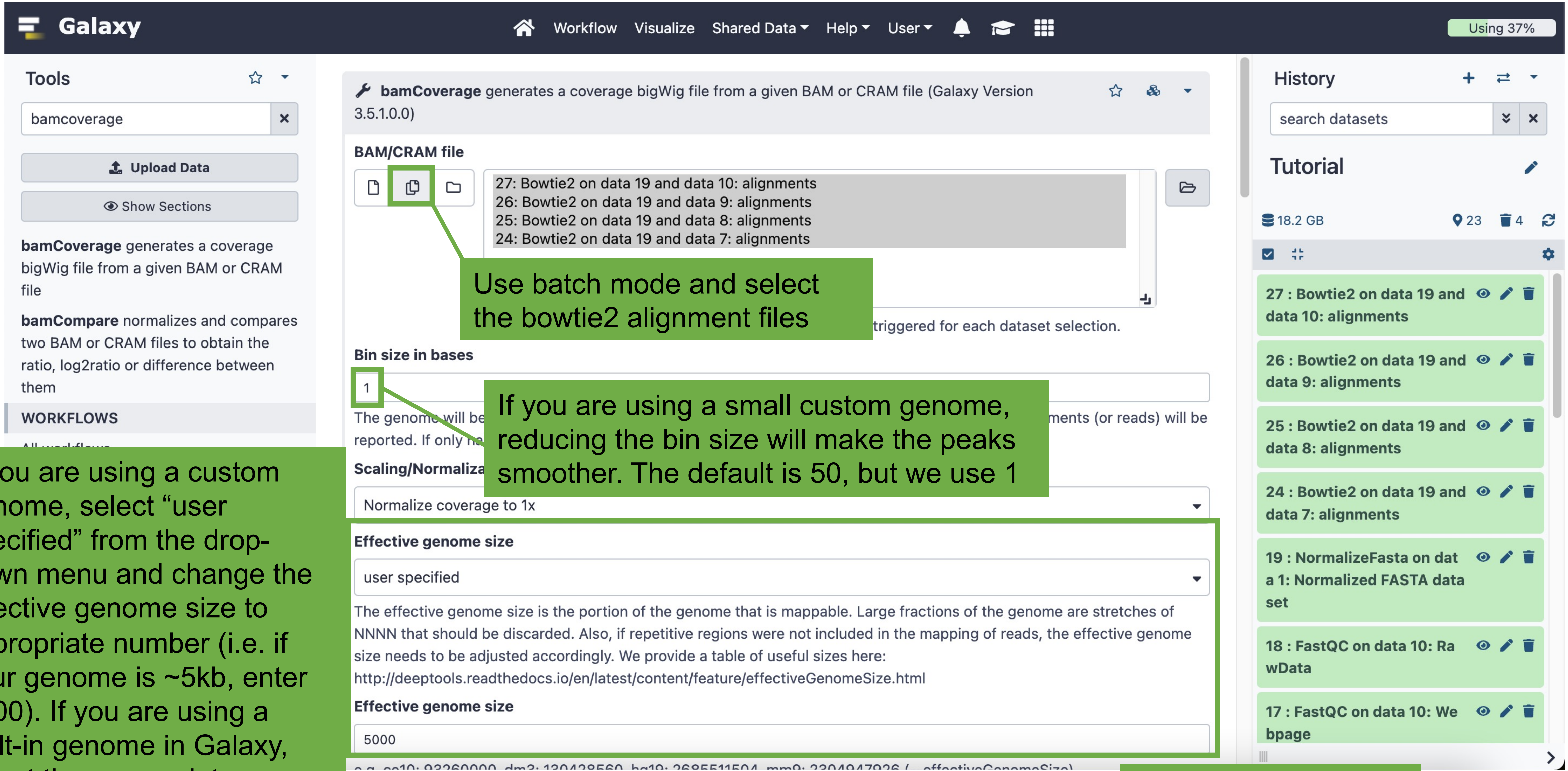


Use batch mode and select the bowtie2 alignment files

If you are using a small custom genome, reducing the bin size will make the peaks smoother. The default is 50, but we use 1

If you are using a custom genome, select “user specified” from the drop-down menu and change the effective genome size to appropriate number (i.e. if your genome is ~5kb, enter 5000). If you are using a built-in genome in Galaxy, select the appropriate organism.

Click “Execute” at the bottom of the page

If your dataset contains “input” or “IgG” conditions, the ChIP runs can be normalized. The bamCompare tool uses the .bam files from the control and ChIP runs, and produces a .bigwig file.

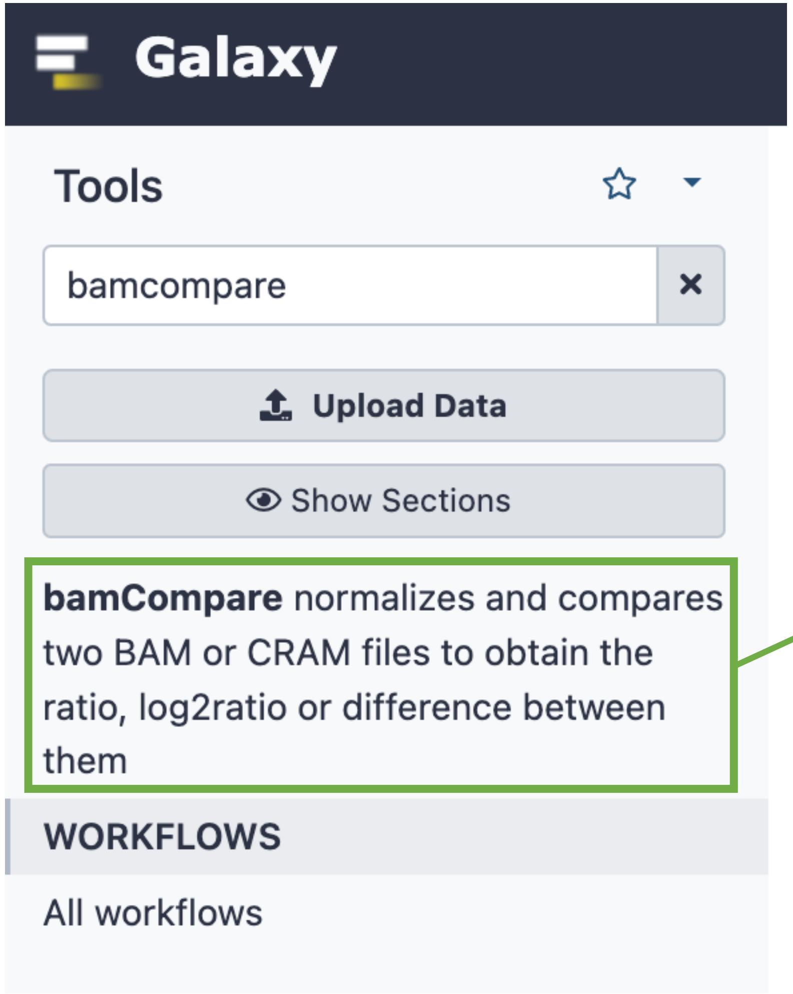


Search for “bamCompare”, then click on the “bamCompare” tool

We recommend setting up the bamCompare jobs one at a time rather than in batch mode to ensure that the correct runs are paired.

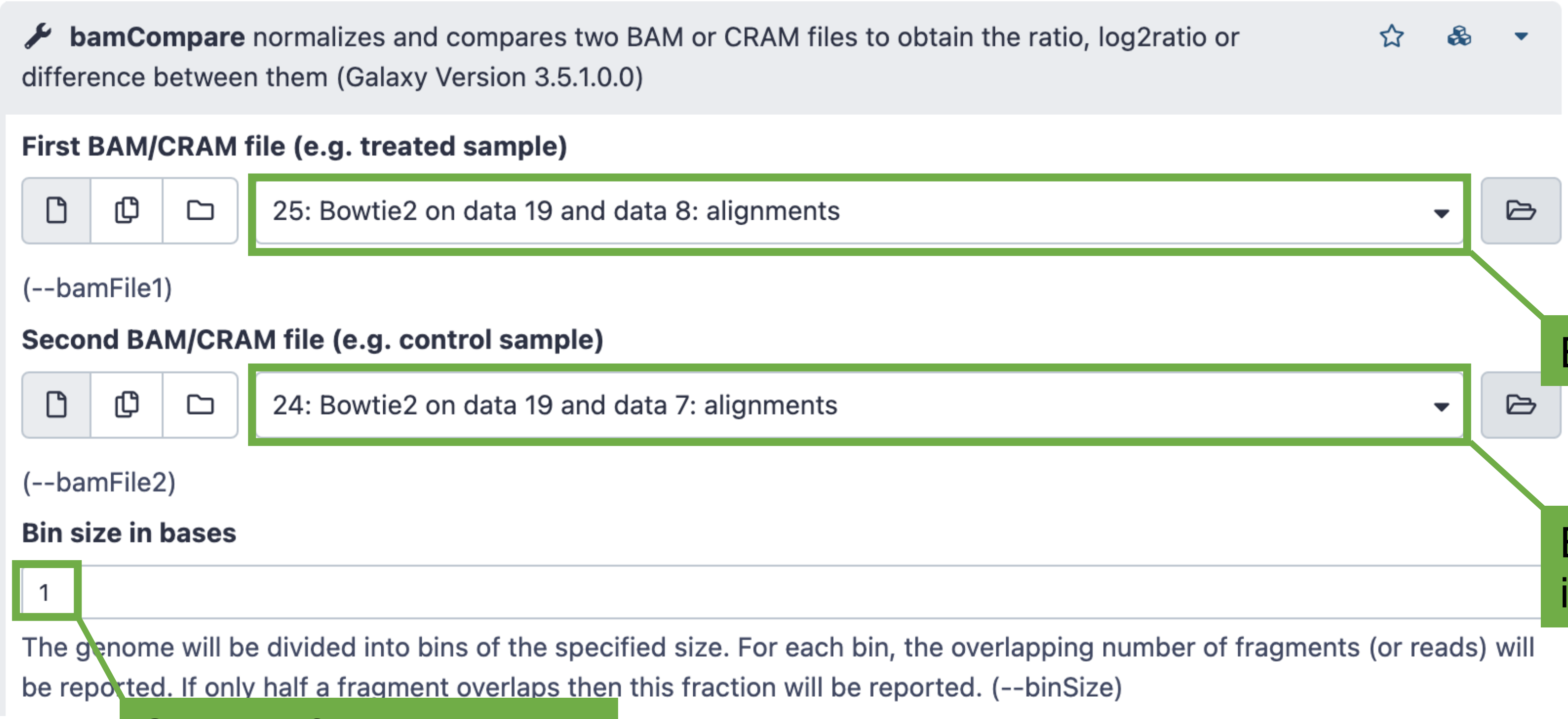


Enter a ChIP run here

Enter the corresponding input/IgG run here

Change if your custom genome is small (default is 50, but we use 1)

Click “Execute” at the bottom of the page

When the bamCoverage and bamCompare jobs are complete, you're done in Galaxy. Download the .bigwig files by clicking on the eye icon for each.

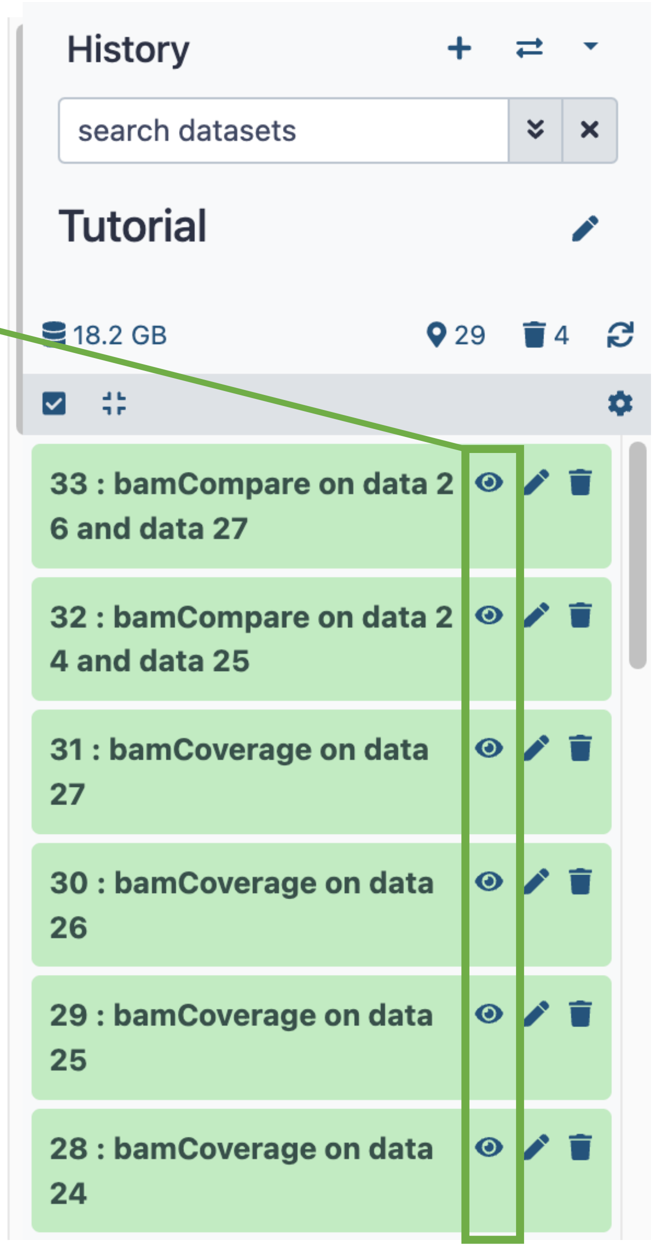

### Visualizing in IGV

Download IGV software:
https://software.broadinstitute.org/software/igv/download

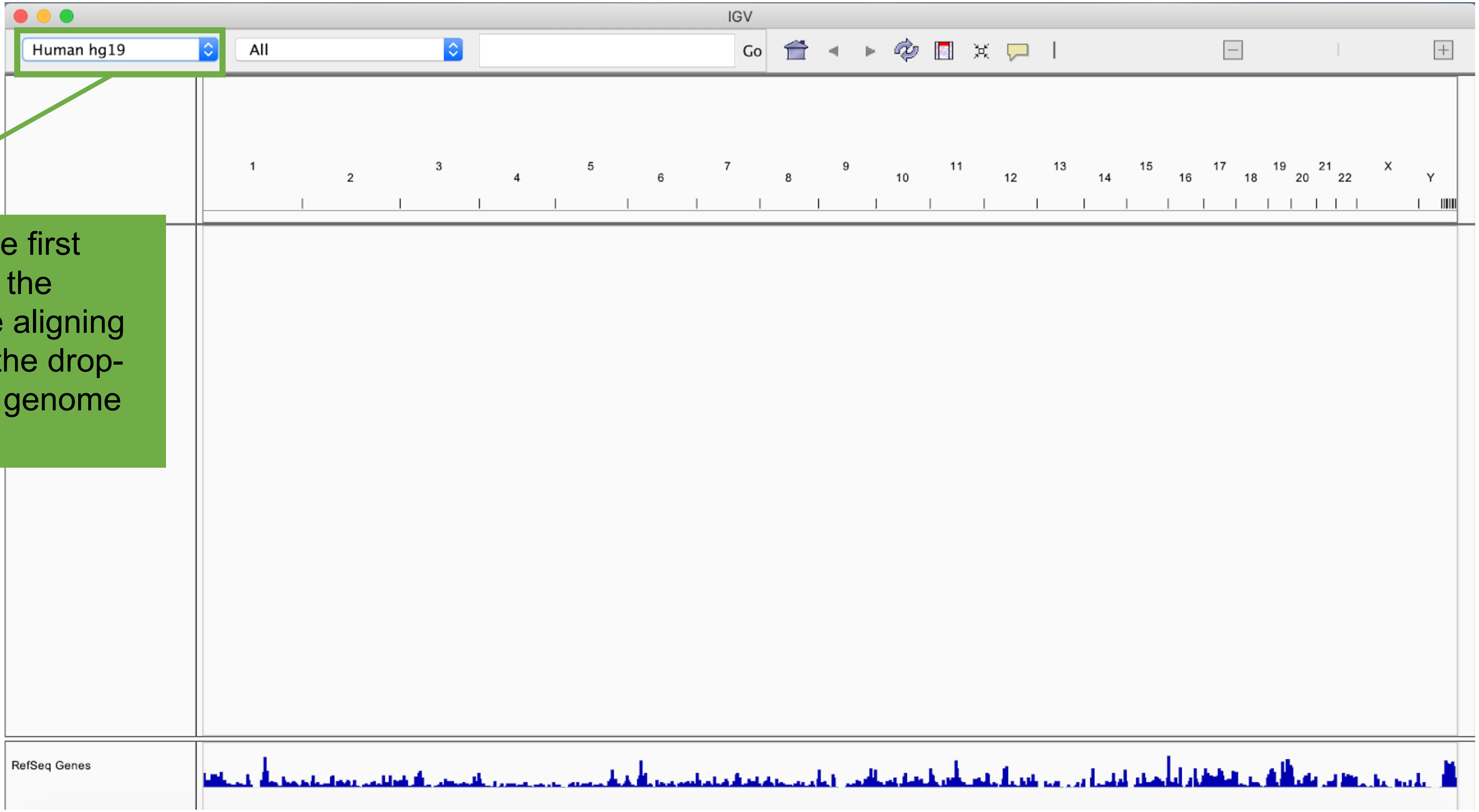


When you open IGV for the first time, it will likely default to the human genome. If you are aligning to an entire genome, use the drop-down menu to select your genome of interest.

Load the genome file (.fa)

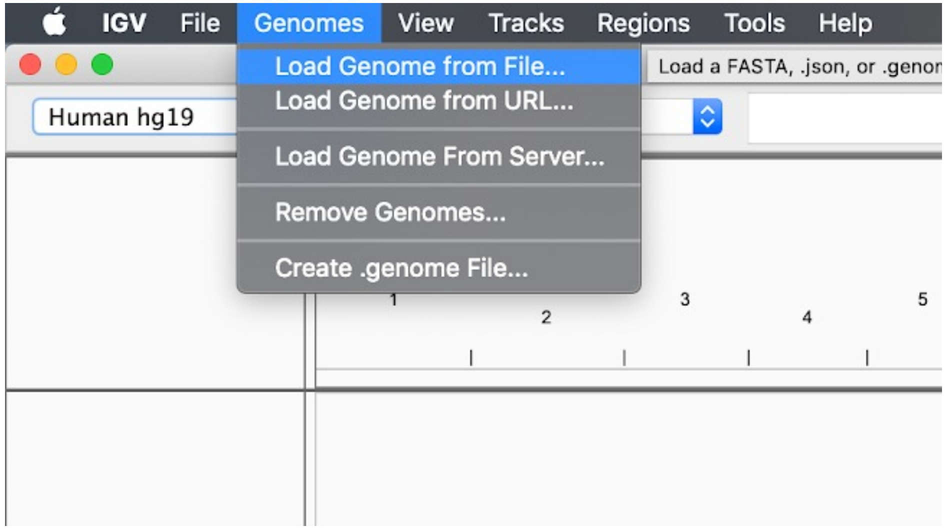


Load the annotation file (.bed)

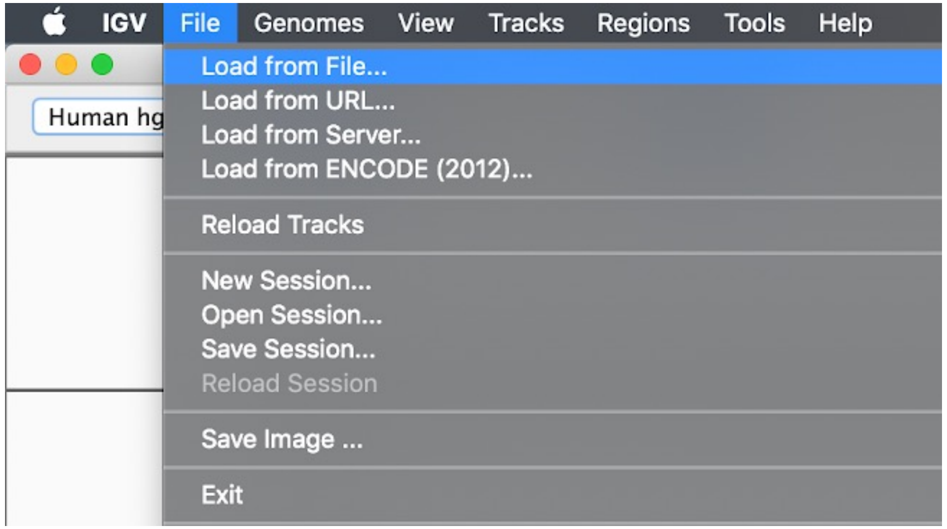


If you are using a custom genome, you will need both the genome file (.fasta or .fa) and an annotation file (.bed). The genome file must be the same as the one you used in Galaxy. The .bed file is not strictly necessary, but it annotates the genome file with features such as genes or transcription factor binding sites.

If you are using a built-in genome, skip this step.

Next: load .bigwig files

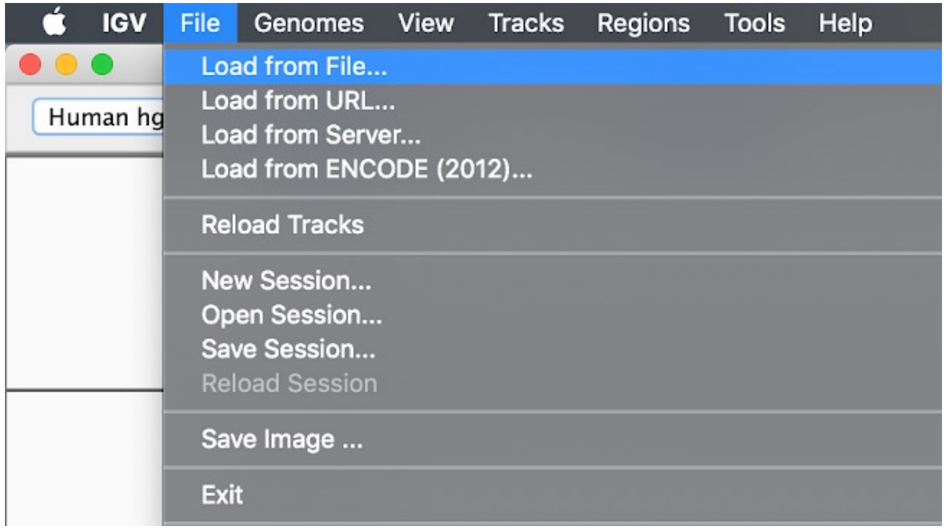


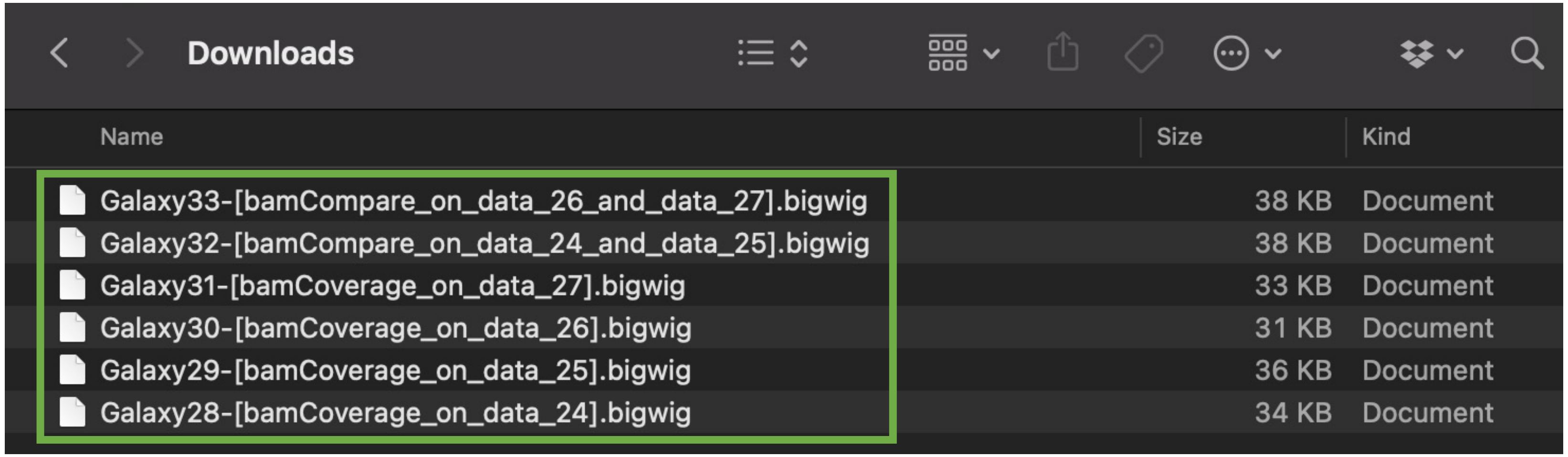

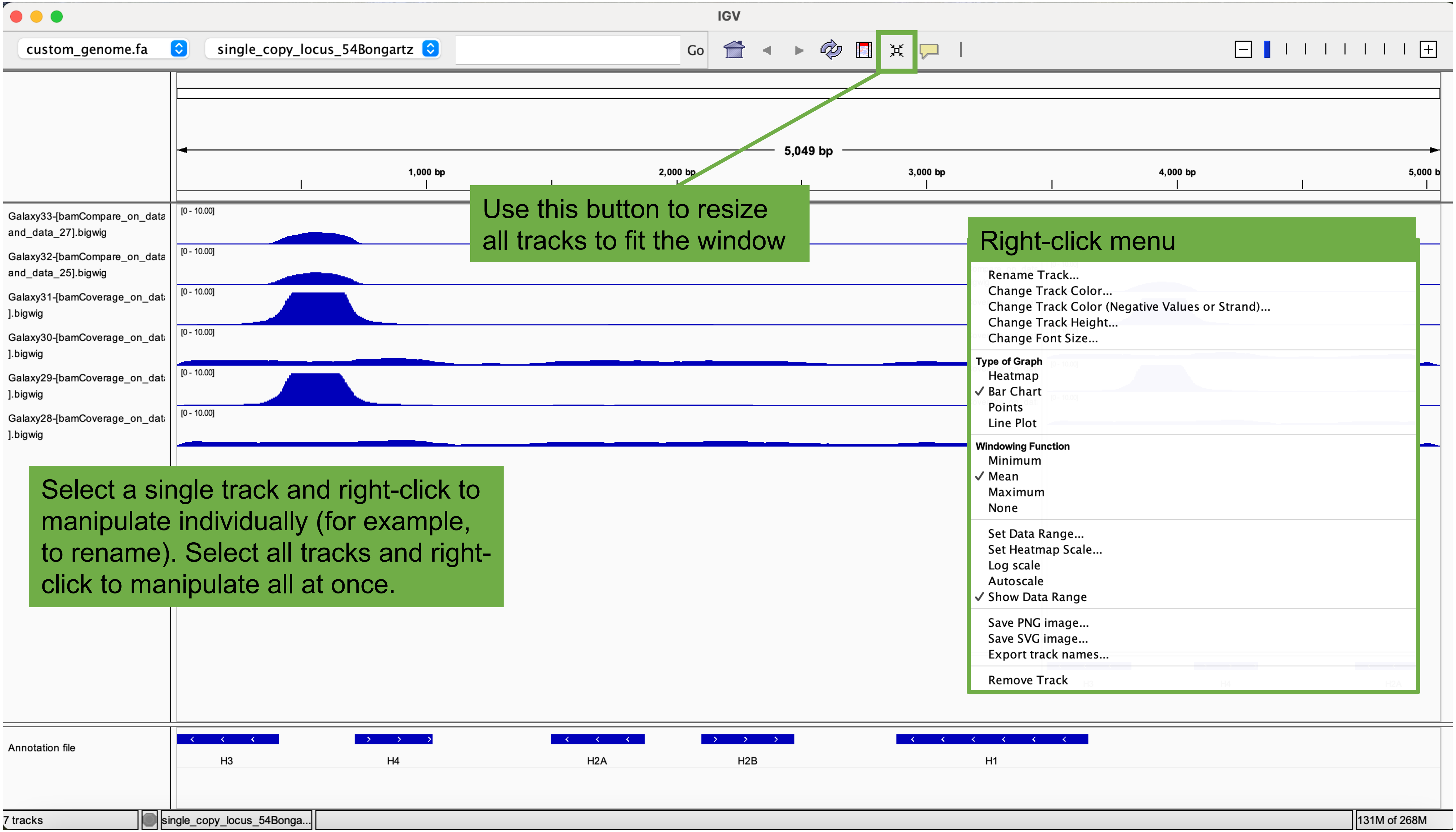
IGV
custom_genome.fa
single_copy_locus_54Bongartz
Go
5,049 bp
1,000 bp
2,000 bp
3,000 bp
4,000 bp
Use this button to resize all tracks to fit the window
Galaxy33-[bamCompare_on_data and_data_27].bigwig
Galaxy32-[bamCompare_on_data and_data_25].bigwig
Galaxy31-[bamCoverage_on_dat ].bigwig
Galaxy30-[bamCoverage_on_dat ].bigwig
Galaxy29-[bamCoverage_on_dat ].bigwig
Galaxy28-[bamCoverage_on_dat ].bigwig
[0 - 10.00]
Right-click menu
Rename Track...
Change Track Color...
Change Track Color (Negative Values or Strand)...
Change Track Height...
Change Font Size...
Type of Graph
Heatmap
✓ Bar Chart
Points
Line Plot
Windowing Function
Minimum
✓ Mean
Maximum
None
Set Data Range...
Set Heatmap Scale...
Log scale
Autoscale
✓ Show Data Range
Save PNG image...
Save SVG image...
Export track names...
Remove Track
Select a single track and right-click to manipulate individually (for example, to rename). Select all tracks and right-click to manipulate all at once.
Annotation file
H3
H4
H2A
H2B
H1
7 tracks
single_copy_locus_54Bonga...
131M of 268M

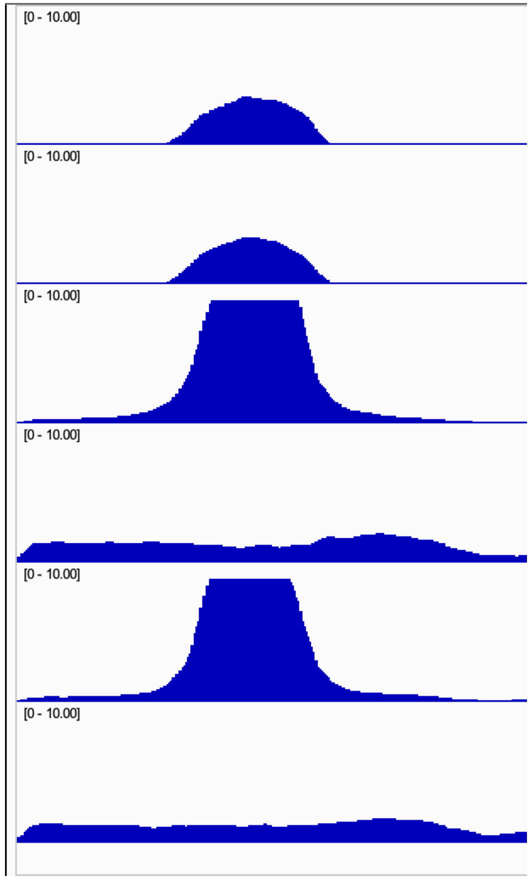


Note here that the scales are the same and some of the tracks look like they are cut off. Highlight all tracks, right click, and select “autoscale” to put each track in its own full data range. Select “group autoscale” to put all tracks in the same data range.

Autoscale

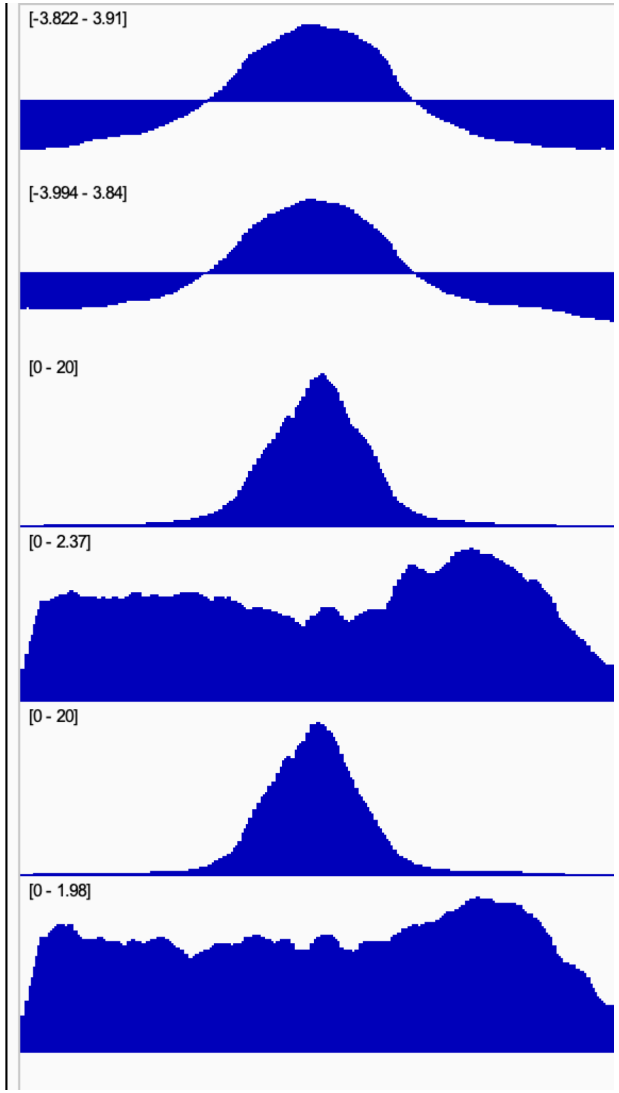


Group autoscale

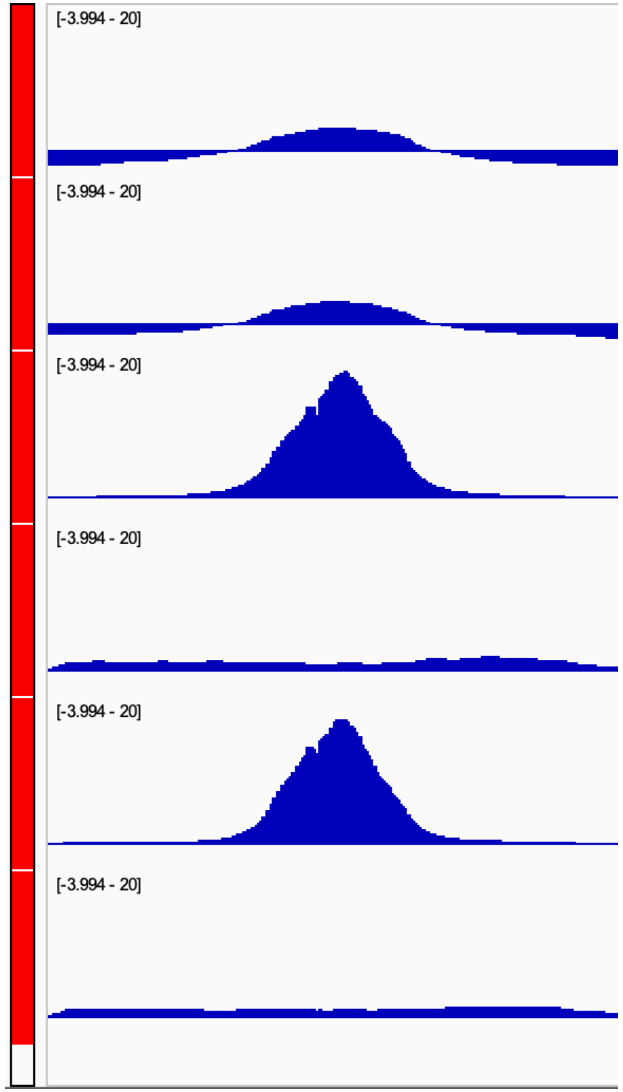

Other recommendations and ideas for IGV:

- Re-order tracks by clicking and dragging
- Rename tracks with descriptive identifier (right click track > Rename Track)
- Change track color (right click track > Change Track Color)
- Change graph type (right click track > type of graph (choose heatmap, bar chart, points, or line plot)
- Overlay two replicate tracks (select two tracks > right click > Overlay Tracks)
- Save a screenshot (File > Save PNG Image)
- Save session for later (File > Save Session)

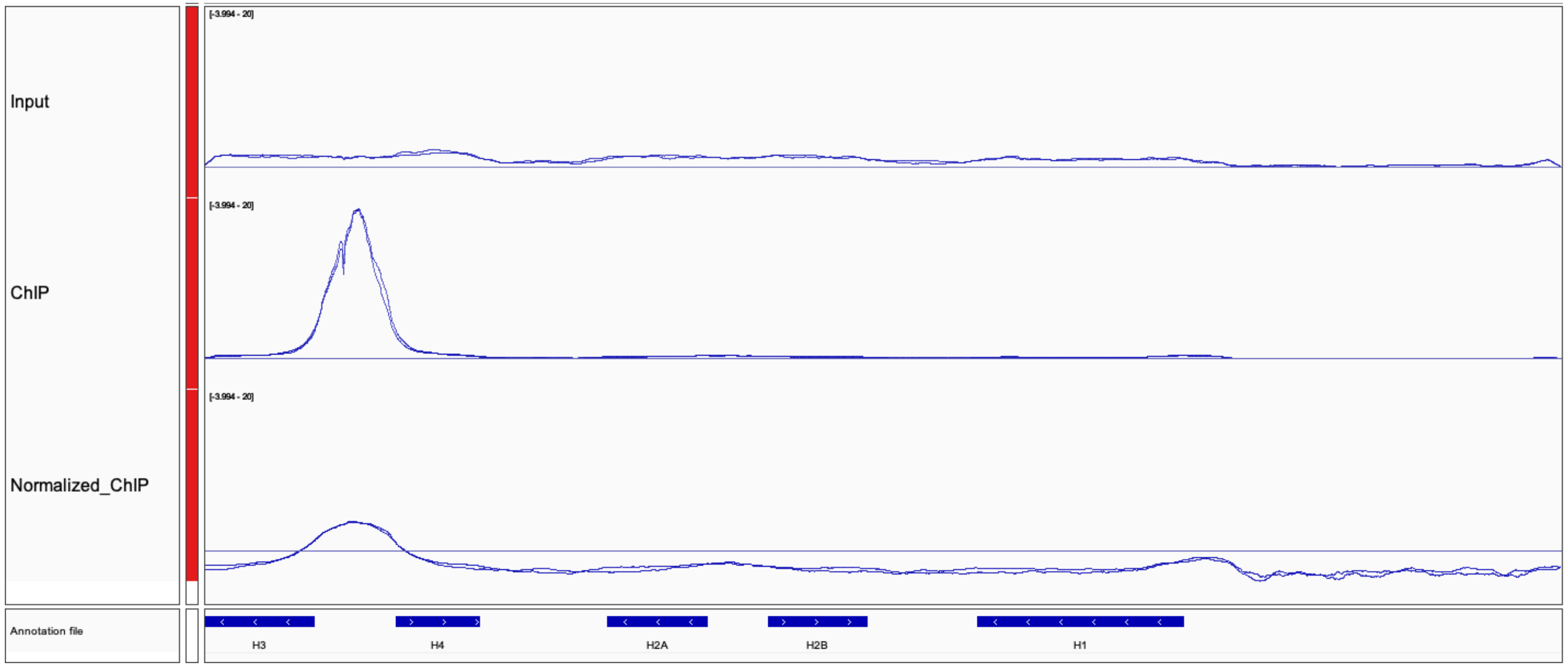


Example screenshot with replicates overlaid and changed to line plot. The font size of the tracks is increased to 16. The tracks are group autoscaled.

The best way to get familiar with IGV is to load tracks and try out different settings!

### Some tips and tricks

You can rename a file in Galaxy, but when you perform a job, the name won't carry over. We recommend using a table to keep track of everything. Here is an example:

| **File** | **Galaxy import** | **FastQC webpage** | **FastQC Raw** | **Bowtie2** | **BamCoverage** | **BamCompare** |
|---|---|---|---|---|---|---|
| Input_rep1 | 7 | 11 | 12 | 24 | 28 | 33 |
| ChIP_rep1 | 8 | 13 | 14 | 25 | 29 | 33 |
| Input_rep2 | 9 | 15 | 16 | 26 | 30 | 32 |
| ChIP_rep2 | 10 | 17 | 18 | 27 | 31 | 32 |

Once you are finished in Galaxy, you can re-name everything in IGV.
